# A common venomous ancestor? Prevalent bee venom genes evolved before the aculeate stinger while few major toxins are bee-specific

**DOI:** 10.1101/2022.01.21.477203

**Authors:** Ivan Koludarov, Mariana Velasque, Thomas Timm, Carola Greve, Alexander Ben Hamadou, Deepak Kumar Gupta, Günter Lochnit, Michael Heinzinger, Andreas Vilcinskas, Rosalyn Gloag, Brock A. Harpur, Lars Podsiadlowski, Burkhard Rost, Timothy N. W. Jackson, Sebastien Dutertre, Eckart Stolle, Björn M von Reumont

## Abstract

Venoms, which have evolved numerous times in animals, are ideal models of convergent trait evolution. However, detailed genomic studies of toxin-encoding genes exist for only a few animal groups. The hyper-diverse hymenopteran insects are the most speciose venomous clade, but investigation of the origin of their venom genes has been largely neglected. Utilising a combination of genomic and proteo-transcriptomic data, we investigated the origin of 11 toxin genes in 29 published and three new hymenopteran genomes and compiled an up-to-date list of prevalent bee venom proteins. Observed patterns indicate that bee venom genes predominantly originate through single gene co-option with gene duplication contributing to subsequent diversification. Most Hymenoptera venom genes are shared by all members of the clade and only melittin and the new venom protein family anthophilin1 appear unique to the bee lineage. Most venom proteins thus predate the mega-radiation of hymenopterans and the evolution of the aculeate stinger.

## Introduction

Molecular processes involved in the evolution of adaptive traits are among the most widely discussed topics in biology^1,2^. Venoms are complex secretory mixtures that are injected into other organisms for predation, defence or competition, using a specialized morphological structure known as the venom apparatus. Venom toxins – the molecules associated with the venomous function – are typically short peptides, enzymes and other proteins^3^. Because the function of many toxin-encoding genes is relatively free from pleiotropic and epistatic complications – one gene typically encodes one toxin with a clear functional role – toxins provide an excellent opportunity for investigation of the molecular mechanisms that facilitate the evolution of adaptive traits. Advances in comparative genomics and sequencing are furthering our efforts to understand these mechanisms at the genomic level^4–7^. Nevertheless, there have been only few large comparative studies focusing on the genomic origins of toxin genes and their weaponization, mostly in snakes and only few other clades such as cnidarians^2,7–11^. One reason for this is the predominant interest in venoms for their pharmacological and agrochemical applications or clinical toxinology^12,13^. Researchers have therefore prioritized groups such as snakes, scorpions, spiders, and cone snails that may not be species-rich compared to insects but are known for powerful venom components of which many cause strong envenomation effects on humans^4,12,13^. This taxonomic bias hinders a deeper understanding of the origins and evolution of venoms, and leaves the vast body of knowledge hidden in the mega-radiations of insects relatively untapped.

Hymenopterans (sawflies, parasitoid wasps, true wasps, ants and bees) are the most species-rich insect group and are of tremendous ecological and economical importance^14^. However, they also feature the largest number of venomous species. Furthermore, their venom delivery system exists in a variety of states within the order, from its origin as an ovipositor that co-injected immunomodulatory “venom” along with eggs into plant hosts (as in extant Symphyta), to the high-pressure venom systems of majority of wasps and bees, to secondary losses in both bee and ant lineages^15,16^. Like snakes, therefore, Hymenoptera provide an opportunity to investigate the co-evolution of toxin genes and associated anatomy within a larger clade. Unsurprisingly, given their economic significance, honeybee and bumblebee venoms have received the lion’s share of toxinological attention, and are among the best-characterized venoms in the animal kingdom^16,17^. The venoms of the remaining species of the hymenopteran radiation, however, including the majority of bees, remain largely unexplored despite recent proteo-transcriptomic studies on several ant and wasp species^18–21^. Where studies of lesser-known Hymenoptera have been conducted, they typically deal with single crude fractions or even individual components either due to technical limitations at the time or because of applied research focus^16,22,23^. In general, proteo-transcriptomic studies focused on injected and functionally described components are rather sparse and often focus on small peptides and/or are available for only few smaller groups or single taxa of hymenopterans, such as honey bees^24^, ants^25,26^, spider wasps^27^, and true wasps^28^. Only the recent study by Robinson et al.^26^ proposed that short toxin peptides of ants, bees and wasps compose a family of aculeatoxins based on the similarity of aligned propeptides sequences, however, a detailed phylogenetic analysis is not provided, see **Figure 1**.

**Figure 1:**
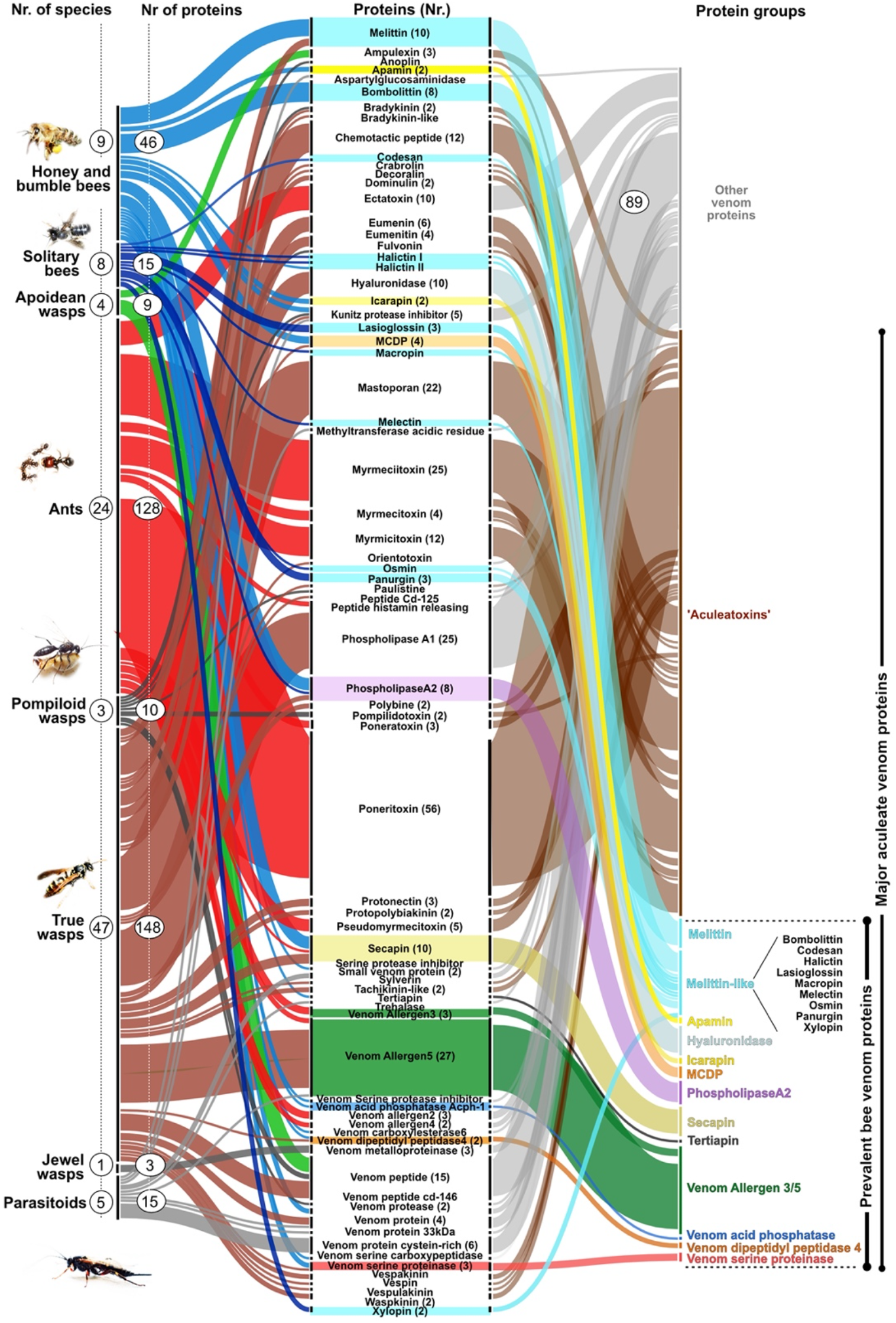
Reviewed venom proteins for hymenopteran taxa in respect to protein and species numbers from UniProt. Major hymenopteran clades are shown on the left (species numbers in circles). The second numbers in circles within the colour-coded lines indicate venom proteins (Grouped according to their names). The twelve herein proposed prevalent bee venom protein families (PBVP) are illustrated on the right, together with the toxins proposed as ‘Aculeatoxins’ (brown) according to Robinson et al.^26^. Novel, and further undescribed peptides and proteins are shown in grey. The hymenopteran groups are based on the recent phylogeny according to Peters et al.^31^.

Our study represents the first taxon-wide comparative genomics analysis of bee venoms. We address two key questions: (1) whether bee venoms are predominantly comprised of toxins that are novel and unique to this clade, and (2) whether single gene co-option is the major mechanism of venom gene evolution in bees, as is the case for parasitoid wasps. We then utilise the insights generated to conjecture as to whether or not ecological and anatomical adaptations are reflected in the patterns of venom gene evolution. Throughout the paper, we distinguish between “venom proteins” (or the genes that encode them) and “toxins” (or toxin-encoding genes). The former are those proteins *associated* with the venom system (often secreted in the venom itself) but not necessarily having toxic functions themselves – we reserve the designation “toxin” for those gene products with *characterised toxic functions within venom*. Given a permissive definition of the label “venomous” (see discussion), our results suggest that the entire extant Hymenoptera lineage may be descended from a “common venomous ancestor”, and indeed that the argument for this may be stronger than the similar argument made for the squamate lineage Toxicofera^29^.

## Results

### The most prevalent bee venom proteins and their genomic framework

We establish here a set of 12 proteins that we identify as the most prevalent injected bee venom components based on mining of published sequences, data of toxins with known activity^18,24,30^ (see **Figure 1**), and own proteo-transcriptome data.

New venom profiles were generated for two phylogenetically distant solitary bees, the great-banded furrow-bee (*Halictus scabiosae*), the violet carpenter bee (*Xylocpopa violacea*) and the honeybee (*A. mellifera*) as complementary data (**Figure 2** and **Supplementary Tables 1-3**). All three venoms predominantly contained low-molecular-weight peptides, in particular melittin, apamin and mast-cell degranulating peptide (MCDP). Larger proteins such as phospholipase A2, venom acid phosphatase, venom dipeptidyl peptidase 4 and venom allergens made up less than 10% of the transcripts based on expression values (see **Figure 2 and Material and Methods**).

**Figure 2:**
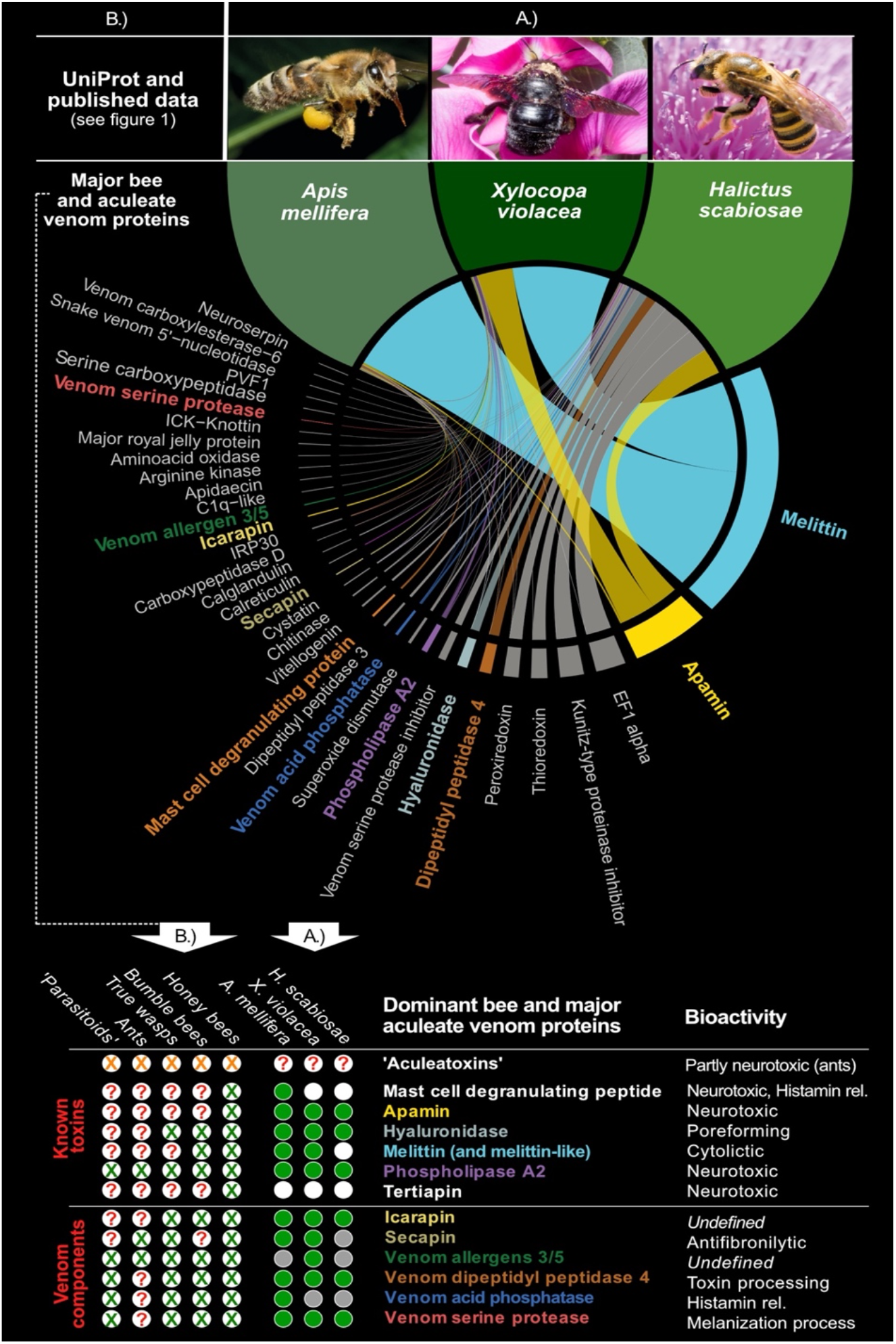
The most prevalent bee venom proteins. Components selected from our own data (A.) *A. mellifera, H. scabiosae* and *X. violacea* profiles, and (B.) published bee and aculeate venom components. In A.) only venom protein transcripts validated by the proteome data are listed. Transcript expression is shown as thickness of the circus plot lines and based on the percentage of scaled transcript per million (TPM) values including only proteome-validated sequences. The twelve selected venom proteins that we discuss herein further as dominant bee venom proteins are printed in bold in the colour code used for these proteins in this manuscript. Peptide names in white were not identified by our proteo-transcriptome data but are present in published data. For our new proteo-transcriptome data (A.) the green circles indicate venom proteins identified by proteo-transcriptomics, grey circles indicate transcriptome-only hits. White circles illustrate missing data. For published data the green X indicate major components identified in literature, red questions marks highlight missing/unclear data. Orange X highlight the ‘aculeatoxin ‘peptides (According to Robison et al.^26^ melittin is also a member of the proposed aculeatoxin family, which is separately shown as part of the PBVPs).

We have to state critically that the heterogeneous picture of venom expression (Figure 2A) could be reasoned by the difficulty to synchronize the physiological state of venom glands, especially for solitary bees. The venom compositions and species-specific differences (especially for *H. scabiosa*) will be discussed in-depth elsewhere (see von Reumont etal. 2022 for *X. violacea*^21^). In general, the new profiles corroborate our selection of prevalent bee venom proteins (Figure 2). Further analysis is restricted to these 12, which include toxins and six auxiliary venom peptide and protein families mostly with known function, but also including two prevalent venom protein families of currently unknown function (Venom allergen 3/5 and Icarapin), see **Supplementary Table 4**. We refer to these venom components from here on as prevalent bee venom proteins (PBVP).

Two major groups are distinguishable in the PBVP – toxins with characterised acutely toxic functions such as neurotoxicity (e.g. Apamin) or cytotoxicity (e.g. Mellitin), and proteins consistently present in the crude venom presumably as accessory components (**Figure 2**). To uncover the evolutionary history of the prevalent bee venom proteins, we analysed corresponding genomic regions by searching for homologs in 29 published genomes (see **Supplementary Table 5**) of bees and outgroups (sawflies, jewel wasp, ants, paper wasps) and our three genomes of two sweat bees and the violet carpenter bee (See material and methods for further details and **Supplementary Table 6**). The selected taxa span 300 million years of evolution and include representatives of the phytophagous sawflies (Symphyta), the most basal hymenopteran group. We used the well-annotated *A. mellifera* reference genome to trace venom genes and their flanking genes based on exon regions. We identified orthologs for each exon in other genomes, which were collected into an extended database. We searched all genomes using this database and the manually inspected the results to establish completeness and microsynteny, which reflects the arrangement and position of flanking exons of genes around venom protein genes (see details in material and methods). “Synteny” refers to shared patterns of gene arrangement (“colinearity”) in homologous genomic regions across taxa. When sufficiently high-quality genomic sequences are available and genes of interest are located in stable regions, the ability to utilise microsyntenic analyses – comparisons of synteny/colinearity in short stretches of the genome – is a key advantage of comparative genomics. Where sequencing is sufficiently contiguous, these analyses reveal the arrangement of genes and their neighbours as physically instantiated in a chromosomal region. By mapping such regions (see, e.g., **Figures 3 and 4**), including genes of interest and their neighbours, it is possible to catalogue rearrangements that occur in diverse taxa. Put simply, observation of the spatial relations between genes of interest and their neighbours (both complete genes and gene fragments) in one species, enables identification of homologous genes in additional taxa by examination of the sequences that flank these genes. Attention to “genomic context”, therefore, enables a clearer identification of orthologs than phylogenetic analyses alone, and provides insight into the mechanisms of duplication and regulation operative within gene families^6,32^. Our results indicate that PBVP, including enzymatic components, are present as multi- or single-copy genes in genomic regions stable enough to facilitate comparative microsyntenic analyses. The stability of these regions across investigated taxa suggests that the origins of these genes are ancient, probably occurring in the most recent common ancestor of sawflies, parasitic wasps, and aculeate wasps. Exceptions to this pattern are the short, single-copy genes encoding toxic peptides known from bees such as apamin/MCDP/tertiapin, and melittin, which appear unique to bees or honeybees, indicating much more recent origins.

**Figure 3.**
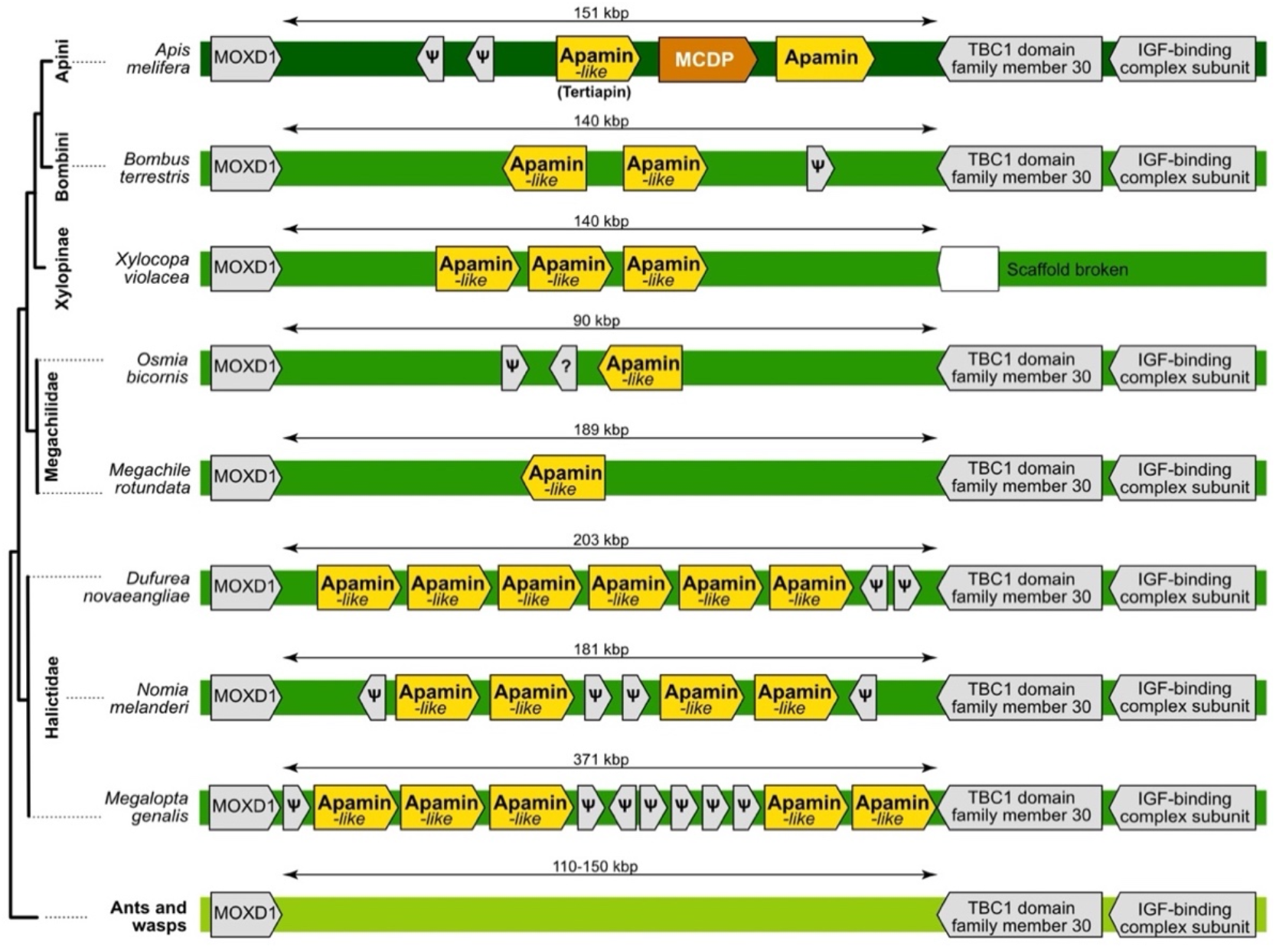
Microsyntenic pattern for the apamin family (Anthophilin1). Question marks indicate coding sequences with products of unknown functions. Pseudogenes are symbolized by ψ. The arrows reflect gene orientation. We show here only species for which the genomic sequence in the region with apamin genes is contiguous. Note that “apamin-like” genes are also known as “tertiapin”.

**Figure 4:**
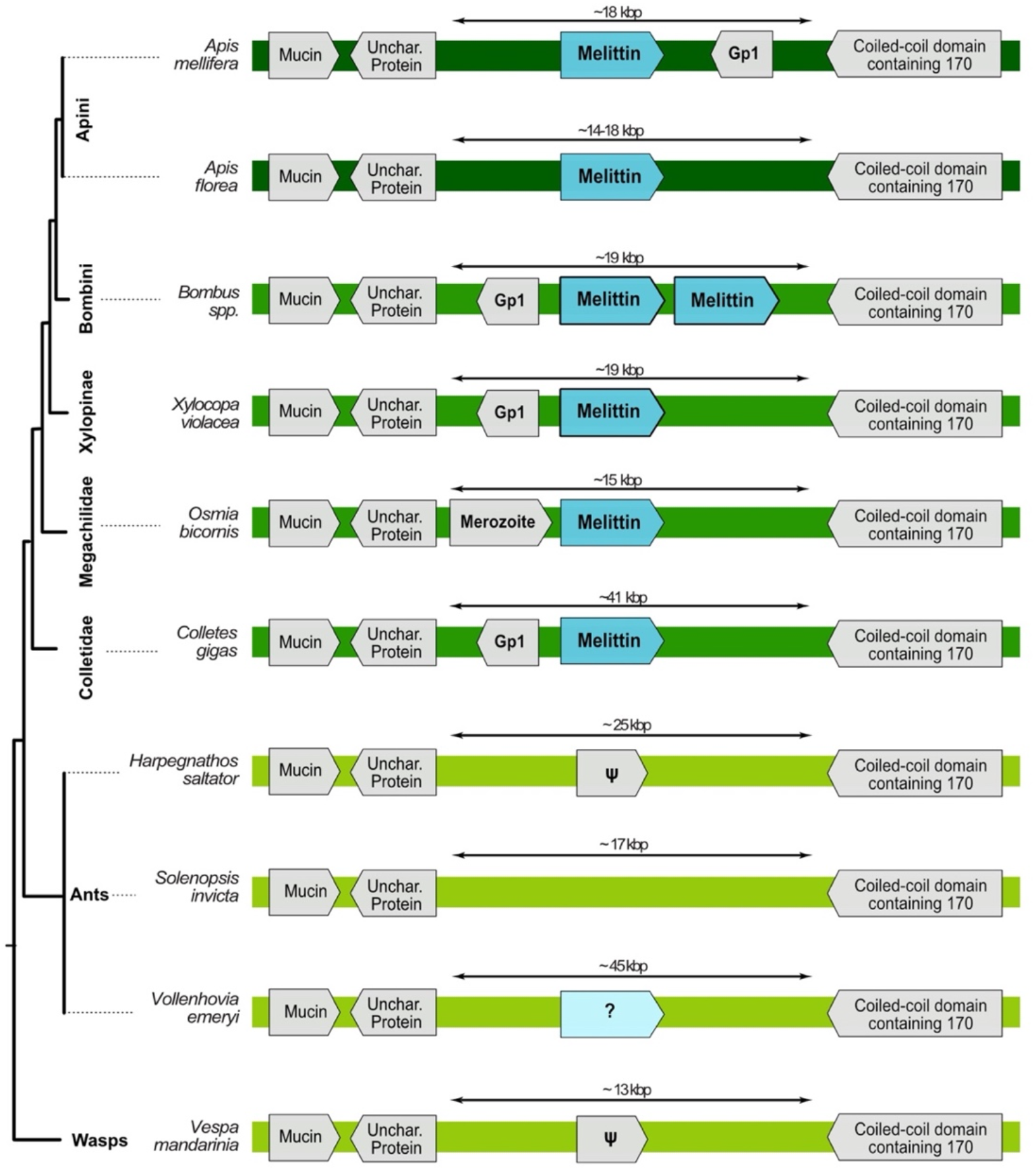
Microsynteny around the melittin sequence. All species for which the genome data allowed for microsyntenic analysis are shown. *Vollenhovia emeryi* was not included in other genomic analyses due to its relatively low genome quality. However, it is shown because it was the only one of the eight analysed ant species that features a seemingly related gene in the correct position but with a very different mature sequence. Genes labelled with ψ in ants and wasps bear little similarity with melittin genes, however, they might be sister genes to the melittin group that underwent severe pseudogenisation. Note, that *Osmia* melittin is also called “osmin”, *Colletes*’ – collectin, *Bombus* – bombolittin, *Xylocopa* – xylopin.

### Apamin is restricted to honeybees and is part of the larger bee unique toxin family Anthophilin1

Apamin, a dominant *A. mellifera* venom component, is encoded by a three-exon gene located next to a very similar three-exon gene encoding MCDP. This tandem duplication is flanked by MOXD1 homolog 2 and TBC1 domain family member 30. Although the two flanking genes are present and identically arranged in the genomes of all the bees we surveyed, we did not detect the full set of apamin or MCDP exons outside of the genus Apis (**Figure 3**). Genomic analysis confirmed that apamin and MCDP (from *Apis*) are restricted to the Apini clade (*Apis* spp.). In addition, we identified a novel apamin-like gene locus in *Apis mellifera* located right next to MCDP gene. This gene encodes the described honeybee toxin peptide named tertiapin^33^. Multiple uncharacterised genes that share microsyntenic position and intron-exon structure with this apamin-homologue (Tertiapin) were observed in Bombini and some other non-*Apis* bees. These apamin-like genes encode peptides that share the cysteine scaffold and signal peptide structure of apamin, MCDP, and tertiapin. They were widespread in bee genomes and we identified six copies in the *Dufourea* genome, five in *Nomia* and *Megachile*, two in *B. terrestris*, and a single copy in *Osmia bicornis, Habropoda* and *Megachile*. This pattern may be indicative of the derivation of apamin and MCDP from the more widespread tertiapin. We identified no similar genes or exons of apamin in homologous regions from other hymenopterans or in other parts of their genomes.

The apamin-like sequences we discovered in the core venom profile of *Xylocopa* and *Halictus* indicate that apamin and MCDP are members of a variable bee-unique family of apamin-like peptides that undergoes independent duplication events in different lineages. We propose here to name this novel family Anthophilin1, reflecting its uniqueness to several lineages within bees (Anthophila), see **Supplementary Figure 1** and **Supplementary File 1** for phylogenetic alignment and tree.

### Melittin is restricted to the bee lineage

Melittin is a pain-inducing peptide in *A. mellifera* venom^24,34^. The synteny of the *A. mellifera* genome shows that melittin is encoded by a two-exon single-copy gene located between two four-exon genes, one of which encodes *vegetative cell wall protein gp1* while the other remains uncharacterized. Melittin-like sequences in other *Apis* species (*A. dorsata, A. cerana* and *A. florea*) feature similar microsynteny (**Figure 4**). Other bee species also possess melittin-like sequences (bombolittin, osmin, collectin, lasioglossin, melectin, codesane, halictin and macropin^35, 39^. Microsynteny analysis provided evidence that osmin, collectin, bombolittin and xylopin are orthologous in at least some species from the genera *Colletes, Osmia* and *Bombus* (**Figure 4**).

In *Bombus vosnesenski*, the melittin gene has undergone a tandem duplication that is apparently unique to *Bombus*. Some *Bombus* genomes show assembly gaps in this region, preventing the detection of all exons, but recently published genomes of several *Bombus* species^40^ show the same sequence and duplication pattern in the microsyntenic region identified in *B. vosnesenski* (**Figure 4**). Although tracing the corresponding genomic region in non-bee Aculeata proved to be difficult because of its relative instability (low synteny/colinearity), we successfully located it in ants and wasps, which lacked melittin homologues. However, one ant genome – *Vollenhovia emeryi* (excluded from our main genomic analysis due to the relatively low genome contiguity) – had a superficially similar looking gene in almost the exact location **(Figure 4)**. That gene has a proline-rich propeptide resembling that of melittin, nevertheless, its mature form is very different. We conclude here that our results support the hypothesis that melittin is restricted to bee lineages, however, its ancestral gene might have had homologs in ancestors of wasps and ants, see **Supplementary Figure 2** and **Supplementary File 2** for phylogenetic alignment and tree.

### A machine learning model of “protein space” does not support the aculeatoxin hypothesis

Given the short peptide sequences and consequent challenges for phylogenetic analyses we utilised a novel, sequence-independent machine learning approach (see methods section for the detail) focused on melittin to test the proposition by Robinson et al.^26^ that based on signal and propeptide melittin is a member of ‘aculeatoxins’ a family that origins with aculeates (**See Figure 1**). These analyses generate a model of the relations of proteins to each other in a 3-dimensional “protein space” similar to the concept of a “configuration space” in physics, or an “arbitrary space” in multi-scale cognition^41^. Our protein space incorporates data concerning the structure and function of mature proteins to generate a 3-dimensional model of protein relations. By observing the clustering patterns of proteins within this 3-dimensional space, we can infer their evolutionary relations to one another. These analyses used all sequences that Robinson et al.^26^ presented in their study (kindly provided to us by the authors) and all melittin-like toxins known from bees included in our study. We created two datasets – with (**Supplementary File 3**) and without **(Supplementary File 4**) signal/propeptides **–** the latter dataset includes more sequences since bee melittins are mostly known from proteomic studies and therefore only their mature sequence is known. The protein space occupied by peptides from wasps and bees was distinct from that occupied by ant peptides, and thus these analyses do not support the aculeatoxin hypothesis (**Figure 5**). However, the results reveal a close similarity between bee and wasp peptides, which was even more apparent when signal peptides were removed (in contrast to the reasoning of Robinson et al., which is based on similarity amongst signal peptides). Microsyntenic analyses reveal that melittin, mastoporans, and poneratoxins are non-homologous, however the protein space analyses may reveal evidence of convergence. Taken together, the results of these analyses indicate that melittin is likely unique to bees but gravitates towards mastoporans (in particular) and poneratoxins (to a lesser extent) in protein space, possibly because of functional convergence. However, addressing the aculeatoxins hypothesis in detail goes beyond the scope of this manuscript and therefore will be a subject of a follow-up study. Nevertheless, it is worth mentioning that all of the proposed members of aculeatoxins are processed by the same enzyme DPP4, which might explain similarities in signal and propeptide sequences.

**Figure 5:**
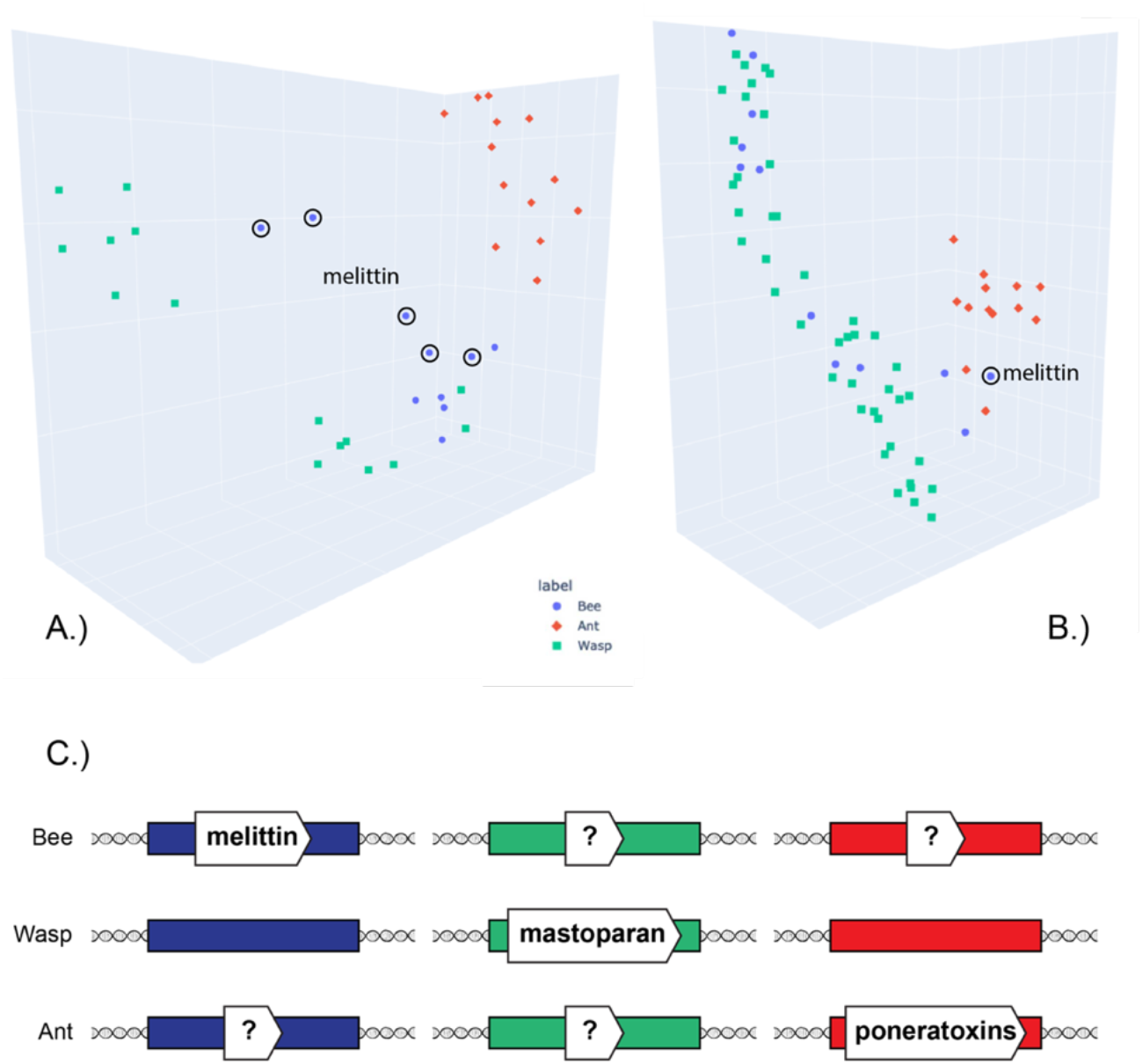
“Protein space” of small peptidic aculeatan toxins as revealed by machine learning analysis and their genomic position in respect to each other. Combined data of available verified toxin sequences from Robinson et al., and the present study. A.) Sequences with signal peptide included B.) Only mature peptides. Melittin sequences are outlined with a black circle, different ones on the panel A represent different transcripts with the same mature peptide. For the interactive plots see **Supplementary Files 5 and 6**. C.) Schematic of genomic position of the three groups of hymenopteran toxins. Coloured rectangles represent regions of microsynteny: blue for melittin, green for mastoparan and red for poneratoxins. See text for details.

### Abundant venom proteins are encoded by more widespread single-copy genes

Phospholipase A2, hyaluronidase and icarapin are among the most abundant bee venom components^16,17,24^. Phospholipase A2 and icarapin are encoded by four-exon single-copy genes, whereas the hyaluronidase single copy-gene features nine exons. Dipeptidyl peptidase-4 has a strongly conserved single gene, which was present in all hymenopterans in our dataset, probably due to its enzymatic role in the maturation of some toxins. These protein families were highly conserved and ubiquitously present in the genomes of bees, wasps and ants (**Figure 6,** see also **Supplementary Files 7-10** and **Supplementary Figures 3-6** for phylogenetic alignments and trees). Our results support the hypothesis that these genes were recruited into venom functions without any associated duplication – similar to cooption of single-copy genes proposed as the main process of venom protein evolution in *Nasonia*^42^. In comparison, phospholipase A2 genes in viperid snakes had multiplied and diversified before recruitment into the venom system^43,44^.

**Figure 6.**
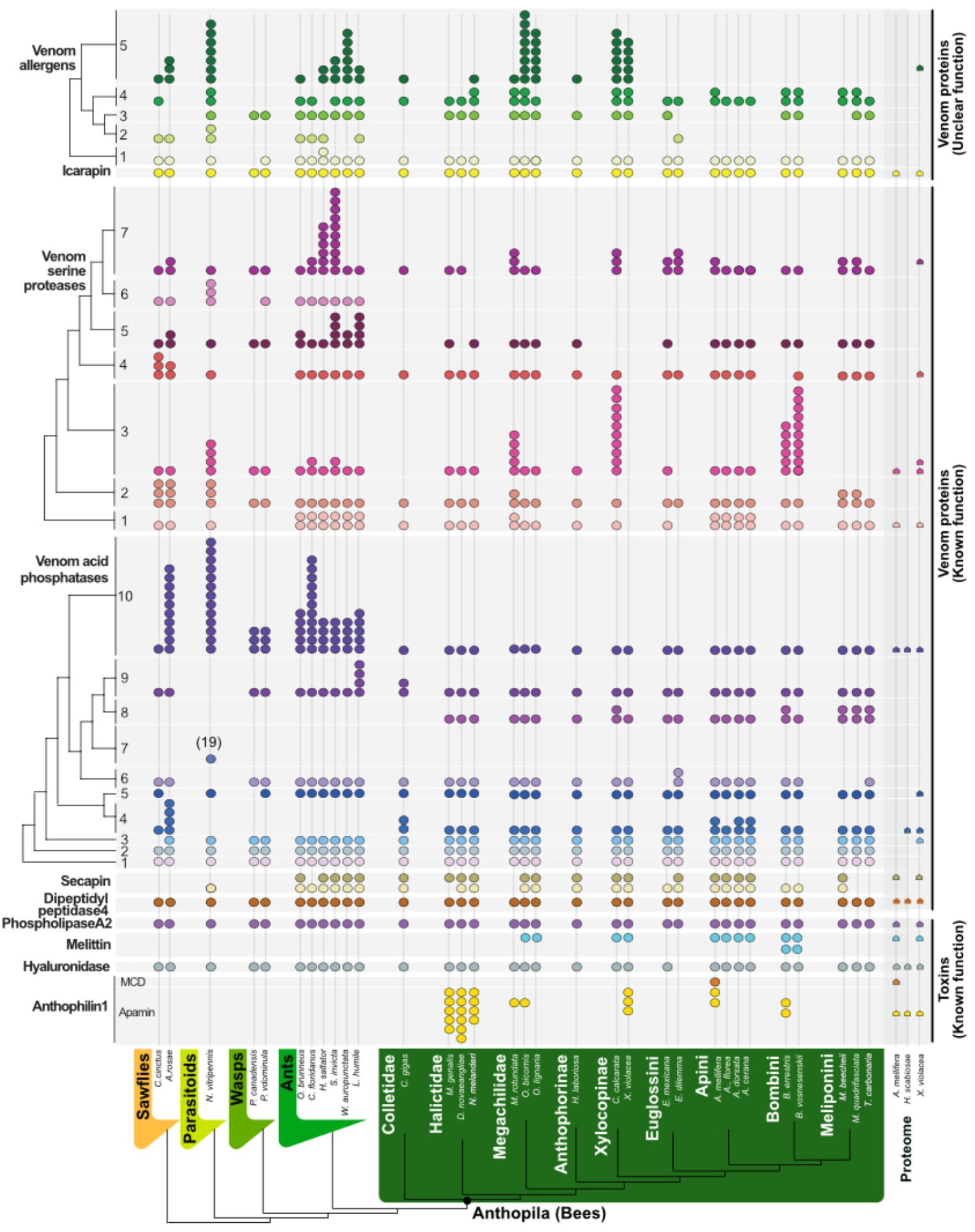
Overview of prevalent bee venom genes. The presence of venom gene orthologs and copy number variation is mapped onto the phylogenetic relationship between the species we surveyed according to Peters et al^31^. Coloured circles represent genes with identical microsynteny in the genomes of the surveyed species. Please note that tertiapin is now included within anthophilin1 as variant of apamin.

### Some venom proteins form multi-copy gene families with ancient duplication events

Larger duplication and diversification events appear restricted to families of enzymatic or larger proteins and not toxin peptides or proteins. Three venom protein classes in the PBVP showed copy number variation across the dataset: venom allergens 3/5, venom acid phosphatases, and serine proteases (**Figure 6**). These genes were in stable genomic regions allowing for the tracing of homologous regions between species by screening for microsynteny.

Among the 10 subfamilies of venom acid phosphatases (**Figure 6**) the largest expansion of genes occurred in subfamily 7, found exclusively in parasitoid wasps. This may support the hypothesis that ancestral APHs functioned as pre-digestion factors that allowed the offspring of parasitoid wasps to feed more easily on their host^45^. In contrast, gene expansion in subfamily 10 appears to be an ancient pattern found in sawflies (9 genes) and parasitoid wasps (13 genes). In all remaining hymenopterans only one or occasionally two to three genes are present. A similar pattern was observed for subfamily 5 with ant species having 2-4 copies, while all other hymenopterans (with the exception of *Athalia*) have 1. Subfamily 3 seems to have undergone multiple duplication events in some bee species with up to 10 copies in *Ceratina* and *Bombini*, while other species have 1-2 copies or lost all genes (*Meliponini*), see **Supplementary File 11** and **Supplementary Figure 7** for phylogenetic alignment and tree. In bees, the retained APHs may be adapted to defensive functions, a conjecture potentially supported by the origin of APH subfamily 8, which is unique to bees.

Our analyses divided venom serine proteases (VSPs) are into seven subfamilies. Subfamily 7 is represented by 1–4 genes in all hymenopterans but has expanded in ants (10 genes). All seven subfamilies are present in the basal lineages of sawflies and parasitoid wasps, with more diversification in families 2, 3 and 4. In bees subfamily 6 appears to have been lost (**Figure 6**; see **Supplementary File 12** and **Supplementary Figure 8** for phylogenetic alignment and tree). VSPs are dual function toxins in bees, triggering the phenoloxidase cascade leading to melanization when injected into insects but acting as spreading factors when injected into mammals, similar to snake VSPs with fibrinogen-degrading activity^46^. We hypothesize that the expansion of VSP genes may be linked to this dual function, achieving more effective defense against insects, arthropods and mammals.

Venom allergens 3/5 have been identified in many hymenopterans^16,47^ and we distinguished five subfamilies in our study. Subfamily 5 appears to have undergone greater diversification in sawflies, parasitoid wasps, ants and the solitary bees (*Ceratina, Osmia*). Only a single member of subfamily 5 is present in the solitary bees *Habropoda, Colletes* and *Nomia*. Eusocial wasps and bees of the family Apidae (*Apis, Bombus, Melipona, Frieseomelitta* and *Eufriesea*) appear to have lost all subfamily 5 genes. Subfamily 1 is present only in parasitoid wasps and ants with a single gene in *Euglossa*. Other subfamilies generally have a single copy in every species with subfamily 4 occasionally experiencing duplication. In general, the distribution of genes in the venom allergen family is dynamic but shows some phylogenetic patterns (see **Supplementary File 13** and **Supplementary Figure 9** for phylogenetic alignment and tree).

Two secapin genes were present in most genomes, but were absent in sawflies (indicating an origin in the stem Apocrita) and wasps of the genus *Polistes*. This class of peptides displayed N-terminal sequence variation but strong C-terminal conservation (see **Supplementary File 14** and **Supplementary Figure 10** for phylogenetic alignment and tree). The location of both genes was also strongly conserved, with one always present between exons of the neurexin-1 gene and the other located near the carbonic anhydrase-related protein 10. Our inability to locate both genes in some species may reflect technical issues relating to genome quality and/or the more general challenges associated with the location of small and highly variable genes.

## Discussion

### Gene expansions are restricted to few venom protein families in major taxa

Most PBVP are encoded as single-copy genes (**Figure 6**), indicative of single gene co-option. Our data supports the hypothesis that gene duplications are a less prevalent evolutionary mechanism in the evolution of hymenopteran venom components than is claimed for (e.g.) snakes. This pattern was previously observed in parasitoid wasps (*Nasonia*)^42^. However, our results indicate a more distinct pattern in which heavier protein and enzyme components represent those families of venom proteins in which large gene duplications and expansions have occurred in conserved genomic regions. These expansions are restricted to particular subfamilies and larger hymenopteran clades (**Figure 6**). The gene duplications and subsequent gene expansions of venom serine proteases, venom allergens and venom acid phosphatases appear to be ‘simple ‘events restricted to the expansion of few genes. This is in contrast to other venomous organisms that have been studied more extensively, such as snakes and cone snails, in which venom genes have evolved rapidly by extensive multiplication, expansion and subsequent deletion^8,48–51^. It should be noted, however, that this picture is based on our preselected PBVP, which includes the most common venom components described.

Venoms are secretions which primarily function (when “actively delivered” via bites or stings) to deter or subdue target organisms. Venoms contain a variety of molecules and not all are necessarily associated with the primary function of the secretion. Some are of as yet unknown function, or may be epiphenomenal (i.e. present in venoms for contingent reasons not associated with any particular functional role). Whilst we use the term “venom protein” (or “component”, or “gene”) to refer to any molecule *associated* with venom (i.e. detected proteomically or transcriptomically within the venom system), we reserve the term “toxin” for those venom components with a *characterised functional role* in the subjugation or deterrence of target organisms. Our results indicate that genes encoding (characterised) toxins and those encoding other (associated) venom proteins evolve differently in bees, suggesting a genuine functional distinction between these groups. This finding should be tested further in the future using extended venom profiles. Complementary activity studies are important to address the still undefined biological functions of many venom components, for example venom “allergens”, which would in turn support a better interpretation of evolutionary patterns. Venom allergens (3/5) show a more heterogeneous pattern of gene duplications than other gene families, especially in subfamily 5. This subfamily has expanded in parasitoid wasps, leafcutter bees (Megachilidae) and carpenter bees (Xylocopinae), but has been lost in other Apidae lineages. We can only speculate about the original and actual biological function of venom allergens in general because until today the only *activities* characterised are related to immune responses in mice and humans linked to allergic reactions^52^. No study so far has addressed the possible bioactivity linked to the ancestral and venom variant’s biological *function*. However, the strong allergenic activity may reflect an ancestral immunomodulatory function in sawflies linked to the modulation of the immune response of plants, which was later adapted to animal hosts in more derived aculeate lineages.

### Bee-specific toxin genes encoding for short peptides

Bees produce apamin and melittin as predominant venom components^24^, but their genomic origin beyond the honeybee lineage has not been investigated before. One major difference between these toxin peptides and previously discussed venom components is that the genomic region in which they are encoded appears more dynamic. This picture is also reflected by taxon-restricted gene duplications. The genomic region containing a tandem repeat of apamin and mast cell degranulating peptide in *Apis* was identifiable in other bee genomes based on microsynteny and the characteristic cysteine scaffold. Interestingly, we discovered multiple duplication events each restricted to single bee lineages. Our conclusion based on this pattern is that apamin and mast cell degranulating peptide are members of a so far unrecognized, highly variable bee-unique peptide family, which we named Anthophilin1. The genes of this family seem to diversify independently in different bee lineages. Interesting is, that in snakes and sea anemones the expansion of toxin gene families is shown to be linked to their selection to generate larger quantities of the venom than novel function^9,11^. Whether the duplication events are linked to neofunctionalization or co-option (as one dominant venom component) remains to be addressed in future studies, the scenarios of gene duplication in venom evolution can be more complex than they often appear^6^. These should include more contiguous genomic data from additional bee lineages and complementary venom proteomes to better understand the recruitment and diversification processes of members of this family in bee venom.

We identified melittin in a genomic region with conserved synteny in the genera *Apis, Osmia, Ceratina* and *Bombus* (families Megachilidae and Apidae), with a tandem duplication in bumblebees. Synteny confirmed that melittin-like peptides produced by solitary bees are members of the melittin family. Accordingly, melittin is not unique to *Apis* but originated before the divergence of megachilid and apid bees. We did not find a syntenic region or sequences similar to melittin in genomes of bees from the families Andrenidae, Halictidae and Colletidae. Whether or not melittin evolved in earlier bee lineages and underwent secondary loss in some families remains unclear from our data due to the lack of high-quality genome assemblies for the early-diverging bee lineages. Our data further indicates that the ant *Vollenhovia* features a gene which may be distantly related to melittin, however, the mature sequence looks very different. Nevertheless, we cannot rule out the possible origin of melittin in earlier aculeate lineages until a larger sampling of taxa from these and earlier bee lineages are available with high-quality proteo-transcriptome-genome data. Regardless, our data suggests that melittin is co-opted as a single copy gene as one major component in bees. In future studies this hypothesis should be further tested by analysing more proteo-transcriptomic venom profiles linked to genomic data.

### Most bee core venom proteins originated in early hymenopterans

The pattern we infer reveals an ancient origin for most of the PBVP in bees (**Figure 7**). Most subgroups of major venom protein gene families exhibit clear-cut orthology with genes already present in the earliest hymenopteran lineage (sawflies). Female sawflies use their ovipositor to lay eggs in plants but also co-inject proteins that biochemically interfere with the physiology and immune response of plants to ensure the offspring’s survival, thus resembling a primitive venom system^15^. The composition of these original hymenopteran venoms has not yet been studied in detail.

**Figure 7.**
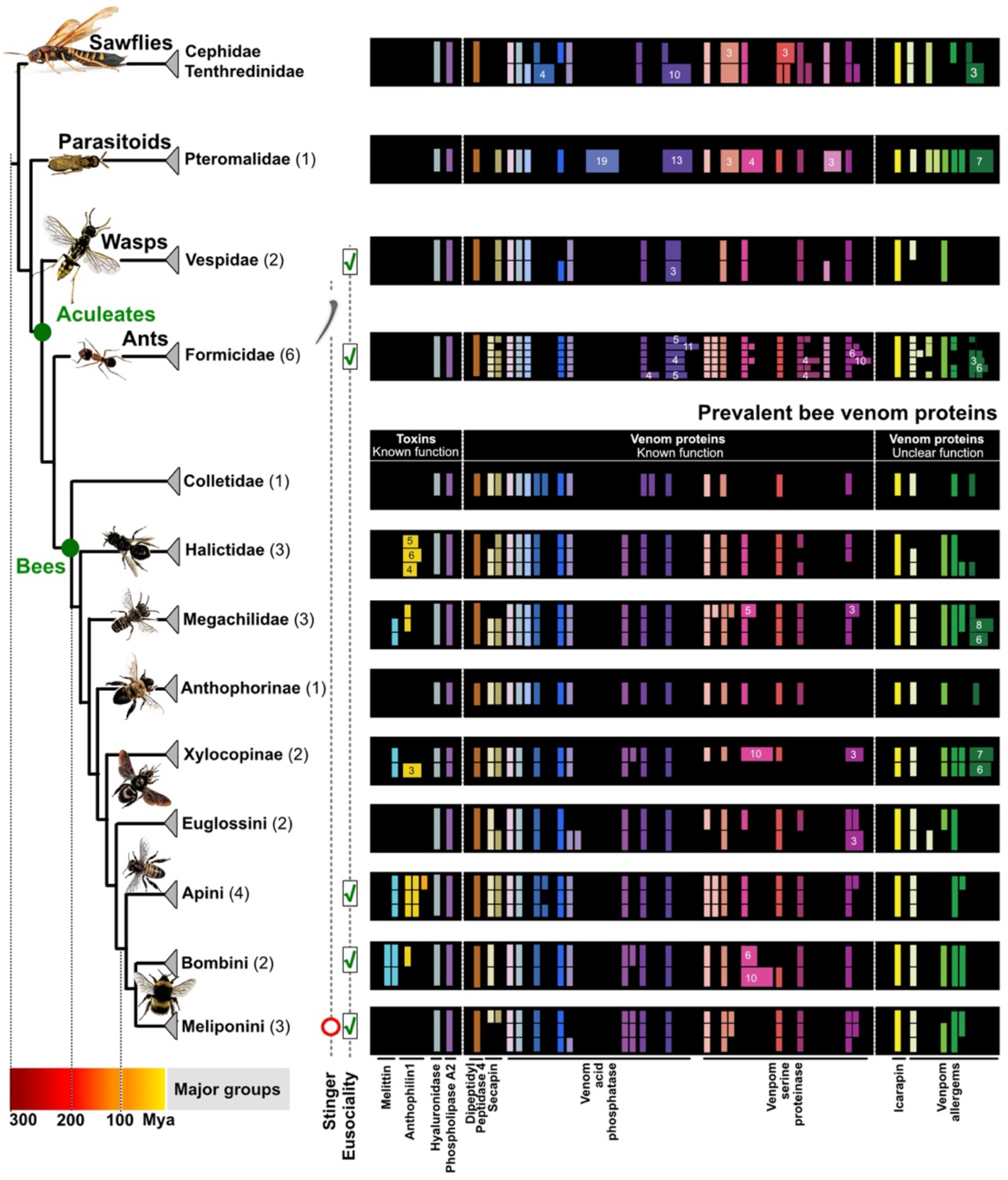
Simplified visualization of the prevalent bee venom proteins and their representation in outgroup taxa. The numbers of genomes are shown in brackets after the family names. Genes are colour-coded and feature a colour range for duplicates. Duplications are summarized by numbers. Phylogeny and divergence times are shown as previously described in Peters et al.^31^.

Our results suggest that the most prevalent venom genes present in bees today were already present in the early Triassic in ancestors of the symphytan lineage, predating the radiation of apocritans starting more than 200 million years ago (**Figure 7**)^31^. The restricted waist of apocritans is needed to manoeuvre the ovipositor in such a way that allows its use for predation, parasitism or defense, and only in aculeate hymenopterans (ants, bees and wasps) is the retractable ovipositor modified into a stinger used exclusively for venom injection. Our data suggest that genes encoding the PBVP emerged before the morphological adaptations of a narrow waist and the stinger in aculeates did, which gave this group its common name – the stinging wasps. The core of the bee venom profile, including known allergens such as phospholipase A2, icarapin and hyaluronidase, was not only already present in sawflies, but is also still present in a group of bees that has secondarily reduced or lost its stinger (stingless bees, Meliponini).

If one accepts Symphyta as ‘venomous’, based on their injection of molecules that modulate the physiology (particularly the immune system) of target organisms (to facilitate feeding of the next generation, similarly to parasitoid wasps), then one might consider the hymenopteran lineage as ‘descending from a common venomous ancestor’. Indeed, this might be much less controversial an assertion for this order than it has turned out to be for toxicoferan reptiles (see, e.g.^29^ and subsequent discussion in the journal *Toxicon*). In this case, our data is consistent with the idea of continuous evolution (i.e. without sharp distinctions or saltatory events) of the hymenopteran venom system through various changes in associated anatomy and ecology. The core of the venom arsenal, comprised of larger proteins which function as immunomodulators or spreading factors, may have been in place early on. Subsequent evolution focused then on the origin and diversification of lineage-specific arrays of peptides which are tailored to the specific venom function (e.g. defence, parasitism, predation) and target (plants, insects, vertebrates) in each lineage. Thus, whilst the peptidic toxins are unique to each lineage within the Aculeata (contrary to the aculeatoxin hypothesis), most enzymatic components are broadly shared, albeit with varying degrees of expansion of specific subfamilies. These differential expansions of enzyme-encoding gene families (e.g. serine proteases) may represent the kind of evolutionary tinkering observed in redundant arrays of toxin-encoding genes in other venomous taxa (see e.g., Jackson et al.^49^), in which slight changes confer adaptation to the biochemical particularities of a new ecological reality. Members of such enzyme classes may thus vary in their activity on specific substrates, linked to modified morphology of the venom apparatus, but are never rendered inactive due to broadly applicable modes of action (i.e. targeting substrates generally conserved across taxa as diverse as plants and vertebrates). The subject of this study, bees, seem to support this view by having little variation in their venom genes, other than within the Anthophilin1 and Melittin groups.

## Conclusion

Our comparative analyses provide insight into the origins and evolution of toxin genes in bees. We found that most genes encoding predominant bee venom proteins originated at the base of the hymenopteran tree, i.e. were potentially present in the ‘venom’ of the last common ancestor of phytophagous sawflies and apocritan Hymenoptera more than 280 million years ago (**Figure 7**). Only the short peptides melittin and the (herein newly described) family Anthophilin1, which is constituted by apamin, apamin-like and MCDP-like genes, are unique to bees. Gene duplications occur, but only in certain (not major toxin) protein families and in only a few hymenopteran lineages, reflecting a diverse pattern of gene origin. Our results thus indicate that short peptides and venom protein genes probably evolve under different evolutionary processes. This study of the PBVP demonstrates the requirement for future studies to provide insight into the evolution of bee and hymenopteran venoms. These should include more high-quality genomes, especially for early bee lineages but also including a more widespread taxon sampling of other hymenopterans. More importantly, extended proteo-transcriptomics venom profile data are essential. Our data shows that venom compositions for solitary bees can be heterogeneous (especially for smaller species such as *H. scabiosae*) which needs to be accounted for. Corresponding to genomes this proteo-transcriptome data improves genome annotations and allows to address appropriately venom protein recruitment processes to differentiate more precisely between gene variants expressed in the venom system and non-venom related genes. Finally, bioassays for many still unknown venom components are needed to identify functional differences linked to the gene evolution and diverse ecology of hymenopteran species.

## Materials and methods

### Data mining of hymenopteran venom proteins and genomes

Reviewed venom proteins of hymenopterans were searched in UniProt resulting in 372 protein matches from 101 species (**Figure 1 and Supplementary Table 4).** Additionally, we searched publications for sequences that are not provided in UniProt and included finally three bee toxins Halictin I and II from *Halictus sexcintus*, and Codesan from *Colletes daviesanus*. For our comparative genomic analysis of venom toxin proteins across the order Hymenoptera, we made use of **29** publicly available genome sequences given in **Supplementary Table 5** and three novel genomes of solitary bees.

### Venom gland RNAseq analyses

For venom gland transcriptomics 15 individuals of *X. violacea*, 17 individuals of *H. scabiosae* and 15 individuals *of A. mellifera* were collected June-July 2019/2020 in the alluvial area of the River Wieseck in Giessen, Germany, and the beehive at the Institute for Insect Biotechnology at Justus-Liebig-University Giessen (Collection permission HNLUG Giessen IV.2 R28). Whole venom systems (Glands and reservoir) were dissected and washed on ice under sterile conditions and the tissue was preserved in RNAlater (Thermo Fisher Scientific) for subsequent RNA sequencing. RNA extraction, library preparation and short-read genome sequencing were outsourced to Macrogen (Seoul, Korea) for *A. mellifera* and *X. violaceae* and to Novogene (Cambridge, UK) for *H. scabiosae*. In short, RNA was extracted with Trizol and the cDNA libraries (150bp, paired end reads) were sequenced using a low input protocol (Illumina Truseq) on an Illumina HiSeq2500 (Macrogen) and Illumina NovaSeq (Novogene). For *H. scabiosae* an in-house ultra-low input protocol was used by Novogene due to very low RNA concentration and quantity. All raw data are submitted to NCBI GenBank PRJNA733472 (SRA entries: SRR14690757, SRR14690758, SRR14690759). Venom gland transcriptomes were assembled separately using Oyster River Pipeline v2.2.6^53^, for resulting BUSCO values see **Supplementary Table 7.**

The resulting assemblies were processed using Transdecoder (minimum length 20 amino acids) to predict peptides, and Kallisto v0.46^54^ to calculate individual transcript abundance, see **Additional Files 1-3**. The assembled transcripts and their corresponding longest ORFs (Transdecoder output) were used as local BLAST queries against ToxProt and UniProt (the latter limited to insects only) with an e-value cutoff of 1 × 10^-3^, see Figure 8. Any highly abundant (TPM > 100) transcripts without significant matches were manually screened using BLAST, InterPro scan and ProteinPredict online suites to determine the closest characterized homolog. For subsequent venom protein identification, we only included transcripts identified in our proteomic dataset representing proteins secreted in the venom system. To compare subsequently all venom proteins in the three datasets we calculated the percentage of scaled TPMs using the package txtimport on R, the script is available via github (https://github.com/marivelasque/VenomEvolution.git), see **Figure 2** and **Supplementary Tables 1-3**.

**Figure 8.**
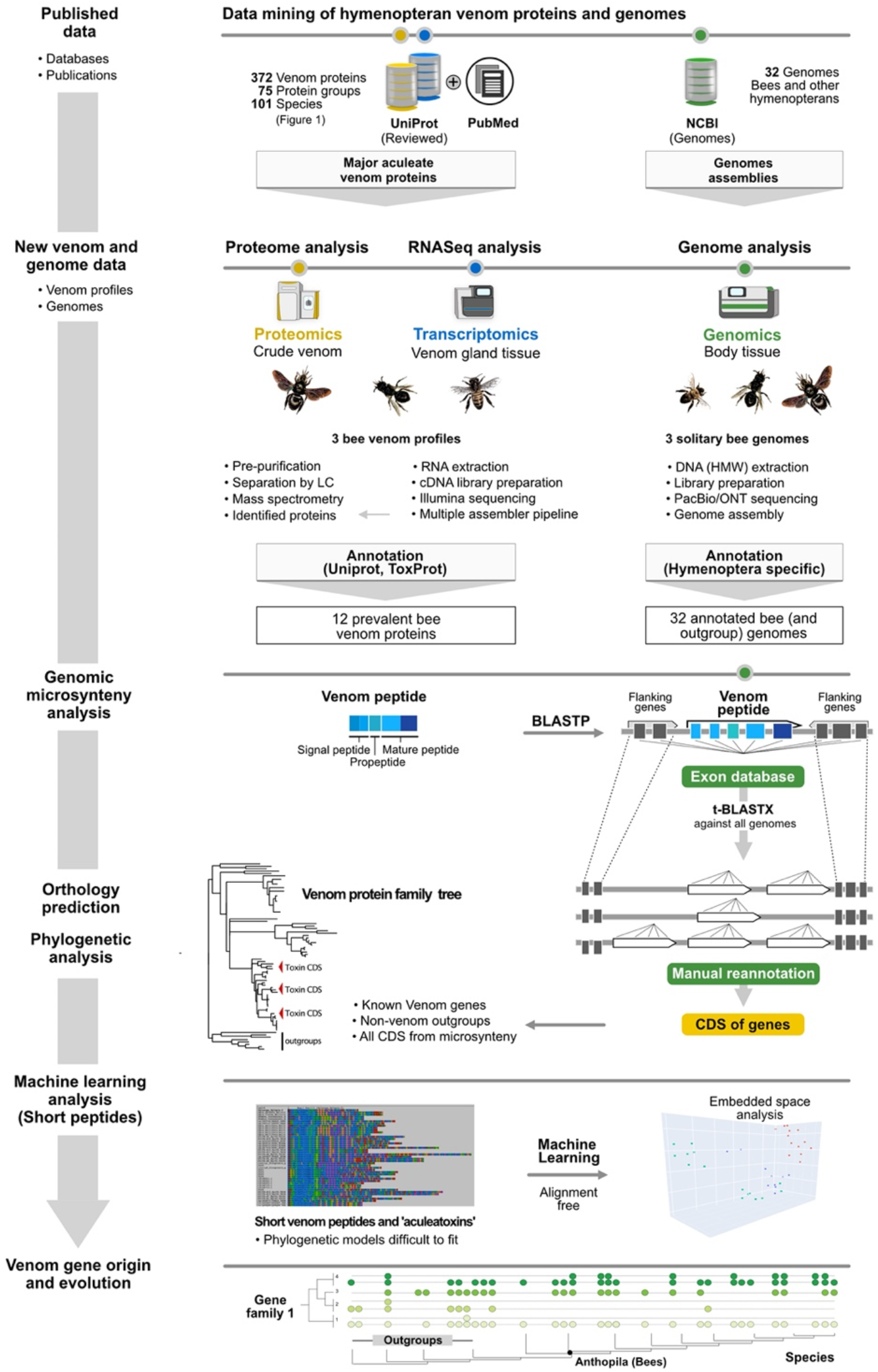
Description of the proteo-transcriptomic and genomic workflow applied in this study. Details of each step are given in material and methods.

### Proteome analysis of crude venom

We extracted crude venom of all specimens from glands and venom reservoirs by squeezing with forceps in sterile ultrapure water (Thermo Fisher Scientific, Waltham, MA, USA) after prewashing twice to minimize hemolymph contamination. All transcriptome assembly-based predicted ORFs were used as specific databases to identify peptides and proteins detected by mass spectrometry from crude venom of the collected specimens. For the tryptic digestion of the crude venom from *H. scabiosae*, we dissolved 10 μg of protein in 10 μl 10 M urea containing 0.1% ProteasMax (Promega. Madison, WI, USA). Cysteine residues were reduced with 5 mM DTT (30 min at 50 °C) and modified with 10 mM iodoacetamide (30 min at 24 °C). The reaction was quenched with an excess of cysteine and trypsin was added at a protein:enzyme ratio of 40:1 in 100 μl 25 mM ammonium bicarbonate (Sigma-Aldrich, Taufkirchen, Germany). After incubation for 16 h at 37 °C, the reaction was stopped by adding 1% trifluoroacetic acid (TFA). The sample was purified using a C18-ZipTip (Merck-Millipore, Darmstadt, Germany), dried under vacuum and redissolved in 10 μl 0.1% TFA. LC-ESI-MS analysis was carried out at 35 °C by loading 1 μg of the sample in 0.1% formic acid (Sigma-Aldrich) onto a 50-cm μPAC C18 column (Pharma Fluidics, Gent, Belgium) mounted on an UltiMate 3000RSLCnano (Thermo Fisher Scientific). Peptides were eluted with a linear gradient of 3–44% acetonitrile over 240 min followed by washing with 72% acetonitrile at a constant flow rate of 300 nl/min. They were then infused via an Advion TriVersa NanoMate (Advion BioSciences, New York, NY, USA) into an Orbitrap Eclipse Tribrid mass spectrometer (Thermo Fisher Scientific) in positive-ionization mode with a NanoMate spray voltage of 1.6 kV and a source temperature of 275 °C. Using data-dependent acquisition mode, the instrument performed full MS scans every 3 s over a mass range of *m/z* 375–1500, with the resolution of the Orbitrap set to 120,000. The RF lens was set to 30%, and auto gain control (AGC) was set to standard with a maximum injection time of 50 ms. In each cycle, the most intense ions (charge states 2–7) above a threshold ion count of 50,000 were selected with an isolation window of 1.6 *m/z* for higher-energy C-trap dissociation at a normalized collision energy of 30%. Fragment ion spectra were acquired in the linear ion trap with the scan rate set to rapid, the mass range to normal and a maximum injection time of 100 ms. After fragmentation, the selected precursor ions were excluded for 15 s for further fragmentation.

Prior to shotgun proteomics, the *X. violacea* and *A. mellifera* venom samples were denatured, reduced, and alkylated. Briefly, each sample (~50 μg) was dissolved in 89 μl 100 mM triethylammonium bicarbonate (TEABC), and cysteine residues were reduced by adding 1 μl 1 M DTT (30 min at 60 °C) and modified by adding 10 μl 0.5 M iodoacetamide (incubation for 30 min in the dark). We then added 2 μg trypsin (Promega) in 100 mM TEABC and incubated overnight at 30 °C. The peptides were then purified and concentrated using OMIX Tips C18 reversed-phase resin (Agilent Technologies, Santa Clara, CA, USA). The peptides were dehydrated in a vacuum centrifuge and analysed by NanoLC-MS/MS. The samples were then resuspended in 20 μl buffer A (0.1% formic acid) and 1 μl was loaded onto an analytical 25 cm reversed-phase column (Acclaim Pepmap 100 C18) with a 75 mm inner diameter (Thermo Fisher Scientific) and separated on the Ultimate 3000 RSLC system coupled via a nano-electrospray source to a Q Exactive HF-X mass spectrometer (Thermo Fisher Scientific). Peptides were separated using a 6–40% gradient of buffer B (80% acetonitrile in 0.1% formic acid) over 123 min at a flow rate of 300 nl/min. Using data-dependent acquisition mode, full MS/MS scans (375–1500 m/z) were performed in the Orbitrap mass analyser (Thermo Fisher Scientific) with a 60,000 resolution at 200 m/z. For the full scans, 3 × 106 ions accumulated within a maximum injection time of 60 ms. The 12 most intense ions with charge states ≥ 2 were sequentially isolated to a target value of 1 × 105 with a maximum injection time of 45 ms and were fragmented by higher-energy collisional dissociation in the collision cell (normalized collision energy 28%) and detected in the Orbitrap mass analyser at a resolution of 30,000. PEAKS Studio v8.5 (Bioinformatics Solutions, Waterloo, ON, Canada) was used to match MS/MS spectra from *X. violacea* and *A. mellifera* venom samples against an in-house database resulting from the annotated transcriptome of each species. Carbamidomethylation was set as a fixed modification, and oxidation of methionine as a variable modification, with a maximum of three missed cleavages for trypsin digestion. Parent and fragment mass error tolerances were set at 5 ppm and 0.015 Da, respectively. A false discovery rate (FDR) of 1% and a unique peptide number ≥ 2 were used to filter out inaccurate proteins. A −10lgP value > 120 was used to estimate whether detected proteins were identified by a sufficient number of reliable peptides. In order to identify more relevant sequences, the Spider algorithm (PEAKS Studio) was used to find additional mutations or to correct sequences. This algorithm corrects the sequences stored in transcriptomic databases with de novo sequences based on MS/MS spectra, allowing the detection of post-translational modifications (PTMs) and mutations. The minimum ion intensity for PTMs and mutations was set to 5%, and the ALC score was set to ≥ 90 for de novo sequences, leading to low precursor mass errors. Transcripts supported by proteomic data were manually filtered by excluding non-venom-related proteins and peptides, such as house-keeping and structural genes (**Supplementary Tables 1-3**). All proteome raw data are submitted to PRIDE (PXD029934, PXD029823, PXD026642).

### Genome sequencing

The genomes and annotations of the stingless bees *Tetragobula carbonaria* and *Melipona beecheii* will be published as part of another study, but have already been uploaded to NCBI. To sequence the genome of *X. violacea* high molecular weight DNA was extracted from four legs of *X. violacea* adapting the protocol from Miller et al.^55^. Final DNA purity and concentrations were measured using NanoPhotometer^®^ (Implen GmbH, Munich, Germany) and Qubit Fluorometer (Thermo Fisher Scientific, Waltham, MA). Two SMRTbell libraries were constructed following the instructions of the SMRTbell Express Prep kit v2.0 with Low DNA Input Protocol (Pacific Biosciences, Menlo Park, CA). The total input DNA for each library was 1.6 μg. The libraries were loaded at an on-plate concentration of 80 pM using diffusion loading. Two SMRT cell sequencing runs were performed on the Sequel System IIe in CCS mode using 30-hour movie time with 2 hours pre-extension and sequencing chemistry v2.0. The PacBio sequencing was outsourced to the Genome technology Center Nijmegen, Netherlands. All reads were assembled using HIFIASM assembler^56^ after fastq read files of *Xylocopa sp*. were generated by consensus calling of Pacbio HIFI sequencing data using CCS tool (https://github.com/PacificBiosciences/ccs). Reads, which did not take part in the formation of circular consensus sequences were separated out using in-house developed Perl script and were used for closing the gaps with the help of Dentist software^57^. The gap-closed assembly was further polished using Bowtie2^58^, Deepvariant^59^, Samtools and BCFtools^60^. Contamination was accounted for by using NCBI Blast and Blobtools^61^, and only scaffolds with Arthropoda and No-Hit category were kept. The final gap-closed and contamination free genome of *Xylocopa species* consisted of 353045797 bases spread over 3524 scaffolds. The genome was predicted to be 99.7% complete according to the Arthropoda busco gene space (For details see Supplementary Table 6). The genome is being published at NCBI under the BioProject (PRJNA733472).

### Genome annotation

We annotated protein-coding genes based on the genome sequence assembly of *C. gigas* (GCA013123115.1, ASM1312311v1. Repeats were soft-masked using RepeatMasker annotations (GCA013123115.1_ASM1312311v1_rm.out) with tabtk, bioawk and seqtk (https://github.com/lh3). We used Funannotate v1.8.1^62^ and Uniprot (sprot) for homology-based evidence based on protein sequences from 11 related bee species: *B. impatiens*: GCF000188095.2, *B. terrestris:* GCF000214255.1, *A. mellifera:* GCF003254395.2,*M. quadrifasciata:* GCA001276565.1, *E. mexicana:* GCF001483705.1, *F. varia* GCA011392965.1, *M. rotundata* GCF000220905.1, *H. laboriosa* GCF001263275.1, *D. novaeangliae* GCF001272555.1, *M. genalis* GCF011865705.1, *N. melanderi* GCF003710045.1. Briefly, funannotate used gene predictions from Genemark-ES, Snap v2006-07-28, glimmerHmm v3.0.4, Augustus v.3.3.3, and CodingQuarry v2.0 together with protein alignments in Evidence Modeler v.1.1.1. Too short, gap-spanning or repeat-overlapping gene models were removed (n = 5446) and tRNA genes were detected (n = 168) with tRNAscan-SE v2.0.6. Genes were functionally annotated using PFAM v33.1, the UniProt database v2018_11, EggNog (eggnog_4.5/hmmdb databases: Arthropoda, Insecta, Hymenoptera, Drosophila), MEROPS v12.0, CAZYmes in dbCAN v7.0, BUSCO Hymenoptera models v3.0.2, Hymenoptera odb9, SignalP v4.1, and InterProScan5 v81.0. The final annotation contained models for 20,016 protein-coding genes and 168 tRNAs, and was estimated to be 87.1% complete (BUSCO4 v4.1.4). The resulting gene annotation files for *C. gigas, E. dilemma, M. beecheii, T. carbonaria* and *Xylocopa violacea* are made available as **Additional Files 4-8** in the Zenodo archive accompanying this manuscript (10.5281/zenodo.5734574).

### Genomic microsynteny analysis

We traced abundant venom gland transcripts that potentially encoded toxins to homologs in the annotated, highly-continuous publicly-available genomes of bees (and wasps, ants, parasitoid wasps and sawflies as outgroup species) using the online BLAST suite against genomic databases. To identify conserved synteny blocks, we first identified the reciprocal best-match paralogs from hymenopteran all-against-all BLASTP comparisons of the venom genes. Based on the matching sequences, we then extracted exons from the candidate venom genes and their flanking genes. We used those to create local BLAST databases to survey the selected genomes using local tblastx with an e-value cutoff of 0.01. We then applied filters to select venom genes containing scaffolds at least 20 kbp in length (to exclude partial genes) with at least two exons. Where gene annotations were insufficient, we manually re-annotated venom genes where possible, following intron boundaries and using known sequences as templates. We extracted the coding sequences of all complete genes for phylogenetic analysis to establish ortholog groups in addition to their microsyntenic patterns. All resulting annotations are available as part of the Additional Materials (**Additional File 9).**

### Orthology prediction and phylogenetic analysis

All toxin transcripts together with toxin genes and their outgroup venom-unrelated homologs (e.g. trypsins and chymotrypsins in case of serine proteases) were arranged by gene family and aligned as translated amino acids using MAFFT^63^ (L-INS-I, 1000 iterations). Name convention was established to differentiate between genomic sequences (first two letters of both genus and species name, followed by the last three digits of a bioinformatic scaffold ID, followed – if applicable – by an abbreviation of a pre-existing gene annotation, followed by letters a to z to differentiate between sequences from the same scaffold); proteo-transcriptomic sequences (names kept the same as generated by transcriptome assemblers); homologues from UniProt and SwissProt databases used to provide outgroups and fill the gaps in sequence space (kept as UniProt or SwissProt IDs, but reduced to 10 characters if needed due to strict limitations of phylip format used by Exabayes). Alignments were manually inspected for overt errors (e.g., proper alignment of the cysteine backbone) and used to construct phylogenetic trees in Exabayes^64^ (four parallel runs of four chains each, runs stopped when average standard deviation of split frequencies of trees reached below 5%). Resulting trees are shown in the **Supplementary Figures 1-10**, with toxin sequences recovered from *Apis, Halictus* or *Xylocopa* venom marked as red arrows and non-toxic physiological sequences marked with grey arrow.

### A novel perspective on relations of short peptides: embedding space analysis

Every year, algorithms improve natural language processing (NLP) tasks such as automated translation or question answering, in particular by feeding large text corpora into Deep Learning (DL) based Language Models (LMs)^65^. These advances have been transferred to protein sequences by learning to predict masked or missing amino acids using large databases of raw protein sequences as input^66,67^. Such methods leverage the wealth of information present in exponentially growing unlabelled protein sequence databases by solely relying on sequential patterns found in the input. Processing the information learned by such protein LMs (pLMs), e.g., by feeding a protein sequence as input to the network and constructing vectors thereof from the activation in the network’s last layers, yields a representation of protein sequences referred to as embeddings^66^. This way, features learned by the pLM can be transferred to any (prediction) task requiring numerical protein representations (transfer learning) which has already been showcased for various aspects ranging from protein structure^68^ over protein function^69^. Further, it was shown that distance in embedding space correlates with protein function and can be used as an orthogonal signal for clustering proteins into functional families^69^.

Here, we used the pLM ProtT5-XL-UniRef5 0^66^ (in the following ProtT5) to create fixed-length vector representations for each protein sequence (per-protein embeddings) irrespective of its length. Towards this, we first created individual vector representations for each residue in a protein. In order to derive fixed-length vector representations for single proteins (per-protein embedding) irrespective of a protein’s length, we then averaged over all residue embeddings in a protein (Fig. 1 in Elnaggar et al.^66^). The protein Language Model (pLM) ProtT5 was trained solely on unlabelled protein sequences from BFD (Big Fantastic Database; 2.5 billion sequences including meta-genomic sequences)^70^ and UniRef50. ProtT5 has been built in analogy to the NLP (Natural Language Processing) T5^65^ ultimately learning some of the constraints of protein sequence. As ProtT5 was only trained on unlabelled protein sequences and no supervised training or fine-tuning was performed, there is no risk of information leakage or overfitting to a certain class or label. As a result, every protein was represented as 1024-dimensional per-protein embeddings. Those high-dimensional representations were projected to 3-d using UMAP (n_neighbors=10, min_dist=0.3, random_state=42, n_components=3) and coulored according to their taxonomic group to allow for visual analysis. Embeddings and 3-d plots were created using the bio_embeddings package^71^.

## Author contributions

BMvR and IK conceived the project and wrote the manuscript draft. Proteo-transcriptomic and genomic data were analysed by IK, MV and BMvR. Mass spectrometry was conducted by S.D, T.T. and G.L. Machine learning analysis was conducted by M.H. and B.R. New genome and annotation data were provided by E.S., R.G., B.A.H, BMvR. DNA extraction, library prep and assembly of *X. violacea* by C.G., A.B and DGP. All authors wrote the final manuscript version.

## Acknowledgements

BMvR and IK thank the German Science Foundation (DFG) for funding this work by a grant to BMvR (DFG RE3454/6-1). BMvR and IK thank Frank Förster for fruitful discussions on bioinformatics. BMvR is grateful to Georg Petschenka and Hermann Falkenhahn for helpful insights and discussion on solitary bees and localities in Giessen. BMvR thanks finally Ingo Ebersberger for his support. BMvR and AV acknowledge generous funding obtained by AV from the Hessian Ministry of Higher Education, Research, and the Arts for the group “Animal Venomics”, which was coordinated by BMvR until end of 2021 at the Institute for Insectbiotechnology at the Justus Liebig University, embedded in the LOEWE Centre for Translational Biodiversity Genomics. We thank the Genome Technology Center (RGTC) at Radboudumc for the use of the Sequencing Core Facility (Nijmegen, The Netherlands), which provided the PacBio SMRT sequencing service on the Sequel IIe platform. We finally are grateful to anonymous reviewers from previous manuscript versions for helpful comments.

## Supplementary Material

**Supplementary Table 1.** Proteo-transcriptomically identified venom components in *X. violacea*.

**Supplementary Table 2.** Proteo-transcriptomically identified venom components in *H. scabiosae*.

**Supplementary Table 3.** Proteo-transcriptomically identified venom components in *X. violacea*.

**Supplementary Table 4.** Listed bioactivity and description of the prevalent bee venom proteins.

**Supplementary Table 5.** List of mined and used high-quality hymenopteran genomes.

**Supplementary Table 6.** Resulting statistics of the new genome sequence of *X. violacea*.

**Supplementary Table 7.** BUSCO statistics for venom gland assemblies of *X. violacea, H. scabiosae, A. mellifera*.

**Supplementary Figure 1.** Phylogenetic tree of anthophilin1 peptides.

**Supplementary Figure 2.** Phylogenetic tree of melittin peptides.

**Supplementary Figure 3.** Phylogenetic tree of phospholipase A2 proteins.

**Supplementary Figure 4.** Phylogenetic tree of hyaluronidase proteins.

**Supplementary Figure 5.** Phylogenetic tree of icarapin proteins.

**Supplementary Figure 6.** Phylogenetic tree of dipeptidyl peptidase 4 proteins.

**Supplementary Figure 7.** Phylogenetic tree of acid phosphatase proteins.

**Supplementary Figure 8.** Phylogenetic tree of venom serine protease proteins.

**Supplementary Figure 9.** Phylogenetic tree of venom allergen proteins.

**Supplementary Figure 10.** Phylogenetic tree of secapin proteins.

**Supplementary File 1.** Alignment of anthophilin1 peptides.

**Supplementary File 2.** Alignment of melittin peptides.

**Supplementary File 3.** Alignment of “aculeatoxins” with signal peptide

**Supplementary File 4.** Alignment of only mature regions of “aculeatoxins**”**

**Supplementary File 5.** Machine learning results for “aculeatoxins” with signal peptide

**Supplementary File 6.** Machine learning results for mature region of “aculeatoxins”

**Supplementary File 7.** Alignment of phospholipase A2 proteins.

**Supplementary File 8.** Alignment of hyaluronidase proteins.

**Supplementary File 9.** Alignment of icarapin proteins.

**Supplementary File 10.** Alignment of dipeptidyl peptidase 4 proteins.

**Supplementary File 11.** Alignment of venom acid phosphatase proteins

**Supplementary File 12.** Alignment of venom serine protease proteins.

**Supplementary File 13.** Alignment of venom allergen proteins.

**Supplementary File 14.** Alignment of secapin proteins.

**Additional material** (DOI: 10.5281/zenodo.6998876).

**Additional File 1.** VG Assembly file of *Xylocopa violacea* following ORF prediction by Transdecoder.

**Additional File 2.** VG Assembly file of *Halictus scabiosae* following ORF prediction by Transdecoder.

**Additional File 3.** VG Assembly file of *Apis mellifera* following ORF prediction by Transdecoder.

**Additional File 4.** Genome annotation of *Colletes gigas*.

**Additional File 5.** Genome annotation of *Euglossa dilemma*.

**Additional File 6.** Genome annotation of *Melipona beecheii*.

**Additional File 7.** Genome annotation of *Tetragonula carbonaria*.

**Additional File 8.** Genome annotation of *Xylocopa violacea*.

**Additional File 9.** Gff files of all toxin gene annotations.

## References

1. Dunn, C. W. & Munro, C. Comparative genomics and the diversity of life. Zoologica Scripta 45, 5–13 (2016).

2. Yin, W. et al. Evolutionary trajectories of snake genes and genomes revealed by comparative analyses of five-pacer viper. Nature Communications 7, 13107 (2016).

3. Casewell, N. R., Wüster, W., Vonk, F. J., Harrison, R. A. & Fry, B. G. Complex cocktails: the evolutionary novelty of venoms. Trends in Ecology & Evolution 28, 219–229 (2013).

4. Drukewitz, S. H. & von Reumont, B. M. The Significance of Comparative Genomics in Modern Evolutionary Venomics. Front. Ecol. Evol. 7, 163 (2019).

5. Zancolli, G. & Casewell, N. R. Venom Systems as Models for Studying the Origin and Regulation of Evolutionary Novelties. Molecular Biology and Evolution 37, 2777–2790 (2020).

6. Jackson, T. N. W. & Koludarov, I. How the Toxin got its Toxicity. Frontiers in Pharmacology 11, 1893 (2020).

7. Almeida, D. D. et al. Tracking the recruitment and evolution of snake toxins using the evolutionary context provided by the Bothrops jararaca genome. Proc Natl Acad Sci USA 118, e2015159118 (2021).

8. Giorgianni, M. W. et al. The origin and diversification of a novel protein family in venomous snakes. Proc Natl Acad Sci USA 117, 10911–10920 (2020).

9. Moran, Y. et al. Concerted Evolution of Sea Anemone Neurotoxin Genes Is Revealed through Analysis of the Nematostella vectensis Genome. Molecular Biology and Evolution 25, 737–747 (2008).

10. Sachkova, M. Y. et al. The Birth and Death of Toxins with Distinct Functions: A Case Study in the Sea Anemone Nematostella. Molecular Biology and Evolution 36, 2001–2012 (2019).

11. Margres, M. J. et al. Quantity, Not Quality: Rapid Adaptation in a Polygenic Trait Proceeded Exclusively through Expression Differentiation. Molecular Biology and Evolution 34, 3099–3110 (2017).

12. von Reumont, B. M. et al. Modern venomics—Current insights, novel methods, and future perspectives in biological and applied animal venom research. GigaScience 11, giac048 (2022).

13. Holford, M., Daly, M., King, G. F. & Norton, R. S. Venoms to the rescue. Science 361, 842–844 (2018).

14. Oeyen, J. P. et al. Sawfly Genomes Reveal Evolutionary Acquisitions That Fostered the Mega-Radiation of Parasitoid and Eusocial Hymenoptera. Genome Biology and Evolution 12, 1099–1188 (2020).

15. Wang, T. et al. Proteomic analysis of the venom and venom sac of the woodwasp, Sirex noctilio - Towards understanding its biological impact. Journal of Proteomics 146, 195–206 (2016).

16. Piek, T. Venoms of the Hymenoptera. (Elsevier, 1986).

17. Danneels, E., Van Vaerenbergh, M., Debyser, G., Devreese, B. & de Graaf, D. Honeybee Venom Proteome Profile of Queens and Winter Bees as Determined by a Mass Spectrometric Approach. Toxins 7, 4468–4483 (2015).

18. Moreno, M. & Giralt, E. Three Valuable Peptides from Bee and Wasp Venoms for Therapeutic and Biotechnological Use: Melittin, Apamin and Mastoparan. Toxins 7, 1126–1150 (2015).

19. Walker, A. A. et al. Entomo-venomics - The evolution, biology and biochemistry of insect venoms. Toxicon 154, 15–27 (2018).

20. Walker, A. A., Robinson, S. D., Hamilton, B. F., Undheim, E. A. B. & King, G. F. Deadly Proteomes: A Practical Guide to Proteotranscriptomics of Animal Venoms. PROTEOMICS 20, 1900324 (2020).

21. von Reumont, B. M., Dutertre, S. & Koludarov, I. Venom profile of the European carpenter bee *Xylocopa violacea*: Evolutionary and applied considerations on its toxin components. Toxicon: X 14, 100117 (2022).

22. Lee, S., Baek, J. & Yoon, K. Differential Properties of Venom Peptides and Proteins in Solitary vs. Social Hunting Wasps. Toxins 8, 32 (2016).

23. dos Santos-Pinto, J. R. A., Perez-Riverol, A., Lasa, A. M. & Palma, M. S. Diversity of peptidic and proteinaceous toxins from social Hymenoptera venoms. Toxicon 148, 172–196 (2018).

24. Pucca, M. B. et al. Bee Updated: Current Knowledge on Bee Venom and Bee Envenoming Therapy. Front. Immunol. 10, 1–15 (2019).

25. Touchard, A. et al. Deciphering the Molecular Diversity of an Ant Venom Peptidome through a Venomics Approach. Journal of Proteome Research 17, 3503–3516 (2018).

26. Robinson, S. D. et al. A comprehensive portrait of the venom of the giant red bull ant, Myrmecia gulosa, reveals a hyperdiverse hymenopteran toxin gene family. Science advances 4, eaau4640 (2018).

27. Dashevsky, D. & Rodriguez, J. A Short Review of the Venoms and Toxins of Spider Wasps (Hymenoptera: Pompilidae). Toxins 13, 744 (2021).

28. Abd El-Wahed, A. et al. Wasp Venom Biochemical Components and Their Potential in Biological Applications and Nanotechnological Interventions. Toxins 13, 206 (2021).

29. Fry, B. G. et al. The structural and functional diversification of the Toxicofera reptile venom system. Toxicon 60, 434–448 (2012).

30. Burzyńska, M. & Piasecka-Kwiatkowska, D. A Review of Honeybee Venom Allergens and Allergenicity. Int J Mol Sci 22, 8371 (2021).

31. Peters, R. S. et al. Evolutionary History of the Hymenoptera. Current biology: CB 27, 1013–1018 (2017).

32. Reams, A. B. & Roth, J. R. Mechanisms of gene duplication and amplification. Cold Spring Harb Perspect Biol 7, a016592 (2015).

33. Światły-Błaszkiewicz, A. et al. The Effect of Bee Venom Peptides Melittin, Tertiapin, and Apamin on the Human Erythrocytes Ghosts: A Preliminary Study. Metabolites 10, 191 (2020).

34. Chen, J., Guan, S.-M., Sun, W. & Fu, H. Melittin, the Major Pain-Producing Substance of Bee Venom. Neurosci. Bull. 32, 265–272 (2016).

35. Choo, Y. M. et al. Molecular cloning and antimicrobial activity of bombolitin, a component of bumblebee Bombus ignitus venom. Comp Biochem Physiol B Biochem Mol Biol 156, 168–173 (2010).

36. Stöcklin, R. et al. Structural identification by mass spectrometry of a novel antimicrobial peptide from the venom of the solitary bee Osmia rufa (Hymenoptera: Megachilidae). Toxicon 55, 20–27 (2010).

37. Čujová, S., Bednárová, L., Slaninová, J., Straka, J. & Čeřovský, V. Interaction of a novel antimicrobial peptide isolated from the venom of solitary bee Colletes daviesanus with phospholipid vesicles and Escherichia coli cells. Journal of Peptide Science 20, 885–895 (2014).

38. Monincová, L. et al. Structure–activity study of macropin, a novel antimicrobial peptide from the venom of solitary bee Macropis fulvipes (Hymenoptera: Melittidae). Journal of Peptide Science 20, 375–384 (2014).

39. Kawakami, H. et al. Isolation of biologically active peptides from the venom of Japanese carpenter bee, Xylocopa appendiculata. Journal of Venomous Animals and Toxins including Tropical Diseases 23, 29 (2017).

40. Sun, C. et al. Genus-Wide Characterization of Bumblebee Genomes Provides Insights into Their Evolution and Variation in Ecological and Behavioral Traits. Molecular Biology and Evolution 38, 486–501 (2021).

41. Fields, C. & Levin, M. Competency in Navigating Arbitrary Spaces as an Invariant for Analyzing Cognition in Diverse Embodiments. Entropy 24, 819 (2022).

42. Martinson, E. O., Mrinalini, Kelkar, Y. D., Chang, C.-H. & Werren, J. H. The Evolution of Venom by Cooption of Single-Copy Genes. Current biology: CB 27, 2007–2013.e8 (2017).

43. Dowell, N. L. et al. Extremely Divergent Haplotypes in Two Toxin Gene Complexes Encode Alternative Venom Types within Rattlesnake Species. Current Biology 28, 1016–1026.e4 (2018).

44. Koludarov, I. et al. Reconstructing the evolutionary history of a functionally diverse gene family reveals complexity at the genetic origins of novelty. 583344 Preprint at https://doi.org/10.1101/583344 (2020).

45. Danneels, E. L., Rivers, D. B. & de Graaf, D. C. Venom proteins of the parasitoid wasp Nasonia vitripennis: recent discovery of an untapped pharmacopee. Toxins 2, 494–516 (2010).

46. Choo, Y. M. et al. Dual Function of a Bee Venom Serine Protease: Prophenoloxidase-Activating Factor in Arthropods and Fibrin(ogen)olytic Enzyme in Mammals. PLoS ONE 5, (2010).

47. Hoffman, D. R. Hymenoptera Venom Allergens. Clinical Reviews in Allergy & Immunology 30, 109–128 (2006).

48. Fry, B. G. et al. The toxicogenomic multiverse: convergent recruitment of proteins into animal venoms. Annual Review of Genomics and Human Genetics 10, 483–511 (2009).

49. Jackson, T. N. W. et al. Rapid Radiations and the Race to Redundancy: An Investigation of the Evolution of Australian Elapid Snake Venoms. Toxins (Basel) 8, 309 (2016).

50. Grandal, M., Hoggard, M., Neely, B., Davis, W. C. & Marí, F. Proteogenomic Assessment of Intraspecific Venom Variability: Molecular Adaptations in the Venom Arsenal of Conus purpurascens. Mol Cell Proteomics 20, 100100 (2021).

51. Dowell, N. L., Giorgianni, M. W., Kassner, V. A. & Selegue, J. E. The Deep Origin and Recent Loss of Venom Toxin Genes in Rattlesnakes. Current Biology (2016).

52. Elieh Ali Komi, D., Shafaghat, F. & Zwiener, R. D. Immunology of Bee Venom. Clinic Rev Allerg Immunol 54, 386–396 (2018).

53. MacManes, M. D. The Oyster River Protocol: a multi-assembler and kmer approach for de novo transcriptome assembly. PeerJ 6, e5428 (2018).

54. Bray, N. L., Pimentel, H., Melsted, P. & Pachter, L. Near-optimal probabilistic RNA-seq quantification. Nature Biotechnology 34, 525–527 (2016).

55. Miller, S. A., Dykes, D. D. & Polesky, H. F. A simple salting out procedure for extracting DNA from human nucleated cells. Nucleic Acids Res 16, 1215 (1988).

56. Cheng, H., Concepcion, G. T., Feng, X., Zhang, H. & Li, H. Haplotype-resolved de novo assembly using phased assembly graphs with hifiasm. Nat Methods 18, 170–175 (2021).

57. Ludwig, A., Pippel, M., Myers, G. & Hiller, M. DENTIST—using long reads for closing assembly gaps at high accuracy. GigaScience 11, giab100 (2022).

58. Langmead, B. & Salzberg, S. L. Fast gapped-read alignment with Bowtie 2. Nat Methods 9, 357–359 (2012).

59. Poplin, R. et al. A universal SNP and small-indel variant caller using deep neural networks. Nat Biotechnol 36, 983–987 (2018).

60. Li, H. A statistical framework for SNP calling, mutation discovery, association mapping and population genetical parameter estimation from sequencing data. Bioinformatics 27, 2987–2993 (2011).

61. Laetsch, D. R. & Blaxter, M. L. BlobTools: Interrogation of genome assemblies. F1000Res 6, 1287 (2017).

62. Palmer, J. M. & Stajich, J. Funannotate v1.8.1: Eukaryotic genome annotation. (2020) doi:10.5281/zenodo.4054262.

63. Katoh, K. & Standley, D. M. MAFFT Multiple Sequence Alignment Software Version 7: Improvements in Performance and Usability. Molecular Biology and Evolution 30, 772–780 (2013).

64. Aberer, A. J., Kobert, K. & Stamatakis, A. ExaBayes: Massively Parallel Bayesian Tree Inference for the Whole-Genome Era. Molecular Biology and Evolution 31, 2553–2556 (2014).

65. Raffel, C. et al. Exploring the Limits of Transfer Learning with a Unified Text-to-Text Transformer. Journal of Machine Learning Research 21, 1–67 (2020).

66. Elnaggar, A. et al. ProtTrans: Towards Cracking the Language of Lifes Code Through Self-Supervised Deep Learning and High Performance Computing. IEEE Transactions on Pattern Analysis and Machine Intelligence 1–1 (2021) doi:10.1109/TPAMI.2021.3095381.

67. Rives, A. et al. Biological structure and function emerge from scaling unsupervised learning to 250 million protein sequences. Proceedings of the National Academy of Sciences 118, e2016239118 (2021).

68. Weißenow, K., Heinzinger, M. & Rost, B. Protein language model embeddings for fast, accurate, alignment-free protein structure prediction. 2021.07.31.454572 Preprint at https://doi.org/10.1101/2021.07.31.454572 (2021).

69. Littmann, M., Heinzinger, M., Dallago, C., Olenyi, T. & Rost, B. Embeddings from deep learning transfer GO annotations beyond homology. Sci Rep 11, 1160 (2021).

70. Steinegger, M., Mirdita, M. & Söding, J. Protein-level assembly increases protein sequence recovery from metagenomic samples manyfold. Nat Methods 16, 603–606 (2019).

71. Dallago, C. et al. Learned Embeddings from Deep Learning to Visualize and Predict Protein Sets. Current Protocols 1, e113 (2021).

