## Supplementary Figure 2 for "A common venomous ancestor? Prevalent bee venom genes evolved before the aculeate stinger while few major toxins are bee-specific"

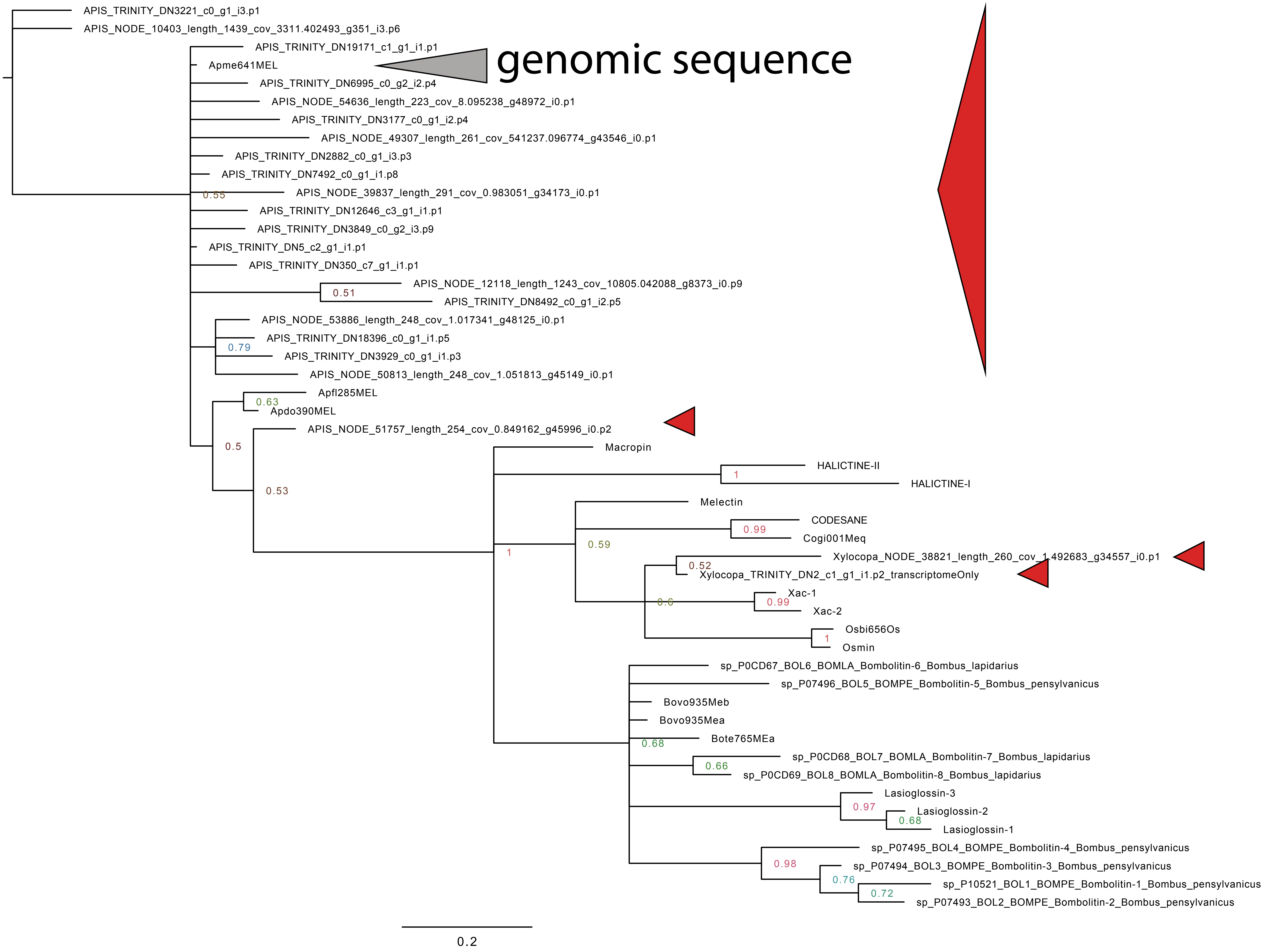


**Phylogenetic tree of the Melittin protein family.** Red arrows mark those that were recovered from the transcriptomes of *X. violacea*, *H. scabiosae* and *A. mellifera* in the present study. Genomic sequences recovered in the present study have naming convention of *Gesp###PPa(U)* where *Ge* stands for first two letters of the genus name, *sp* stands for first two letters of the species name, *###* stands for the last three digits of the genomic scaffold ID, *PP* stands for the protein label and *a* stands for A to Z identifier given to homologous genes if several were found on the same continuous genomic scaffold. Where genomic and transcriptomic sequences were identical, we kept transcriptomic sequence. UniProt and GeneBank IDs were kept in their original form. Forms known from proteomes only were kept with the names given in the original studies.
