## Supplementary Figure 6 for "A common venomous ancestor? Prevalent bee venom genes evolved before the aculeate stinger while few major toxins are bee-specific"

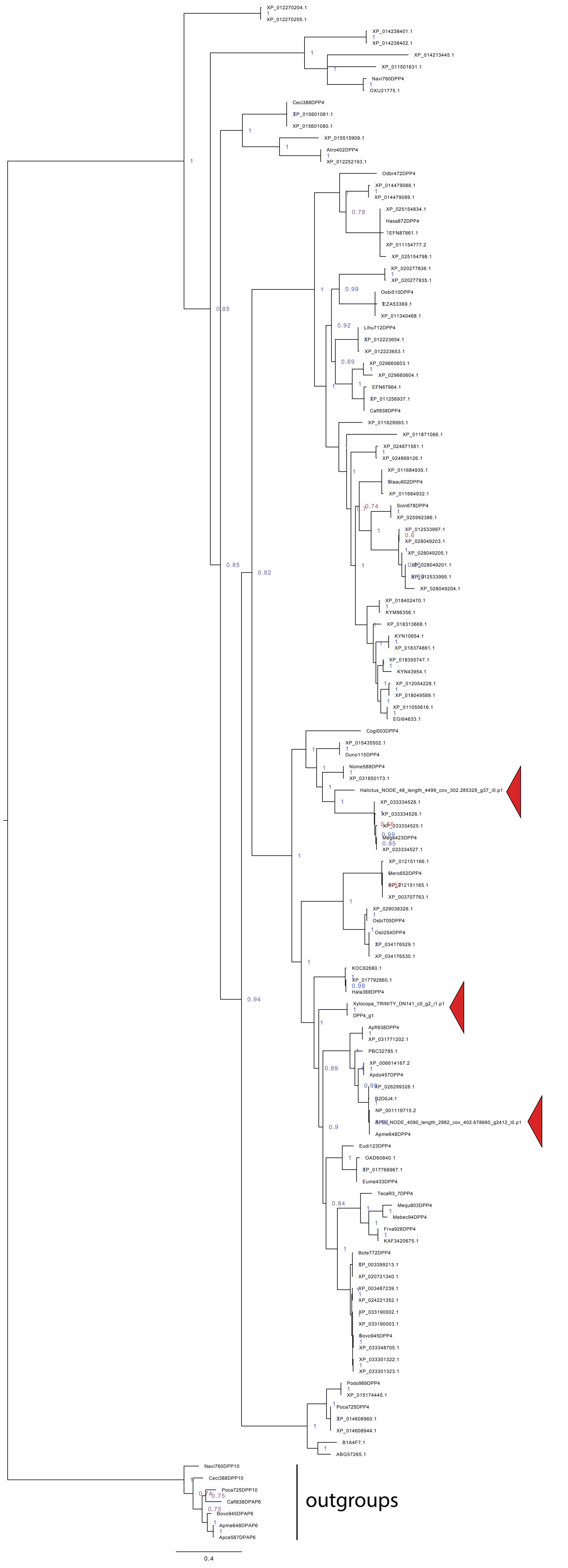


**Phylogenetic tree of the Dipeptidyl peptidase-4 protein family.** Red arrows mark those that were recovered from the transcriptomes of *X. violacea*, *H. scabiosae* and *A. mellifera* in the present study. Genomic sequences recovered in the present study have naming convention of *Gesp###PPPP##* where *Ge* stands for first two letters of the genus name, *sp* stands for first two letters of the species name, *###* stands for the last three digits of the genomic scaffold ID, *PPPP##* stands for the protein label. Where genomic and transcriptomic sequences were identical, we kept transcriptomic sequence. UniProt and GeneBank IDs were kept in their original form.
