## Supplementary Table 1 for "A common venomous ancestor? Prevalent bee venom genes evolved before the aculeate stinger while few major toxins are bee-specific"

*Apis mellifera*

**Transcriptome data analysis**

| Transdecoder protein IDs | Transcript ID | Expression value | Blast-P toxprot (sseqid) | E-value | Bitscore | Blast-P UniProt Apocrita (sseqid) | E-value | Bitscore | Blast-P UniProt All (sseqid) | E-value | Bitscore | Physiological association | Venom protein family |
|---|---|---|---|---|---|---|---|---|---|---|---|---|---|
| J1024640.p1 | J1024640 | 28.98183226 | NA | NA | NA | NA | NA | NA | Uncharacterized protein/V-type proton ATPase subuni | 1.60E-164 | 1,203 | No venom function | NA |
| NODE_10174_length_1471_cov_85.495702_g6621_i0.p1 | NODE_10174_length_1471_cov_85.495702_g6621_i0 | 15.13873563 | Blarina toxin | 2.19E-28 | 104 | Chymotrypsin-1 | 4.74E-30 | 105 | Peptidase S1 domain-containing protein | 5.37E-163 | 455 | Toxin | Peptidase S1 domain (Venom Serine Protease) |
| NODE_10403_length_1439_cov_3311.402493_g351_i3.p6 | NODE_10403_length_1439_cov_3311.402493_g351_i3 | 12.32147836 | Melittin | 4.2E-12 | 51.6 | Melittin | 4.46E-13 | 51.6 | Melittin | 3.59E-09 | 52.4 | Toxin | Mellitin |
| NODE_10530_length_1432_cov_7.620915_g7215_i0.p1 | NODE_10530_length_1432_cov_7.620915_g7215_i0 | 3.519486718 | Venom serine protease 34 | 0 | 828 | Venom serine protease 34 | 0 | 828 | Venom serine protease 34 | 0 | 828 | Toxin | Peptidase S1 domain (Venom Serine Protease) |
| NODE_11040_length_1349_cov_3010.901099_g4121_i3.p1 | NODE_11040_length_1349_cov_3010.901099_g4121_i3 | 1607.576301 | Phospholipase A2 | 2.42E-127 | 350 | Phospholipase A2 | 2.57E-128 | 350 | Phospholipase A2 | 5.4E-124 | 350 | Toxin | Phospholipase A2 |
| NODE_11279_length_1337_cov_237.078783_g5817_i1.p1 | NODE_11279_length_1337_cov_237.078783_g5817_i1 | 116.3440908 | NA | NA | NA | NA | NA | NA | Uncharacterized protein/Protein CREG1 | 4.31E-43 | 150 | No venom function | NA |
| NODE_11920_length_1264_cov_21752.067825_g8218_i0.p1 | NODE_11920_length_1264_cov_21752.067825_g8218_i0 | 10874.7012 | Mast cell degranulating peptide | 6.36E-32 | 100 | Mast cell degranulating peptide | 6.75E-33 | 100 | Mast cell degranulating peptide | 5.63E-29 | 101 | Toxin | Mast cell degranulating |
| NODE_11920_length_1264_cov_21752.067825_g8218_i0.p3 | NODE_11920_length_1264_cov_21752.067825_g8218_i0 | 10874.7012 | Apamin | 2.14E-30 | 96.7 | Apamin | 2.27E-31 | 96.7 | Apamin protein | 4.77E-27 | 96.7 | Toxin | Apamin |
| NODE_12118_length_1243_cov_10805.042088_g8373_i0.p9 | NODE_12118_length_1243_cov_10805.042088_g8373_i0 | 5.837841716 | Melittin | 1.23E-07 | 38.9 | Melittin | 1.31E-08 | 38.9 | Melittin | 0.000272 | 38.5 | Toxin | Mellitin |
| NODE_13401_length_1105_cov_468.260000_g9314_i0.p1 | NODE_13401_length_1105_cov_468.260000_g9314_i0 | 2.385192673 | Peroxiredoxin-4 (Fragments) | 8.72E-18 | 68.9 | NA | NA | NA | Thioredoxin domain-containing protein | 3.38E-109 | 314 | Venom component | Peroxiredoxin4 |
| NODE_143_length_9096_cov_32.937286_g93_i0.p1 | NODE_143_length_9096_cov_32.937286_g93_i0 | 11.20478868 | NA | NA | NA | NA | NA | NA | Anion exchange protein | 0 | 2126 | No venom function | NA |
| NODE_14318_length_1026_cov_588.271885_g10016_i0.p1 | NODE_14318_length_1026_cov_588.271885_g10016_i0 | 308.5600124 | NA | NA | NA | NA | NA | NA | Superoxide dismutase [Cu-Zn] | 5.76E-106 | 304 | Venom component (parasitoids) | Superoxide dismutase |
| NODE_14549_length_987_cov_148.449561_g9868_i0.p1 | NODE_14549_length_987_cov_148.449561_g9868_i0 | 98.59614894 | NA | NA | NA | NA | NA | NA | Peptidyl-prolyl cis-trans isomerase | 1.82E-147 | 413 | No venom function | NA |
| NODE_14992_length_970_cov_72249.791257_g6125_i4.p2 | NODE_14992_length_970_cov_72249.791257_g6125_i4 | 25687.32858 | NA | NA | NA | NA | NA | NA | Uncharacterized protein (with signalpeptide) | 3.86E-67 | 201 | Unknown function | NA |
| NODE_15131_length_938_cov_131.254925_g10349_i0.p2 | NODE_15131_length_938_cov_131.254925_g10349_i0 | 88.59293273 | NA | NA | NA | Histone H4 | 1.44E-70 | 199 | Histone H4 (Fragment) | 9.39E-67 | 201 | No venom function | NA |
| NODE_15441_length_913_cov_534.093079_g10606_i0.p1 | NODE_15441_length_913_cov_534.093079_g10606_i0 | 349.3200055 | Peroxiredoxin-4 (Fragments) | 4.9E-17 | 67.8 | NA | NA | NA | Thioredoxin domain-containing protein | 2.52E-143 | 402 | Venom component | Peroxiredoxin4 |
| NODE_1577_length_4346_cov_815.361920_g1019_i0.p1 | NODE_1577_length_4346_cov_815.361920_g1019_i0 | 10.55190026 | NA | NA | NA | NA | NA | NA | Uncharacterized protein/Actin | 0 | 784 | No venom function | NA |
| NODE_16160_length_860_cov_4556.777070_g11199_i0.p1 | NODE_16160_length_860_cov_4556.777070_g11199_i0 | 17.80246022 | Secapin | 3.9E-53 | 156 | Secapin | 4.14E-54 | 156 | Secapin | 8.68E-50 | 156 | Toxin | Secapin |
| NODE_18028_length_766_cov_1892.343179_g8026_i1.p1 | NODE_18028_length_766_cov_1892.343179_g8026_i1 | 1057.334334 | Omega-conotoxin-like protein 1 | 1.36E-52 | 154 | Omega-conotoxin-like protein 1 | 1.44E-53 | 154 | Inhibitor cysteine knot peptide | 3.02E-49 | 154 | Toxin | ICK-Knottin |
| NODE_18339_length_747_cov_15153.239884_g8218_i2.p1 | NODE_18339_length_747_cov_15153.239884_g8218_i2 | 168.368931 | Mast cell degranulating peptide | 6.36E-32 | 100 | Mast cell degranulating peptide | 6.75E-33 | 100 | Mast cell degranulating peptide | 5.63E-29 | 101 | Toxin | Mast cell degranulating |
| NODE_19733_length_678_cov_176.370787_g14498_i0.p1 | NODE_19733_length_678_cov_176.370787_g14498_i0 | 108.5329073 | NA | NA | NA | Histone H4 | 1.44E-70 | 199 | Histone H4 (Fragment) | 9.39E-67 | 201 | No venom function | NA |
| NODE_20261_length_635_cov_63.437500_g14817_i0.p1 | NODE_20261_length_635_cov_63.437500_g14817_i0 | 47.3117786 | NA | NA | NA | NA | NA | NA | Odorant binding protein OBP15 | 8.65E-90 | 263 | No venom function | NA |
| NODE_20389_length_648_cov_68298.664418_g15085_i0.p1 | NODE_20389_length_648_cov_68298.664418_g15085_i0 | 19.47422899 | NA | NA | NA | NA | NA | NA | Odorant binding protein OBP21 | 7.21E-92 | 267 | No venom function | NA |
| NODE_2447_length_3705_cov_272.814050_g1430_i0.p1 | NODE_2447_length_3705_cov_272.814050_g1430_i0 | 142.7420516 | Calglandulin | 3E-40 | 129 | NA | NA | NA | Putative calmodulin isoform x3 (Fragment) | 2.92E-105 | 305 | Venom component | Calglandulin |
| NODE_24540_length_505_cov_3301.937778_g18995_i0.p1 | NODE_24540_length_505_cov_3301.937778_g18995_i0 | 1934.144672 | NA | NA | NA | NA | NA | NA | C1q-like venom protein | 7.5E-115 | 328 | Venom component | C1q-like |
| NODE_2528_length_3602_cov_40.640541_g1631_i0.p30 | NODE_2528_length_3602_cov_40.640541_g1631_i0 | 3.633832509 | NA | NA | NA | NA | NA | NA | Uncharacterized/Mellitin | 2.50E-08 | 125 | Toxin | Mellitin |
| NODE_27106_length_438_cov_721.168044_g21398_i0.p1 | NODE_27106_length_438_cov_721.168044_g21398_i0 | 266.1292996 | NA | NA | NA | NA | NA | NA | PVF1 variant 1 (Fragment) | 5.22E-49 | 161 | Venom component | PVF1 |
| NODE_28629_length_412_cov_2765.735905_g22893_i0.p3 | NODE_28629_length_412_cov_2765.735905_g22893_i0 | 927.4201223 | NA | NA | NA | Apidaecins type 73 (Fragment) | 4.56E-74 | 216 | Apidaecin (Fragment) | 1.63E-71 | 217 | Venom component, AMP? | Apidaecin |
| NODE_3011_length_3324_cov_1067.924748_g1019_i1.p1 | NODE_3011_length_3324_cov_1067.924748_g1019_i1 | 79.88520308 | NA | NA | NA | NA | NA | NA | Uncharacterized protein/Actin | 0 | 784 | No venom function | NA |
| NODE_3142_length_3280_cov_453.149766_g1714_i1.p1 | NODE_3142_length_3280_cov_453.149766_g1714_i1 | 245.4757559 | NA | NA | NA | NA | NA | NA | Uncharacterized protein/Isoform Sap-mu-6 | 8.10E-45 | 173 | No venom function | NA |
| NODE_3646_length_3065_cov_183.230769_g1489_i3.p1 | NODE_3646_length_3065_cov_183.230769_g1489_i3 | 67.82802985 | NA | NA | NA | Major royal jelly protein 1 | 5.76E-169 | 474 | Major royal jelly protein 9 | 0 | 867 | Venom component | Major Royal Jelly |
| NODE_39837_length_291_cov_0.983051_g34173_i0.p1 | NODE_39837_length_291_cov_0.983051_g34173_i0 | 2.635374381 | Melittin | 3.92E-22 | 77.4 | Melittin | 4.16E-23 | 77.4 | Melittin | 8.72E-19 | 77.4 | Toxin | Mellitin |
| NODE_40674_length_288_cov_0.824034_g35010_i0.p2 | NODE_40674_length_288_cov_0.824034_g35010_i0 | 1.34689755 | Phospholipase A2 | 1.04E-36 | 116 | Phospholipase A2 | 1.1E-37 | 116 | Phospholipase A2 | 2.31E-33 | 116 | Toxin | Phospholipase A2 |
| NODE_4090_length_2882_cov_402.678660_g2412_i0.p1 | NODE_4090_length_2882_cov_402.678660_g2412_i0 | 171.858226 | Venom dipeptidyl peptidase 4 | 0 | 1600 | Venom dipeptidyl peptidase 4 | 0 | 1600 | Venom dipeptidyl peptidase 4 | 0 | 1614 | Toxin | Venom dipeptidyl peptidase 4 |
| NODE_4142_length_2818_cov_303.087948_g453_i1.p1 | NODE_4142_length_2818_cov_303.087948_g453_i1 | 52.75760505 | NA | NA | NA | NA | NA | NA | Heat shock protein beta-1 | 2.62E-168 | 468 | No venom function | NA |
| NODE_4236_length_2786_cov_241.753936_g2779_i0.p1 | NODE_4236_length_2786_cov_241.753936_g2779_i0 | 11.23584173 | NA | NA | NA | NA | NA | NA | Uncharacterized protein (with signalpeptide) | 8.49E-68 | 203 | Unknown function | NA |
| NODE_44489_length_278_cov_0.748768_g38728_i0.p1 | NODE_44489_length_278_cov_0.748768_g38728_i0 | 0.727234945 | Icarapin | 2E-55 | 167 | Icarapin-like | 2.12E-56 | 167 | Icarapin variant 2 (Fragment) | 4.16E-55 | 174 | Toxin | Icarapin |
| NODE_49307_length_261_cov_541237.096774_g43546_i0.p1 | NODE_49307_length_261_cov_541237.096774_g43546_i0 | 9.628469185 | Melittin | 7.65E-11 | 49.3 | Melittin | 8.12E-12 | 49.3 | Melittin | 8.83E-08 | 50.1 | Toxin | Mellitin |
| NODE_50813_length_248_cov_1.051813_g45149_i0.p1 | NODE_50813_length_248_cov_1.051813_g45149_i0 | 4.689105126 | Melittin | 2.56E-18 | 67.4 | Melittin | 2.72E-19 | 67.4 | Melittin | 5.7E-15 | 67.4 | Toxin | Mellitin |
| NODE_5128_length_2490_cov_1011.854620_g3398_i0.p1 | NODE_5128_length_2490_cov_1011.854620_g3398_i0 | 264.9695056 | NA | NA | NA | NA | NA | NA | Chitinase protein Idgf4 | 2.95E-165 | 468 | No venom function | Chitinase |
| NODE_51757_length_254_cov_0.849162_g45996_i0.p2 | NODE_51757_length_254_cov_0.849162_g45996_i0 | 62.31264808 | Melittin | 1.53E-16 | 62.8 | Melittin | 1.62E-17 | 62.8 | Melittin | 3.4E-13 | 62.8 | Toxin | Mellitin |
| NODE_53677_length_249_cov_0.873563_g47916_i0.p1 | NODE_53677_length_249_cov_0.873563_g47916_i0 | 0.929055166 | Phospholipase A2 | 5.38E-28 | 94 | Phospholipase A2 | 5.71E-29 | 94 | Phospholipase A2 (Fragment) | 6.86E-25 | 93.6 | Toxin | Phospholipase A2 |
| NODE_53886_length_248_cov_1.017341_g48125_i0.p1 | NODE_53886_length_248_cov_1.017341_g48125_i0 | 17.14304722 | Melittin | 2.83E-20 | 72.4 | Melittin | 3E-21 | 72.4 | Melittin | 6.3E-17 | 72.4 | Toxin | Mellitin |
| NODE_54636_length_223_cov_8.095238_g48972_i0.p1 | NODE_54636_length_223_cov_8.095238_g48972_i0 | 13.22701702 | Melittin | 3.35E-22 | 77.8 | Melittin | 3.55E-23 | 77.8 | Melittin | 7.45E-19 | 77.8 | Toxin | Mellitin |
| NODE_6700_length_2072_cov_10229.525037_g4464_i1.p1 | NODE_6700_length_2072_cov_10229.525037_g4464_i1 | 2847.929228 | Icarapin | 5.07E-163 | 445 | Icarapin-like | 5.38E-164 | 445 | Icarapin-like | 1.13E-159 | 445 | Toxin | Icarapin |
| NODE_6769_length_2084_cov_39.210553_g4191_i0.p7 | NODE_6769_length_2084_cov_39.210553_g4191_i0 | 12.06708042 | NA | NA | NA | NA | NA | NA | ML domain-containing protein | 1.28E-23 | 90.9 | No venom function | NA |
| NODE_6915_length_2046_cov_279.860984_g4286_i0.p1 | NODE_6915_length_2046_cov_279.860984_g4286_i0 | 140.3027714 | NA | NA | NA | NA | NA | NA | Uncharacterized protein (with signalpeptide) | 8.49E-68 | 203 | Unknown function | NA |
| NODE_7578_length_1914_cov_211.404024_g4758_i0.p1 | NODE_7578_length_1914_cov_211.404024_g4758_i0 | 123.9178495 | NA | NA | NA | NA | NA | NA | Enolase | 0 | 829 | No venom function | Enolase |
| NODE_7698_length_1875_cov_36.041209_g4442_i1.p3 | NODE_7698_length_1875_cov_36.041209_g4442_i1 | 3.059739947 | NA | NA | NA | NA | NA | NA | ML domain-containing protein | 1.28E-23 | 90.9 | No venom function | NA |
| NODE_7890_length_1840_cov_98.634734_g5077_i1.p1 | NODE_7890_length_1840_cov_98.634734_g5077_i1 | 40.02176792 | NA | NA | NA | NA | NA | NA | Uncharacterized protein/Carboxypeptidase | 2.74E-172 | 497 | Venom component | Carboxypeptidase |
| NODE_7908_length_1851_cov_1172.670045_g4121_i1.p4 | NODE_7908_length_1851_cov_1172.670045_g4121_i1 | 78.11391077 | Phospholipase A2 | 8.94E-34 | 109 | Phospholipase A2 | 9.48E-35 | 109 | Phospholipase A2 | 1.1E-30 | 110 | Toxin | Phospholipase A2 |
| NODE_7958_length_1825_cov_1940.759322_g5357_i0.p1 | NODE_7958_length_1825_cov_1940.759322_g5357_i0 | 811.7356109 | NA | NA | NA | NA | NA | NA | Transferrin | 0 | 1252 | No venom function | NA |
| NODE_827_length_5603_cov_227.112112_g520_i1.p1 | NODE_827_length_5603_cov_227.112112_g520_i1 | 94.17465483 | NA | NA | NA | NA | NA | NA | Phosphotransferase | 0 | 921 | No venom function | NA |
| NODE_8360_length_1756_cov_674.852440_g5640_i0.p1 | NODE_8360_length_1756_cov_674.852440_g5640_i0 | 5.44999885 | Allergen Api m 6.03 / Api m 6.04 | 4.03E-36 | 114 | Allergen Api m 6.03 / Api m 6.04 | 4.28E-37 | 114 | Allergen Api m 6.03 / Api m 6.04 | 8.97E-33 | 114 | Toxin | Allergens |
| NODE_8622_length_1713_cov_138.656212_g588_i1.p1 | NODE_8622_length_1713_cov_138.656212_g588_i1 | 27.75089414 | NA | NA | NA | NA | NA | NA | Carboxypeptidase D | 0 | 943 | Venom component | CarboxypeptidaseD |
| NODE_9018_length_1656_cov_105.966477_g5751_i0.p1 | NODE_9018_length_1656_cov_105.966477_g5751_i0 | 59.14403364 | NA | NA | NA | NA | NA | NA | Malate dehydrogenase | 0 | 682 | No venom function | NA |
| NODE_9154_length_1625_cov_145.924204_g588_i2.p1 | NODE_9154_length_1625_cov_145.924204_g588_i2 | 28.58326591 | NA | NA | NA | NA | NA | NA | Carboxypeptidase D | 0 | 886 | Venom component | CarboxypeptidaseD |
| NODE_9157_length_1632_cov_16.547848_g5858_i0.p1 | NODE_9157_length_1632_cov_16.547848_g5858_i0 | 1.86670295 | Venom acid phosphatase Acph-1 | 0 | 800 | Venom acid phosphatase Acph-1 | 0 | 800 | Venom acid phosphatase Acph-1 | 0 | 826 | Toxin | Venom acid phosphatase Acph-1 |
| R1024725.p1 | R1024725 | 177.3056982 | Hyaluronidase | 0 | 691 | Hyaluronidase | 0 | 691 | Hyaluronidase | 0 | 691 | Toxin | Hyaluronidase |
| R1024865.p1 | R1024865 | 103.0537267 | Snake venom 5'-nucleotidase | 2.25E-139 | 414 | NA | NA | NA | Uncharacterized protein | 0 | 1222 | Venom component | Snake venom 5'-nucleotidase |
| R1025397.p1 | R1025397 | 235.814867 | NA | NA | NA | Major royal jelly protein 1 | 0 | 519 | Major royal jelly protein 8 | 0 | 854 | Venom component | Major Royal Jelly |
| R1025449.p1 | R1025449 | 114.4219697 | NA | NA | NA | NA | NA | NA | Aldose 1-epimerase | 0 | 783 | No venom function | NA |
| R1025530.p1 | R1025530 | 87.58483805 | Cystatin-1 | 1.71E-10 | 56.2 | NA | NA | NA | Uncharacterized protein | 0 | 1817 | Venom component | Cystatin |
| R1025617.p1 | R1025617 | 160.7505809 | Venom serine protease Bi-VSP | 7.6E-171 | 476 | Venom serine protease Bi-VSP | 8.06E-172 | 476 | Peptidase S1 domain-containing protein | 0 | 733 | Toxin | Peptidase S1 domain (Venom Serine Protease) |
| R1027063.p1 | R1027063 | 61.62505129 | NA | NA | NA | NA | NA | NA | Chitin-binding type-2 domain-containing protein | 0 | 1113 | No venom function | Chitin-binding |
| R1027523.p1 | R1027523 | 25.37254108 | Snake venom serine protease BITS01A | 4.11E-30 | 110 | Chymotrypsin-1 | 3.39E-32 | 112 | Peptidase S1 domain-containing protein | 0 | 540 | Toxin | Peptidase S1 domain (Venom Serine Protease) |
| R1028514.p1 | R1028514 | 69.69317471 | NA | NA | NA | NA | NA | NA | Malate dehydrogenase | 0 | 683 | No venom function | NA |
| R1029348.p1 | R1029348 | 63.79385514 | NA | NA | NA | NA | NA | NA | DnaJ subfamily B member | 0 | 764 | No venom function | NA |
| R1036651.p1 | R1036651 | 949.9860718 | Phospholipase A2 | 3.5E-49 | 156 | Phospholipase A2 | 3.71E-50 | 156 | PA2c domain-containing protein | 5.07E-127 | 362 | Toxin | PhospholipaseA2 |
| S1036426.p2 | S1036426 | 9.331919492 | NA | NA | NA | NA | NA | NA | LYS1 (Fragment) | 1.3E-62 | 192 | No venom function | NA |
| S1045097.p1 | S1045097 | 17.30434736 | Reticulocalbin-2 | 4.4E-60 | 191 | NA | NA | NA | Uncharacterized protein/Calumenin | 0 | 1,543 | No venom function | NA |
| S1047992.p1 | S1047992 | 24.05216591 | NA | NA | NA | NA | NA | NA | Gal_mutarotas_2 domain-containing protein | 0 | 1919 | No venom function | NA |
| TRINITY_DN10322_c0_g1_i2.p6 | TRINITY_DN10322_c0_g1_i2 | 0 | Melittin | 8.11E-08 | 39.3 | Melittin | 8.6E-09 | 39.3 | Melittin | 0.00018 | 39.3 | Toxin | Mellitin |
| TRINITY_DN1093_c0_g3_i2.p1 | TRINITY_DN1093_c0_g3_i2 | 179.524656 | NA | NA | NA | NA | NA | NA | Superoxide dismutase [Cu-Zn] | 9.92E-124 | 351 | Venom component (parasitoids) | Superoxide dismutase |
| TRINITY_DN11539_c0_g1_i7.p1 | TRINITY_DN11539_c0_g1_i7 | 39.166806 | Venom carboxylesterase-6 | 0 | 724 | Venom carboxylesterase-6 | 0 | 724 | Carboxylic ester hydrolase | 0 | 724 | Venom component | Venom carboxylesterase-6 |
| TRINITY_DN11539_c0_g1_i9.p2 | TRINITY_DN11539_c0_g1_i9 | 44.07062859 | Venom carboxylesterase-6 | 1.37E-105 | 308 | Venom carboxylesterase-6 | 1.46E-106 | 308 | Venom carboxylesterase-6 | 3.06E-102 | 308 | Toxin | Venom carboxylesterase-6 |
| TRINITY_DN1249_c0_g1_i1.p11 | TRINITY_DN1249_c0_g1_i1 | 18.00976628 | NA | NA | NA | NA | NA | NA | Uncharacterized/Mellitin | 2.30E-17 | 188 | Toxin | Mellitin |
| TRINITY_DN12646_c3_g1_i1.p1 | TRINITY_DN12646_c3_g1_i1 | 12.93459961 | Melittin | 4.39E-15 | 59.3 | Melittin | 4.66E-16 | 59.3 | Melittin | 9.77E-12 | 59.3 | Toxin | Mellitin |
| TRINITY_DN1306_c0_g1_i1.p1 | TRINITY_DN1306_c0_g1_i1 | 102.3583593 | NA | NA | NA | NA | NA | NA | Glucose-regulated protein | 0 | 1332 | No venom function | NA |
| TRINITY_DN14348_c0_g2_i1.p1 | TRINITY_DN14348_c0_g2_i1 | 3.510212621 | NA | NA | NA | NA | NA | NA | Uncharacterized protein/Glucose dehydrogenase | 1.35E-36 | 135 | No venom function | NA |
| TRINITY_DN1533_c3_g1_i1.p1 | TRINITY_DN1533_c3_g1_i1 | 22.82425731 | Venom acid phosphatase Acph-1 | 3.4E-70 | 221 | Venom acid phosphatase Acph-1 | 3.6E-71 | 221 | Uncharacterized protein | 0 | 754 | Toxin | Venom acid phosphatase Acph-1 |
| TRINITY_DN176_c0_g2_i29.p1 | TRINITY_DN176_c0_g2_i29 | 9.562062815 | Venom metalloproteinase 3 | 6.8E-34 | 129 | Venom metalloproteinase 3 | 7.22E-35 | 129 | Peptidase M12B domain-containing protein | 0 | 1202 | Toxin | Metalloprotease M12B domain |
| TRINITY_DN18396_c0_g1_i1.p5 | TRINITY_DN18396_c0_g1_i1 | 2.88596734 | Melittin | 4.03E-08 | 39.7 | Melittin | 4.28E-09 | 39.7 | Melittin | 0.0000716 | 39.7 | Toxin | Mellitin |
| TRINITY_DN19039_c1_g1_i1.p1 | TRINITY_DN19039_c1_g1_i1 | 1.463235366 | Melittin | 8.03E-10 | 48.9 | Melittin | 8.52E-11 | 48.9 | Uncharacterized protein | 5.32E-92 | 284 | Toxin | Mellitin |
| TRINITY_DN19063_c0_g1_i4.p1 | TRINITY_DN19063_c0_g1_i4 | 10.28040319 | Venom serine protease inhibitor | 3.63E-12 | 52.8 | Chymotrypsin inhibitor | 4.64E-38 | 115 | Chymotrypsin inhibitor | 1.66E-48 | 152 | No venom function | Chymotrypsin inhibitor |
| TRINITY_DN1908_c0_g1_i1.p1 | TRINITY_DN1908_c0_g1_i1 | 95.3626321 | Vascular endothelial growth factor A | 4.61E-08 | 48.9 | NA | NA | NA | PDGF_2 domain-containing protein | 0 | 552 | No venom function | NA |
| TRINITY_DN19171_c1_g1_i1.p1 | TRINITY_DN19171_c1_g1_i1 | 9.718814787 | Melittin | 4.8E-14 | 56.2 | Melittin | 5.09E-15 | 56.2 | Melittin | 6.42E-11 | 56.6 | Toxin | Mellitin |
| TRINITY_DN2004_c0_g1_i1.p1 | TRINITY_DN2004_c0_g1_i1 | 19.89218392 | NA | NA | NA | NA | NA | NA | Uncharacterized protein/Hypoxia up-regulated protein | 0 | 741 | No venom function | NA |
| TRINITY_DN2082_c1_g1_i4.p9 | TRINITY_DN2082_c1_g1_i4 | 13.60368853 | NA | NA | NA | NA | NA | NA | ML domain-containing protein | 1.24E-23 | 90.9 | No venom function | NA |
| TRINITY_DN21858_c0_g1_i1.p1 | TRINITY_DN21858_c0_g1_i1 | 36.72202147 | Calglandulin | 3E-40 | 129 | NA | NA | NA | Putative calmodulin isoform x3 (Fragment) | 2.92E-105 | 305 | Venom component | Calglandulin |
| TRINITY_DN2306_c0_g1_i1.p1 | TRINITY_DN2306_c0_g1_i1 | 116.4239432 | NA | NA | NA | NA | NA | NA | Glutaredoxin-C4 | 4.48E-69 | 207 | No venom function | NA |
| TRINITY_DN2882_c0_g1_i3.p3 | TRINITY_DN2882_c0_g1_i3 | 34.73149601 | Melittin | 3.14E-13 | 53.9 | Melittin | 3.34E-14 | 53.9 | Melittin | 7E-10 | 53.9 | Toxin | Mellitin |
| TRINITY_DN30275_c0_g1_i1.p1 | TRINITY_DN30275_c0_g1_i1 | 42.85076456 | NA | NA | NA | NA | NA | NA | Sep15_SelM domain-containing protein | 6.83E-112 | 319 | No venom function | NA |
| TRINITY_DN3177_c0_g1_i2.p4 | TRINITY_DN3177_c0_g1_i2 | 13.43895102 | Melittin | 0.00000646 | 34.3 | Melittin | 6.86E-07 | 34.3 | NA | NA | NA | Toxin | Mellitin |
| TRINITY_DN3221_c0_g1_i3.p1 | TRINITY_DN3221_c0_g1_i3 | 202.6841927 | Melittin | 2.3E-13 | 55.8 | Melittin | 2.44E-14 | 55.8 | Melittin | 2.51E-10 | 56.2 | Toxin | Mellitin |
| TRINITY_DN337_c0_g1_i3.p1 | TRINITY_DN337_c0_g1_i3 | 32.08444826 | NA | NA | NA | NA | NA | NA | Uncharacterized protein/CarboxypeptidaseD | 7.41E-86 | 312 | Venom component | CarboxypeptidaseD |
| TRINITY_DN350_c7_g1_i1.p1 | TRINITY_DN350_c7_g1_i1 | 17808.41482 | Melittin | 6.38E-15 | 58.5 | Melittin | 6.77E-16 | 58.5 | Melittin | 5.2E-12 | 59.3 | Toxin | Mellitin |
| TRINITY_DN354_c0_g1_i1.p1 | TRINITY_DN354_c0_g1_i1 | 104.5798778 | NA | NA | NA | NA | NA | NA | Uncharacterized protein (with signalpeptide) | 0 | 831 | Unknown function | NA |
| TRINITY_DN36046_c0_g1_i1.p1 | TRINITY_DN36046_c0_g1_i1 | 28.27763789 | NA | NA | NA | Histone H4 | 1.44E-70 | 199 | Histone H4 (Fragment) | 9.39E-67 | 201 | No venom function | NA |
| TRINITY_DN3709_c1_g1_i1.p1 | TRINITY_DN3709_c1_g1_i1 | 13.93598945 | NA | NA | NA | NA | NA | NA | Uncharacterized protein (with signalpeptide) | 0 | 568 | Unknown function | NA |
| TRINITY_DN3723_c0_g2_i1.p1 | TRINITY_DN3723_c0_g2_i1 | 63.79607937 | NA | NA | NA | Cytochrome c | 2.42E-79 | 222 | Cytochrome c | 3.6E-75 | 226 | No venom function | NA |
| TRINITY_DN38_c0_g2_i3.p1 | TRINITY_DN38_c0_g2_i3 | 918.4574656 | NA | NA | NA | Apidaecins type 22 | 7.24E-57 | 166 | Apidaecins type 22 | 1.52E-52 | 166 | Venom component, AMP? | Apidaecin |
| TRINITY_DN3849_c0_g2_i3.p9 | TRINITY_DN3849_c0_g2_i3 | 19.99364921 | Melittin | 4.18E-09 | 42.4 | Melittin | 4.44E-10 | 42.4 | Melittin | 0.00000774 | 42.4 | Toxin | Mellitin |
| TRINITY_DN38821_c0_g1_i1.p5 | TRINITY_DN38821_c0_g1_i1 | 3.853925957 | NA | NA | NA | NA | NA | NA | Uncharacterized/RecBCD enzyme subunit RecC | 1.70E+00 | 77 | No venom function | NA |
| TRINITY_DN3929_c0_g1_i1.p3 | TRINITY_DN3929_c0_g1_i1 | 23.31658417 | Melittin | 5.9E-08 | 39.3 | Melittin | 6.27E-09 | 39.3 | Melittin | 0.000131 | 39.3 | Toxin | Mellitin |
| TRINITY_DN400_c0_g1_i2.p1 | TRINITY_DN400_c0_g1_i2 | 12.21032338 | NA | NA | NA | NA | NA | NA | Renin receptor | 0 | 777 | No venom function | NA |
| TRINITY_DN45320_c0_g1_i1.p1 | TRINITY_DN45320_c0_g1_i1 | 171.521325 | NA | NA | NA | NA | NA | NA | ATP synthase subunit beta | 0 | 1034 | No venom function | NA |
| TRINITY_DN5_c2_g1_i1.p1 | TRINITY_DN5_c2_g1_i1 | 328360.2366 | Melittin | 4.65E-47 | 140 | Melittin | 4.94E-48 | 140 | Melittin | 1.04E-43 | 140 | Toxin | Mellitin |
| TRINITY_DN5_c2_g1_i1.p2 | TRINITY_DN5_c2_g1_i1 | 328360.2366 | Melittin | 4.65E-47 | 140 | Melittin | 4.94E-48 | 140 | Melittin | 1.04E-43 | 140 | Toxin | Mellitin |
| TRINITY_DN55_c0_g3_i1.p1 | TRINITY_DN55_c0_g3_i1 | 305.6365745 | Venom serine carboxypeptidase | 0 | 962 | Venom serine carboxypeptidase | 0 | 962 | Venom serine carboxypeptidase | 0 | 962 | Toxin | Venom serine carboxypeptidase |
| TRINITY_DN5978_c0_g1_i1.p1 | TRINITY_DN5978_c0_g1_i1 | 39.22797479 | NA | NA | NA | NA | NA | NA | Uncharacterized protein/Peptidylglycine alpha-hydroxy | 3.48E-138 | 399 | No venom function | NA |
| TRINITY_DN68_c0_g1_i1.p1 | TRINITY_DN68_c0_g1_i1 | 473.3351107 | NA | NA | NA | NA | NA | NA | Uncharacterized protein/Actin | 0 | 784 | No venom function | NA |
| TRINITY_DN6995_c0_g2_i2.p4 | TRINITY_DN6995_c0_g2_i2 | 19.97189537 | Melittin | 9.34E-12 | 49.7 | Melittin | 9.91E-13 | 49.7 | Melittin | 9.32E-09 | 50.4 | Toxin | Mellitin |
| TRINITY_DN7033_c0_g1_i3.p1 | TRINITY_DN7033_c0_g1_i3 | 73.17558486 | NA | NA | NA | NA | NA | NA | Aldose 1-epimerase | 0 | 695 | No venom function | NA |
| TRINITY_DN7492_c0_g1_i1.p8 | TRINITY_DN7492_c0_g1_i1 | 6.615274449 | Melittin | 1.89E-14 | 56.2 | Melittin | 2.01E-15 | 56.2 | Melittin | 2.17E-11 | 56.6 | Toxin | Mellitin |
| TRINITY_DN8464_c0_g1_i1.p1 | TRINITY_DN8464_c0_g1_i1 | 18.88211464 | NA | NA | NA | NA | NA | NA | Uncharacterized protein/Endoplasmic reticulum reside | 1.35E-36 | 135 | No venom function | NA |
| TRINITY_DN8464_c0_g1_i2.p1 | TRINITY_DN8464_c0_g1_i2 | 5.036395315 | NA | NA | NA | NA | NA | NA | Uncharacterized protein/Endoplasmic reticulum reside | 3.40E-45 | 155 | No venom function | NA |
| TRINITY_DN8492_c0_g1_i2.p5 | TRINITY_DN8492_c0_g1_i2 | 162.610691 | Melittin | 1.96E-11 | 50.1 | Melittin | 2.08E-12 | 50.1 | Melittin | 4.22E-08 | 50.1 | Toxin | Mellitin |
| TRINITY_DN8560_c0_g1_i1.p2 | TRINITY_DN8560_c0_g1_i1 | 24.08815798 | Venom acid phosphatase Acph-1 | 0 | 800 | Venom acid phosphatase Acph-1 | 0 | 800 | Venom acid phosphatase Acph-1 | 0 | 826 | Toxin | Venom acid phosphatase Acph-1 |
| TRINITY_DN9_c0_g1_i1.p1 | TRINITY_DN9_c0_g1_i1 | 5370.349558 | Secapin | 3.9E-53 | 156 | Secapin | 4.14E-54 | 156 | Secapin | 8.68E-50 | 156 | Toxin | Secapin |
| TRINITY_DN999_c3_g1_i1.p1 | TRINITY_DN999_c3_g1_i1 | 68.79533918 | Hyaluronidase | 0 | 691 | Hyaluronidase | 0 | 691 | Hyaluronidase | 0 | 691 | Toxin | Hyaluronidase |

**Proteome data analysis**

| Protein Grou | Protein ID | Accession | -10lgP | Coverage (% | #Peptides | #Unique | PTM | Avg. Mass | Description |
|---|---|---|---|---|---|---|---|---|---|
| 299 | 90 | J1024640.p1 | 104.79 | 19 | 4 | 4 | | 27219 | GENE.J1024640~~J1024640.p1 ORF type:complete len:238 (+) score=35.56 zf-met2\|PF12907.7\|0.39 zf-met2\|PF12907.7\|7.5e+02 J1024640:66-779(+) |
| 270 | 129 | NODE_10174_length_1471_cov_85.495702_g6621_i0.p1 | 77.69 | 24 | 4 | 4 | Carbamidomethylation; Pyro-glu from Q | 24579 | Trypsin\|PF00089.26\|1.1e-50 Trypsin_2\|PF13365.6\|0.00015 NODE_1... |
| 9 | 72 | NODE_10403_length_1439_cov_3311.402493_g351_i3.p6 | 372.43 | 51 | 98 | 9 | Oxidation (M); Amidation; Acetylation (K); Acetylation (Protein N-term); Biotinylation; 68 more | 6829 | score=1.43 Melittin\|PF01372.17\|6e-17 Melittin\|PF01372.17\|7.2e+03 ... |
| 171 | 85 | NODE_10530_length_1432_cov_7.620915_g7215_i0.p1 | 155.27 | 22 | 6 | 6 | Carbamidomethylation; Ammonia-loss (N) | 44697 | Trypsin\|PF00089.26\|2e-48 Trypsin_2\|PF13365.6\|0.00012 DUF19... |
| 3 | 10 | NODE_11040_length_1349_cov_3010.901099_g4121_i3.p1 | 437.95 | 79 | 171 | 19 | Carbamidomethylation; Oxidation (M); Amidation; Acetylation (K); Acetylation (N-term); 96 more | 19058 | score=19.23 Phospholip_A2_2\|PF05826.12\|1.2e-34 Phospholip_A2_1\|PF00... |
| 884 | 148 | NODE_11279_length_1337_cov_237.078783_g5817_i1.p1 | 86.59 | 11 | 2 | 2 | | 33763 | Pyrid_oxidase_2\|PF13883.6\|1e-47 NODE_11279_length_1337_cov... |
| 42 | 133 | NODE_11920_length_1264_cov_21752.067825_g8218_i0.p1 | 127.8 | 20 | 9 | 9 | Carbamidomethylation; Amidation; Formylation; Hydroxylation; Ubiquitin; 9 more | 5767 | 3.06 Bee_toxin\|PF17454.2\|2.7e-32 NODE_11920_length_1264_co... |
| 45 | 206 | NODE_11920_length_1264_cov_21752.067825_g8218_i0.p3 | 148 | 43 | 13 | 3 | Carbamidomethylation; Oxidation (M); Carboxymethyl; Deamidation (NQ); Dehydration; 10 more | 5223 | 0.58 Bee_toxin\|PF17454.2\|2e-28 NODE_11920_length_1264_cov_... |
| 18 | 146 | NODE_12118_length_1243_cov_10805.042088_g8373_i0.p9 | 279.13 | 100 | 67 | 2 | Oxidation (M); Amidation; Acetylation (K); Acetylation (Protein N-term); Dehydration; 66 more | 3145 | score=-2.34 Melittin\|PF01372.17\|2.9e-12 NODE_12118_length_1... |
| 1534 | 167 | NODE_13401_length_1105_cov_468.260000_g9314_i0.p1 | 63.7 | 13 | 2 | 2 | Carbamidomethylation (DHKE X@N-term) | 18290 | AhpC-TSA\|PF00578.21\|7e-23 1-cysPrx_C\|PF10417.9\|3.5e-15 Red... |
| 101 | 242 | NODE_143_length_9096_cov_32.937286_g93_i0.p1 | 60.95 | 2 | 4 | 4 | Deoxy; Mutation | 119336 | HCO3_cotransp\|PF00955.21\|8.9e-236 Band_3_cyto\|PF07565.13\|1.6e-100 NODE_... |
| 73 | 31 | NODE_14318_length_1026_cov_588.271885_g10016_i0.p1 | 168.41 | 70 | 11 | 11 | Carbamidomethylation; Deamidation (NQ); Formylation (Protein N-term); Methyl ester; Phosphorylation (STY); Carbamidomethylation (DHKE X@N-term) | 15634 | GENE.NODE_14318_length_1026_cov_588.271885_g10016_i0~~NODE_14318_length_1026_cov_588.271885_g10016_i0.p1 ORF type:complete len:153 (+) score=45.85 Sod_Cu\|PF00080.20\|1.1e-51 NODE_14318_length_1026_cov_58... |
| 58 | 19 | NODE_14549_length_987_cov_148.449561_g9868_i0.p1 | 192.35 | 57 | 15 | 15 | Carbamidomethylation; Oxidation (M); Deamidation (NQ); Dehydration; Oxidation (HW); 3 more | 22644 | Pro_isomerase\|PF00160.21\|2.2e-51 NODE_14549_length_987_cov_14... |
| 67 | 79 | NODE_14992_length_970_cov_72249.791257_g6125_i4.p2 | 191.32 | 65 | 14 | 14 | Carbamidomethylation; Acetylation (Protein N-term); Hydroxylation; Pyro-glu from E; Carbamidomethylation (DHKE X@N-term); 2 more | 11234 | GENE.NODE_14992_length_970_cov_72249.791257_g6125_i4~~NODE_14992_length_970_cov_72249.791257_g6125_i4.p2 ORF type:complete len:100 (+) score=12.68 DX\|PF01666.17\|0.0049 Nodulin_late\|PF07127.11\|2.8e+02 No... |
| 209 | 88 | NODE_15131_length_938_cov_131.254925_g10349_i0.p2 | 87.29 | 50 | 5 | 5 | Oxidation (M) | 11381 | CENP-T_C\|PF15511.6\|2.2e-08 TAF\|PF02969.17\|3.6e-05 CENP-S\|P... |
| 1534 | 168 | NODE_15441_length_913_cov_534.093079_g10606_i0.p1 | 63.7 | 11 | 2 | 2 | Carbamidomethylation (DHKE X@N-term) | 21815 | AhpC-TSA\|PF00578.21\|1.7e-38 1-cysPrx_C\|PF10417.9\|4.7e-15 R... |
| 109 | 62 | NODE_1577_length_4346_cov_815.361920_g1019_i0.p1 | 115.36 | 31 | 9 | 9 | Amidation; Dehydration; Pyro-glu from Q; Carbamidomethylation (DHKE X@N-term) | 41806 | Actin\|PF00022.19\|9.3e-151 NODE_1577_length_4346_cov_815.36192... |
| 41 | 311 | NODE_16160_length_860_cov_4556.777070_g11199_i0.p1 | 57.99 | 10 | 2 | 2 | Methylation; Sodium adduct; Carbamidomethylation (DHKE X@N-term); Carboxymethyl (KW X@N-term); Dihydroxy; Mutation | 8664 | GENE.NODE_16160_length_860_cov_4556.777070_g11199_i0~~NODE_16160_length_860_cov_4556.777070_g11199_i0.p1 ORF type:complete len:78 (+) score=6.95 Secapin\|PF17521.2\|1e-34 NODE_16160_length_860_cov_4556.77... |
| 271 | 199 | NODE_18028_length_766_cov_1892.343179_g8026_i1.p1 | 127.33 | 42 | 3 | 3 | Carbamidomethylation; Oxidation (M); Methyl ester | 8298 | Toxin_30\|PF08117.11\|0.0063 Cys_Knot_tox\|PF17486.2\|0.0061 To... |
| 42 | 132 | NODE_18339_length_747_cov_15153.239884_g8218_i2.p1 | 127.8 | 20 | 9 | 9 | Carbamidomethylation; Amidation; Formylation; Hydroxylation; Ubiquitin; 9 more | 5767 | Bee_toxin\|PF17454.2\|2.7e-32 NODE_18339_length_747_cov_15... |
| 209 | 87 | NODE_19733_length_678_cov_176.370787_g14498_i0.p1 | 87.29 | 50 | 5 | 5 | Oxidation (M) | 11381 | CENP-T_C\|PF15511.6\|2.2e-08 TAF\|PF02969.17\|3.6e-05 CENP-S\|P... |
| 1137 | 124 | NODE_20261_length_635_cov_63.437500_g14817_i0.p1 | 111.87 | 33 | 3 | 3 | Carbamidomethylation | 18106 | score=11.30 PBP_GOBP\|PF01395.22\|1.2e-15 NODE_20261_length_635_cov_6... |
| 100 | 119 | NODE_20389_length_648_cov_68298.664418_g15085_i0.p1 | 89.67 | 34 | 4 | 4 | Carbamidomethylation; Amidation | 15204 | score=26.18 PBP_GOBP\|PF01395.22\|2.9e-12 NODE_20389_length_648_co... |
| 1139 | 223 | NODE_2447_length_3705_cov_272.814050_g1430_i0.p1 | 89.26 | 20 | 2 | 2 | Acetylation (Protein N-term) | 16811 | hand_1\|PF00036.32\|1.2e-09 EF-hand_1\|PF00036.32\|8.9e-09 EF-... |
| 70 | 104 | NODE_24540_length_505_cov_3301.937778_g18995_i0.p1 | 162.25 | 41 | 6 | 6 | Piperidination | 16911 | score=26.62 C1q\|PF00386.21\|3.3e-07 NODE_24540_length_505_cov_... |
| 24 | 145 | NODE_2528_length_3602_cov_40.640541_g1631_i0.p30 | 275.38 | 95 | 42 | 16 | Amidation; Carbamylation; Dehydration; Formylation (Protein N-term); Carboxylation (E); 15 more | 1899 | NODE_2528_length_3602_cov_40.640541_g1631_i0:1-63(+) |
| 136 | 139 | NODE_27106_length_438_cov_721.168044_g21398_i0.p1 | 127.91 | 53 | 4 | 2 | more | 8752 | score=8.60 NODE_27106_length_438_cov_721.168044_g21398_i0:2-232(+... |
| 248 | 330 | NODE_28629_length_412_cov_2765.735905_g22893_i0.p3 | 48.9 | 13 | 2 | 2 | | 14690 | score=45.94 Apidaecin\|PF00807.17\|9.9e-06 Apidaecin\|PF00807.17... |
| 109 | 61 | NODE_3011_length_3324_cov_1067.924748_g1019_i1.p1 | 115.36 | 31 | 9 | 9 | Amidation; Dehydration; Pyro-glu from Q; Carbamidomethylation (DHKE X@N-term) | 41806 | Actin\|PF00022.19\|9.3e-151 NODE_3011_length_3324_cov_1067.9... |
| 49 | 5 | NODE_3142_length_3280_cov_453.149766_g1714_i1.p1 | 301.81 | 40 | 37 | 37 | Carbamidomethylation; Oxidation (M); Acetylation (N-term); Carbamylation; Deamidation (NQ); 6 more | 100085 | SapB_2\|PF03489.17\|2.2e+03 SapB_2\|PF03489.17\|2.2e-10 SapB_2\|P... |
| 50 | 6 | NODE_3646_length_3065_cov_183.230769_g1489_i3.p1 | 297.03 | 66 | 29 | 19 | Carbamidomethylation; Oxidation (M); Deamidation (NQ); Hexose (NSY); Methyl ester; 7 more | 48688 | MRJP\|PF03022.16\|1.5e-92 NODE_3646_length_3065_cov_183.230769_... |
| 38 | 210 | NODE_39837_length_291_cov_0.983051_g34173_i0.p1 | 137.59 | 39 | 13 | 6 | Acetylation (N-term); Carbamylation; Methyl ester; Oxidation (HW); Palmitoylation; 8 more | 8034 | Melittin\|PF01372.17\|2.8e-15 Trypan_PARP\|PF05887.11\|0.0018 NO... |
| 110 | 181 | NODE_40674_length_288_cov_0.824034_g35010_i0.p2 | 96.93 | 40 | 6 | 2 | Carbamidomethylation; Acetylation (N-term); Carbamylation; Deamidation (NQ); Dehydration; 4 more | 6431 | Phospholip_A2_2\|PF05826.12\|0.0015 NODE_40674_length_288_cov_... |
| 37 | 4 | NODE_4090_length_2882_cov_402.678660_g2412_i0.p1 | 329.14 | 47 | 42 | 27 | Carbamidomethylation; Oxidation (M); Acetylation (N-term); Deamidation (NQ); Hydroxylation; 9 more | 99461 | score=207.25 DPPIV_N\|PF00930.21\|2.2e-78 Peptidase_S9\|PF00326.21\|5.6... |
| 407 | 142 | NODE_4142_length_2818_cov_303.087948_g453_i1.p1 | 66.95 | 13 | 2 | 2 | | 25565 | HSP20\|PF00011.21\|2.1e-18 ArsA_HSP20\|PF17886.1\|0.0026 AATF-Che1\|P... |
| 65 | 45 | NODE_4236_length_2786_cov_241.753936_g2779_i0.p1 | 179.29 | 58 | 12 | 11 | Carbamidomethylation; Oxidation (M); Methyl ester; Methylation; Sodium adduct; 4 more | 11411 | DUF4789\|PF16033.5\|7.3e-05 NODE_4236_length_2786_cov_241.753936... |
| 76 | 113 | NODE_44489_length_278_cov_0.748768_g38728_i0.p1 | 123.64 | 35 | 6 | 2 | Dihydroxy | 10587 | NODE_44489_length_278_cov_0.748768_g38728_i0:2-277(+) |
| 4 | 48 | NODE_49307_length_261_cov_541237.096774_g43546_i0.p1 | 372.56 | 87 | 126 | 25 | Oxidation (M); Amidation; Acetylation (K); Acetylation (Protein N-term); Acetylation (N-term); 74 more | 8697 | score=19.63 Melittin\|PF01372.17\|7.8e-16 Melittin\|PF01372.17\|7.... |
| 40 | 178 | NODE_50813_length_248_cov_1.051813_g45149_i0.p1 | 152.06 | 23 | 14 | 7 | Acetylation (K); Acetylation (N-term); Carbamylation; Myristoylation; Oxidation (HW); 9 more | 7221 | Melittin\|PF01372.17\|5.5e-11 Trypan_PARP\|PF05887.11\|0.00098 S... |
| 79 | 56 | NODE_5128_length_2490_cov_1011.854620_g3398_i0.p1 | 141.79 | 49 | 7 | 7 | Carbamidomethylation; Methyl ester; Carbamidomethylation (DHKE X@N-term) | 25317 | score=61.51 Glyco_hydro_18\|PF00704.28\|8.4e-29 NODE_5128_length_2... |
| 32 | 73 | NODE_51757_length_254_cov_0.849162_g45996_i0.p2 | 222.77 | 71 | 43 | 30 | Amidation; Acetylation (N-term); Dehydration; Hexose (NSY); Oxidation (HW); 17 more | 6196 | Melittin\|PF01372.17\|2.6e-16 Trypan_PARP\|PF05887.11\|0.0011 D... |
| 44 | 76 | NODE_53677_length_249_cov_0.873563_g47916_i0.p1 | 169.24 | 77 | 15 | 4 | Carbamidomethylation; Acetylation (K); Deamidation (NQ); Dehydration; Carboxylation (E); 13 more | 5237 | Phospholip_A2_2\|PF05826.12\|1.4e-11 NODE_53677_length_249_cov... |
| 19 | 153 | NODE_53886_length_248_cov_1.017341_g48125_i0.p1 | 253.69 | 48 | 66 | 16 | Oxidation (M); Amidation; Acetylation (K); Acetylation (N-term); Dehydration; 56 more | 7060 | Melittin\|PF01372.17\|4.5e-14 Trypan_PARP\|PF05887.11\|0.0019 SP... |
| 16 | 154 | NODE_54636_length_223_cov_8.095238_g48972_i0.p1 | 287.42 | 32 | 75 | 11 | Oxidation (M); Amidation; Acetylation (K); Acetylation (N-term); Biotinylation; 65 more | 8035 | Melittin\|PF01372.17\|1.4e+03 Melittin\|PF01372.17\|6.3e-14 Pilosulin\|... |
| 36 | 24 | NODE_6700_length_2072_cov_10229.525037_g4464_i1.p1 | 261.69 | 65 | 31 | 26 | Oxidation (M); Deamidation (NQ); Hydroxylation; Methyl ester; Oxidation (HW); 16 more | 24490 | score=41.71 NODE_6700_length_2072_cov_10229.525037_g4464_i1:1025-16... |
| 2096 | 205 | NODE_6769_length_2084_cov_39.210553_g4191_i0.p7 | 58.37 | 64 | 2 | 2 | Carbamidomethylation | 4649 | E1_DerP2_DerF2\|PF02221.15\|0.022 NODE_6769_length_2084_cov_39.21055... |
| 65 | 44 | NODE_6915_length_2046_cov_279.860984_g4286_i0.p1 | 179.29 | 58 | 12 | 11 | Carbamidomethylation; Oxidation (M); Methyl ester; Methylation; Sodium adduct; 4 more | 11411 | DUF4789\|PF16033.5\|7.3e-05 NODE_6915_length_2046_cov_279.860984... |
| 357 | 96 | NODE_7578_length_1914_cov_211.404024_g4758_i0.p1 | 112.03 | 11 | 4 | 4 | 3-sulfanylpropanoyl | 47186 | Enolase_C\|PF00113.22\|4.7e-153 Enolase_N\|PF03952.16\|6.4e-57 E... |
| 2096 | 204 | NODE_7698_length_1875_cov_36.041209_g4442_i1.p3 | 58.37 | 64 | 2 | 2 | Carbamidomethylation | 4649 | E1_DerP2_DerF2\|PF02221.15\|0.022 NODE_7698_length_1875_cov_36.04120... |
| 712 | 184 | NODE_7890_length_1840_cov_98.634734_g5077_i1.p1 | 54.21 | 5 | 2 | 2 | Carbamidomethylation | 56432 | Peptidase_S28\|PF05577.12\|2.4e-103 Abhydrolase_1\|PF00561.20\|2.8e+... |
| 21 | 77 | NODE_7908_length_1851_cov_1172.670045_g4121_i1.p4 | 401.76 | 86 | 105 | 4 | more | 7740 | Phospholip_A2_2\|PF05826.12\|7.1e-12 NODE_7908_length_1851_cov... |
| 64 | 16 | NODE_7958_length_1825_cov_1940.759322_g5357_i0.p1 | 182.92 | 31 | 16 | 14 | Hex3 | 66882 | Transferrin\|PF00405.17\|4.3e-22 Transferrin\|PF00405.17\|4.5... |
| 711 | 98 | NODE_827_length_5603_cov_227.112112_g520_i1.p1 | 104.18 | 19 | 6 | 6 | Carbamidomethylation; Amidation; Deamidation (NQ) | 50580 | Hexokinase_2\|PF03727.16\|2.6e-83 Hexokinase_1\|PF00349.21\|6.9e-70 Hex... |
| 29 | 38 | NODE_8360_length_1756_cov_674.852440_g5640_i0.p1 | 222.73 | 61 | 44 | 44 | Carbamidomethylation; Acetylation (K); Acetylation (N-term); Carbamylation; Carboxymethyl; 16 more | 10283 | TIL\|PF01826.17\|7.5e+03 TIL\|PF01826.17\|2.7e-12 NODE_8360_length_... |
| 99 | 55 | NODE_8622_length_1713_cov_138.656212_g588_i1.p1 | 152.84 | 19 | 6 | 2 | Carbamidomethylation; Phosphorylation (STY) | 51690 | Peptidase_M14\|PF00246.24\|3.6e-80 CarboxypepD_reg\|PF13620.6\|1.4e... |
| 358 | 118 | NODE_9018_length_1656_cov_105.966477_g5751_i0.p1 | 76.25 | 14 | 5 | 5 | Carbamidomethylation (DHKE X@N-term) | 35802 | Ldh_1_C\|PF02866.18\|2e-51 Ldh_1_N\|PF00056.23\|3.9e-47 3Beta_HSD... |
| 99 | 54 | NODE_9154_length_1625_cov_145.924204_g588_i2.p1 | 152.84 | 21 | 6 | 2 | Carbamidomethylation; Phosphorylation (STY) | 48573 | Peptidase_M14\|PF00246.24\|3e-80 CarboxypepD_reg\|PF13620.6\|1.3e-14... |
| 35 | 1 | NODE_9157_length_1632_cov_16.547848_g5858_i0.p1 | 347.08 | 66 | 43 | 43 | Carbamidomethylation; Oxidation (M); Acetylation (N-term); Biotinylation; Carbamylation; 21 more | 46872 | His_Phos_2\|PF00328.22\|2.7e-12 His_Phos_2\|PF00328.22\|4e-06 NODE_9... |
| 33 | 8 | R1024725.p1 | 305.58 | 69 | 46 | 43 | Carbamidomethylation; Oxidation (M); Amidation; Acetylation (K); Acetylation (N-term); 21 more | 38425 | 1721(+) |
| 486 | 143 | R1024865.p1 | 63.26 | 4 | 2 | 2 | | 65843 | 3366(+) |
| 53 | 11 | R1025397.p1 | 285.25 | 54 | 27 | 25 | Carbamidomethylation; Deamidation (NQ); Methyl ester; Oxidation (HW); Sodium adduct; 6 more | 46958 | GENE.R1025397~~R1025397.p1 ORF type:complete len:417 (+) score=40.94 MRJP\|PF03022.16\|2.2e-92 R1025397:57-1307(+) |
| 487 | 150 | R1025449.p1 | 74.24 | 8 | 3 | 3 | Acetylation (N-term); Carbamidomethylation (DHKE X@N-term) | 42255 | GENE.R1025449~~R1025449.p1 ORF type:complete len:379 (+) score=44.39 Aldose_epim\|PF01263.20\|1.1e-86 R1025449:579-1715(+) |
| 57 | 13 | R1025530.p1 | 214.66 | 25 | 18 | 18 | Carbamidomethylation; Deamidation (NQ); Methyl ester; Phosphorylation (STY); Sulfation; 2 more | 100380 | Cystatin\|PF00031.21\|0.68 Cystatin\|PF00031.21\|5.9e-12 Cystatin\|PF00031.21\|6.6e+03 Inhibitor_I29\|PF08246.12\|1.4e+04... |
| 161 | 115 | R1025617.p1 | 160.3 | 24 | 5 | 5 | Carbamidomethylation; Deamidation (NQ); Pyro-glu from Q; Carbamidomethylation (DHKE X@N-term) | 39483 | DUF316\|PF03761.15\|0.022 Trypsin_2\|PF13365.6\|0.11 DUF1986\|PF09342.11\|0.12 R1025617:179-1240(+) |
| 95 | 49 | R1027063.p1 | 121.95 | 16 | 7 | 7 | Carbamidomethylation; Carbamidomethylation (DHKE X@N-term) | 60299 | FDF\|PF09532.10\|0.22 R1027063:217-1809(+) |
| 270 | 130 | R1027523.p1 | 77.69 | 20 | 4 | 4 | Carbamidomethylation; Pyro-glu from Q | 29505 | GENE.R1027523~~R1027523.p1 ORF type:complete len:261 (+) score=25.40 Trypsin\|PF00089.26\|2e-53 Trypsin_2\|PF13365.6\|0.00025 R1027523:345-1127(+) |
| 359 | 126 | R1028514.p1 | 63.85 | 23 | 5 | 5 | Carbamidomethylation | 36159 | 33 R1028514:110-1111(+) |
| 125 | 117 | R1029348.p1 | 117.54 | 13 | 7 | 7 | Deamidation (NQ); Mutation | 41283 | R1029348:18-1118(+) |
| 56 | 34 | R1036651.p1 | 193.83 | 36 | 11 | 11 | Carbamidomethylation; Deamidation (NQ); Hydroxylation; Methylation; Oxidation (HW); 3 more | 31288 | Pep1_7\|PF17232.2\|0.26 Phospholip_A2_1\|PF00068.19\|12 Phospholip_A2_1\|PF00068.19\|5.7e+03 R1036651:255-1043(-) |
| 221 | 144 | S1036426.p2 | 120.19 | 32 | 2 | 2 | Carbamidomethylation; Carbamidomethylation (DHKE X@N-term) | 10142 | GENE.S1036426~~S1036426.p2 ORF type:3prime_partial len:93 (+) score=9.43 Lys\|PF00062.20\|2.7e-12 SLT\|PF01464.20\|0.088 S1036426:421-696(+) |
| 1142 | 225 | S1045097.p1 | 62.02 | 6 | 2 | 2 | | 37772 | GENE.S1045097~~S1045097.p1 ORF type:complete len:325 (+) score=104.17 EF-hand_1\|PF00036.32\|0.00036 EF-hand_1\|PF00036.32\|0.16 EF-hand_1\|PF00036.32\|0.00044 EF-hand_1\|PF00036.32\|0.0097 EF-hand_1\|PF00036.32\|7.9 EF-hand_6\|PF13405.6\|0.0011 EF-hand... |
| 135 | 57 | S1047992.p1 | 120.27 | 7 | 5 | 5 | | 107976 | GENE.S1047992~~S1047992.p1 ORF type:complete len:934 (+) score=176.03 Glyco_hydro_31\|PF01055.26\|1.7e-143 Gal_mutarotas_2\|PF13802.6\|3.4e-24 Gal_mutarotas_2\|PF13802.6\|1.9e+03 DUF5110\|PF17137.4\|2.3e-05 DUF5110\|PF17137.4\|1.1e+04 DUF4968\|PF16338.... |
| 17 | 121 | TRINITY_DN10322_c0_g1_i2.p6 | 280.5 | 92 | 67 | 3 | Oxidation (M); Amidation; Acetylation (K); Acetylation (Protein N-term); Dehydration; 66 more | 2939 | TRINITY_DN10322_c0_g1_i2:1-81(+) |
| 1141 | 183 | TRINITY_DN1093_c0_g3_i2.p1 | 55.95 | 14 | 2 | 2 | | 18920 | 636(+) |
| 62 | 32 | TRINITY_DN11539_c0_g1_i7.p1 | 190.19 | 29 | 11 | 11 | Carbamidomethylation; Oxidation (M); Deamidation (NQ); Dehydration; Hydroxylation; 5 more | 43341 | Abhydrolase_3\|PF07859.13\|1e-07 Peptidase_S9\|PF00326.21\|0.095 TRINITY_DN11539_c0_g1_i7:390-1538(+) |
| 150 | 114 | TRINITY_DN11539_c0_g1_i9.p2 | 128.55 | 41 | 5 | 5 | Carbamidomethylation; Carbamidomethylation (DHKE X@N-term) | 17161 | TRINITY_DN11539_c0_g1_i9:1905-2351(+) |
| 47 | 211 | TRINITY_DN1249_c0_g1_i1.p11 | 263.11 | 94 | 19 | 18 | Biotinylation; Dehydration; Formylation (Protein N-term); Hydroxylation; Pyro-glu from E; 8 more | 3118 | TRINITY_DN1249_c0_g1_i1:3-98(+) |
| 34 | 226 | TRINITY_DN12646_c3_g1_i1.p1 | 182.56 | 59 | 27 | 13 | Acetylation (N-term); Carbamylation; Dehydration; Carboxylation (E); Hexose (NSY); 14 more | 6520 | Trypan_PARP\|PF05887.11\|0.00046 TRINITY_DN12646_c3_g1_i1:61-243(+) |
| 149 | 75 | TRINITY_DN1306_c0_g1_i1.p1 | 111.67 | 11 | 5 | 5 | Carbamidomethylation; Methyl ester | 72866 | MreB_Mbl\|PF06723.13\|8.5e-14 FtsA\|PF14450.6\|3.5 FtsA\|PF14450.6\|0.78 FtsA\|PF14450.6\|4.1e... |
| 43 | 291 | TRINITY_DN14348_c0_g2_i1.p1 | 130.82 | 11 | 10 | 4 | Acetylation (N-term); Biotinylation; Carbamylation; Formylation; Pyro-glu from E; 5 more | 17021 | TRINITY_DN14348_c0_g2_i1:2-445(+) |
| 142 | 58 | TRINITY_DN1533_c3_g1_i1.p1 | 119.84 | 20 | 7 | 6 | Carbamidomethylation; Carbamidomethylation (DHKE X@N-term); Dihydroxy; Mutation | 42664 | 11 His_Phos_2\|PF00328.22\|7.6e-05 TRINITY_DN1533_c3_g1_i1:98-1183(+) |
| 220 | 152 | TRINITY_DN176_c0_g2_i29.p1 | 79.46 | 7 | 3 | 3 | Carbamidomethylation (DHKE X@N-term) | 66525 | Reprolysin_2\|PF13574.6\|2.9e-10 Reprolysin\|PF01421.19\|49 Reprolysin\|PF01421.19\|2... |
| 20 | 261 | TRINITY_DN18396_c0_g1_i1.p5 | 234.62 | 95 | 52 | 2 | Oxidation (M); Amidation; Acetylation (K); Acetylation (Protein N-term); Dehydration; 55 more | 2381 | 162(+) |
| 12 | 83 | TRINITY_DN19039_c1_g1_i1.p1 | 350.5 | 23 | 86 | 9 | Oxidation (M); Amidation; Acetylation (K); Acetylation (N-term); Biotinylation; 66 more | 18155 | SSF\|PF00474.17\|1e-07 DUF3792\|PF12670.7\|0.017 CDP-OH_P_transf\|PF01066.21\|1.7 C... |
| 201 | 137 | TRINITY_DN19063_c0_g1_i4.p1 | 137.4 | 46 | 3 | 3 | Carbamidomethylation; Carbamidomethylation (DHKE X@N-term); Mutation | 8130 | 417(+) |
| 52 | 15 | TRINITY_DN1908_c0_g1_i1.p1 | 284.05 | 47 | 23 | 21 | Carbamidomethylation; Oxidation (M); Acetylation (N-term); Deamidation (NQ); Hydroxylation; 12 more | 44732 | PDGF\|PF00341.17\|1.8e+04 TRINITY_DN1908_c0_g1_i1:1-1179(+) |
| 23 | 94 | TRINITY_DN19171_c1_g1_i1.p1 | 247.39 | 90 | 47 | 24 | Amidation; Acetylation (K); Biotinylation; Carbamylation; Deamidation (NQ); 36 more | 5449 | Trypan_PARP\|PF05887.11\|0.023 Pilosulin\|PF17499.2\|0.13 TRINITY_DN19171_c1_g1_i1:3-158(+) |
| 219 | 112 | TRINITY_DN2004_c0_g1_i1.p1 | 91.88 | 5 | 4 | 4 | Carbamidomethylation (DHKE X@N-term) | 107634 | HSP70\|PF00012.20\|8e+03 MreB_Mbl\|PF06723.13\|2.8e+02 MreB_Mbl\|PF06723.13\|0.00018 MreB_Mb... |
| 2096 | 203 | TRINITY_DN2082_c1_g1_i4.p9 | 58.37 | 67 | 2 | 2 | Carbamidomethylation | 4464 | TRINITY_DN2082_c1_g1_i4:654-782(-) |
| 1139 | 222 | TRINITY_DN21858_c0_g1_i1.p1 | 89.26 | 20 | 2 | 2 | Acetylation (Protein N-term) | 16811 | 09 EF-hand_1\|PF00036.32\|7.7e-10 EF-hand_1\|PF00036.32\|6e-10 EF-hand_7\|PF13499.... |
| 2095 | 166 | TRINITY_DN2306_c0_g1_i1.p1 | 52.53 | 28 | 2 | 2 | | 10942 | Thioredoxin_3\|PF13192.6\|0.00028 DSBA\|PF01323.20\|0.0024 DSBA\|PF01323.20\|3e+02 Thioredoxin_2\|PF13098.6\|0.0... |
| 8 | 53 | TRINITY_DN2882_c0_g1_i3.p3 | 367.32 | 69 | 103 | 2 | more | 5193 | 148(+) |
| 356 | 95 | TRINITY_DN30275_c0_g1_i1.p1 | 99.17 | 32 | 4 | 4 | Carbamidomethylation | 17832 | TRINITY_DN30275_c0_g1_i1:105-572(+) |
| 39 | 123 | TRINITY_DN3177_c0_g1_i2.p4 | 125.64 | 100 | 13 | 2 | 3 more | 2170 | 71(+) |
| 5 | 47 | TRINITY_DN3221_c0_g1_i3.p1 | 381.58 | 40 | 115 | 3 | Oxidation (M); Amidation; Acetylation (K); Acetylation (N-term); Biotinylation; 71 more | 9388 | TRINITY_DN3221_c0_g1_i3:40-285(+) |
| 51 | 74 | TRINITY_DN337_c0_g1_i3.p1 | 161.28 | 6 | 6 | 2 | Carbamidomethylation; Phosphorylation (STY); Sulfation | 160356 | Peptidase_M14\|PF00246.24\|2.2e-82 Peptidase_M14\|PF00246.24\|0.0021 Peptidase_M14\|PF00246.24\|28 Peptidase_... |
| 2 | 46 | TRINITY_DN350_c7_g1_i1.p1 | 454.63 | 94 | 150 | 4 | Oxidation (M); Amidation; Acetylation (K); Acetylation (Protein N-term); Acetylation (N-term); 75 more | 5479 | Pilosulin\|PF17499.2\|0.16 TRINITY_DN350_c7_g1_i1:1-159(+) |
| 59 | 122 | TRINITY_DN354_c0_g1_i1.p1 | 165.6 | 17 | 9 | 9 | Methyl ester; Sodium adduct | 52044 | TRINITY_DN354_c0_g1~~TRINITY_DN354_c0_g1_i1.p1 ORF type:complete len:492 (+) score=324.47 TRINITY_DN354_c0_g1_i1:563-2038(+) |
| 209 | 89 | TRINITY_DN36046_c0_g1_i1.p1 | 87.29 | 50 | 5 | 5 | Oxidation (M) | 11381 | S\|PF15630.6\|6.3e-05 Histone\|PF00125.24\|0.0002 CBFD_NFYB_HMF\|PF00808.23\|0.0004... |
| 172 | 86 | TRINITY_DN3709_c1_g1_i1.p1 | 122.54 | 27 | 6 | 6 | Deamidation (NQ); Carbamidomethylation (DHKE X@N-term) | 37314 | TRINITY_DN3709_c1_g1~~TRINITY_DN3709_c1_g1_i1.p1 ORF type:5prime_partial len:329 (+) score=104.76 TRINITY_DN3709_c1_g1_i1:2-988(+) |
| 1535 | 177 | TRINITY_DN3723_c0_g2_i1.p1 | 59.6 | 21 | 2 | 2 | | 11910 | Cytochrome_CBB3\|PF13442.6\|2.7e-07 Cytochrom_C550\|PF14495.6\|0.0072 TRINITY_DN3723_c0_g2_i1:93-419(+) |
| 248 | 329 | TRINITY_DN38_c0_g2_i3.p1 | 48.9 | 19 | 2 | 2 | | 10027 | Apidaecin\|PF00807.17\|1.9e-17 TRINITY_DN38_c0_g2_i3:71-337(+) |
| 13 | 185 | TRINITY_DN3849_c0_g2_i3.p9 | 316.85 | 95 | 85 | 21 | Oxidation (M); Amidation; Acetylation (K); Acetylation (Protein N-term); Biotinylation; 68 more | 2354 | 70(+) |
| 85 | 267 | TRINITY_DN38821_c0_g1_i1.p5 | 64.05 | 29 | 2 | 2 | Amidation | 3380 | TRINITY_DN38821_c0_g1~~TRINITY_DN38821_c0_g1_i1.p5 ORF type:complete len:32 (-) score=9.00 TRINITY_DN38821_c0_g1_i1:91-186(-) |
| 27 | 288 | TRINITY_DN3929_c0_g1_i1.p3 | 149.07 | 76 | 13 | 10 | Carbamidomethylation; Amidation; Acetylation (K); Biotinylation; Carbamylation; 12 more | 2398 | TRINITY_DN3929_c0_g1~~TRINITY_DN3929_c0_g1_i1.p3 ORF type:complete len:22 (+) score=-0.27 Melittin\|PF01372.17\|6.4e-12 TRINITY_DN3929_c0_g1_i1:2-67(+) |
| 116 | 63 | TRINITY_DN400_c0_g1_i2.p1 | 152.13 | 24 | 7 | 7 | Acetylation (N-term); Deamidation (NQ); Methyl ester; Phosphorylation (STY); Carbamidomethylation (DHKE X@N-term) | 44208 | TRINITY_DN400_c0_g1~~TRINITY_DN400_c0_g1_i2.p1 ORF type:complete len:396 (+) score=91.81 Renin_r\|PF07850.14\|5.9e-30 Presenilin\|PF01080.17\|0.31 TRINITY_DN400_c0_g1_i2:85-1272(+) |
| 173 | 165 | TRINITY_DN45320_c0_g1_i1.p1 | 63.17 | 8 | 3 | 3 | Carbamidomethylation (DHKE X@N-term) | 55130 | synt_ab_N\|PF02874.23\|2.3e-21 ATPase_2\|PF01637.18\|0.0092 NB-ARC\|PF00931.22\|0.012 NB-ARC\|PF00931.2... |
| 1 | 43 | TRINITY_DN5_c2_g1_i1.p1 | 467.82 | 76 | 159 | 4 | Oxidation (M); Amidation; Acetylation (K); Acetylation (N-term); Biotinylation; 73 more | 7585 | OAD_gamma\|PF04277.13\|0.049 OAD_gamma\|PF04277.13\|2.8e+03 SPESP1\|PF15754.5\|0.034 Pilosuli... |
| 1 | 42 | TRINITY_DN5_c2_g1_i1.p2 | 467.82 | 76 | 159 | 4 | Oxidation (M); Amidation; Acetylation (K); Acetylation (N-term); Biotinylation; 73 more | 7585 | OAD_gamma\|PF04277.13\|0.049 OAD_gamma\|PF04277.13\|2.8e+03 SPESP1\|PF15754.5\|0.034 Pilosuli... |
| 84 | 84 | TRINITY_DN55_c0_g3_i1.p1 | 158.44 | 13 | 4 | 4 | Carbamidomethylation; Carbamidomethylation (DHKE X@N-term) | 53720 | 2902(+) |
| 300 | 92 | TRINITY_DN5978_c0_g1_i1.p1 | 103.12 | 15 | 4 | 4 | Carbamidomethylation | 38714 | Cu2_monooxygen\|PF01082.20\|4.4e-30 Cu2_monooxygen\|PF01082.20\|2.8e+02 TRINITY_DN5978_c0_g1_i1:1048-2073(+... |
| 109 | 60 | TRINITY_DN68_c0_g1_i1.p1 | 115.36 | 31 | 9 | 9 | Amidation; Dehydration; Pyro-glu from Q; Carbamidomethylation (DHKE X@N-term) | 41806 | TRINITY_DN68_c0_g1~~TRINITY_DN68_c0_g1_i1.p1 ORF type:complete len:377 (+) score=48.93 Actin\|PF00022.19\|9.3e-151 TRINITY_DN68_c0_g1_i1:171-1301(+) |
| 6 | 52 | TRINITY_DN6995_c0_g2_i2.p4 | 368.02 | 100 | 104 | 5 | Oxidation (M); Amidation; Acetylation (K); Acetylation (Protein N-term); Biotinylation; 70 more | 3935 | 108(+) |
| 487 | 151 | TRINITY_DN7033_c0_g1_i3.p1 | 74.24 | 8 | 3 | 3 | Acetylation (N-term); Carbamidomethylation (DHKE X@N-term) | 45046 | 1479(+) |
| 11 | 68 | TRINITY_DN7492_c0_g1_i1.p8 | 375.71 | 100 | 96 | 3 | Oxidation (M); Amidation; Acetylation (K); Acetylation (Protein N-term); Biotinylation; 67 more | 3018 | 89(+) |
| 68 | 17 | TRINITY_DN8464_c0_g1_i1.p1 | 148.55 | 49 | 11 | 11 | Carbamidomethylation; Carbamidomethylation (DHKE X@N-term) | 27543 | ERp29\|PF07749.12\|6e+03 ERp29\|PF07749.12\|4.8e-25 RNA_pol_Rpb5_C\|PF01191.19\|20 RNA_po... |
| 68 | 18 | TRINITY_DN8464_c0_g1_i2.p1 | 148.55 | 49 | 11 | 11 | Carbamidomethylation; Carbamidomethylation (DHKE X@N-term) | 27543 | ERp29\|PF07749.12\|6e+03 ERp29\|PF07749.12\|4.8e-25 RNA_pol_Rpb5_C\|PF01191.19\|20 RNA_po... |
| 7 | 70 | TRINITY_DN8492_c0_g1_i2.p5 | 364.63 | 77 | 110 | 10 | Oxidation (M); Amidation; Acetylation (K); Acetylation (Protein N-term); Biotinylation; 75 more | 6856 | 190(+) |
| 35 | 2 | TRINITY_DN8560_c0_g1_i1.p2 | 347.08 | 66 | 43 | 43 | Carbamidomethylation; Oxidation (M); Acetylation (N-term); Biotinylation; Carbamylation; 21 more | 46872 | TRINITY_DN8560_c0_g1_i1:4028-5233(+) |
| 41 | 312 | TRINITY_DN9_c0_g1_i1.p1 | 57.99 | 10 | 2 | 2 | Methylation; Sodium adduct; Carbamidomethylation (DHKE X@N-term); Carboxymethyl (KW X@N-term); Dihydroxy; Mutation | 8664 | TRINITY_DN9_c0_g1~~TRINITY_DN9_c0_g1_i1.p1 ORF type:complete len:78 (+) score=6.95 Secapin\|PF17521.2\|1e-34 TRINITY_DN9_c0_g1_i1:130-363(+) |
| 33 | 9 | TRINITY_DN999_c3_g1_i1.p1 | 305.58 | 69 | 46 | 43 | Carbamidomethylation; Oxidation (M); Amidation; Acetylation (K); Acetylation (N-term); 21 more | 38425 | Ribosomal_L31\|PF01197.18\|0.026 TRINITY_DN999_c3_g1_i1:593-1585(+) |
