## Supplementary Table 2 for "A common venomous ancestor? Prevalent bee venom genes evolved before the aculeate stinger while few major toxins are bee-specific"

*Halictus scabiosae*

### Transcriptomic data analysis

| Transdecoder protein IDs | Transcript ID | Expression value | Blast-P toxprot (sseqid) | E-value | Bitscore | Blast-P UniProt Apocrita (sseqid) | E-value | Bitscore | Blast-P UniProt All (sseqid) | E-value | Bitscore | Physiological association | Venom protein family |
|---|---|---|---|---|---|---|---|---|---|---|---|---|---|
| J646877.p1 | J646877 | 29.27139316 | NA | NA | NA | NA | NA | NA | Protein translation factor SUI1 like protein | 3.33E-78 | 230 | No venom function | NA |
| J647002.p1 | J647002 | 0.168754513 | NA | NA | NA | NA | NA | NA | Peptidylprolyl isomerase | 2.91E-141 | 400 | No venom function | NA |
| NODE_11040_length_424_cov_3.833811_g10395_i0.p1 | NODE_11040_length_424_cov_3.833811_g10395_i0 | 1.121949383 | NA | NA | NA | NA | NA | NA | Apolipophorins | 2.32E-47 | 169 | No venom function | NA |
| NODE_11359_length_430_cov_8.613333_g10819_i0.p1 | NODE_11359_length_430_cov_8.613333_g10819_i0 | 2.0324027 | NA | NA | NA | NA | NA | NA | Apolipophorin (Fragment) | 2.09E-56 | 195 | No venom function | NA |
| NODE_1159_length_1441_cov_15.445095_g901_i0.p1 | NODE_1159_length_1441_cov_15.445095_g901_i0 | 3.067365532 | NA | NA | NA | NA | NA | NA | Transgelin | 2.64E-130 | 370 | No venom function | NA |
| NODE_14217_length_383_cov_79.085366_g13669_i0.p2 | NODE_14217_length_383_cov_79.085366_g13669_i0 | 27.77990047 | Venom serine protease inhibitor | 8.71E-12 | 52.4 | Chymotrypsin inhibitor | 2.29E-20 | 70.9 | Venom serine protease inhibitor | 8.95E-26 | 95.9 | Venom component | Venom serine protease inhibitor |
| NODE_145_length_3358_cov_30.248553_g77_i1.p1 | NODE_145_length_3358_cov_30.248553_g77_i1 | 4.786309015 | NA | NA | NA | NA | NA | NA | Dolichyl-diphosphooligosaccharide--protein | 0 | 1411 | No venom function | NA |
| NODE_150_length_3308_cov_392.762140_g82_i0.p1 | NODE_150_length_3308_cov_392.762140_g82_i0 | 70.10773263 | NA | NA | NA | Elongation factor 1-alpha | 0.00000265 | 43.5 | Elongation factor 1-alpha | 0 | 1732 | Venom component? | EF1 alpha |
| NODE_15838_length_356_cov_2.583630_g15179_i0.p1 | NODE_15838_length_356_cov_2.583630_g15179_i0 | 1.242733601 | L-amino-acid oxidase | 4.53E-12 | 56.2 | NA | NA | NA | Smox_0 protein | 1.80E-24 | 100 | No venom function | Venom component |
| NODE_1794_length_1216_cov_13.130060_g1569_i0.p1 | NODE_1794_length_1216_cov_13.130060_g1569_i0 | 1.717444945 | NA | NA | NA | NA | NA | NA | Glyoxalase domain-containing protein 4 | 0 | 506 | No venom function | NA |
| NODE_18897_length_323_cov_3.048387_g18238_i0.p1 | NODE_18897_length_323_cov_3.048387_g18238_i0 | 2.432877846 | NA | NA | NA | NA | NA | NA | Thioredoxin-dependent peroxide mitochond | 4.34E-66 | 204 | No venom function | NA |
| NODE_1926_length_1102_cov_536.289192_g1566_i0.p1 | NODE_1926_length_1102_cov_536.289192_g1566_i0 | 98.65430645 | Translationally-controlled tumor protein ho | 6.81E-57 | 173 | NA | NA | NA | Translationally-controlled tumor protein like | 2.48E-115 | 329 | No venom function | NA |
| NODE_1966_length_1151_cov_15.059307_g1725_i0.p1 | NODE_1966_length_1151_cov_15.059307_g1725_i0 | 1.154304537 | NA | NA | NA | Major royal jelly protein 1 | 4.37E-91 | 273 | Major royal jelly protein 2 | 2.38E-113 | 341 | Venom component | Major royal jelly |
| NODE_19706_length_317_cov_5.557252_g19153_i0.p1 | NODE_19706_length_317_cov_5.557252_g19153_i0 | 1.863806317 | NA | NA | NA | NA | NA | NA | Putative polyamine oxidase 5 (Fragment) | 3.43E-37 | 135 | No venom function | NA |
| NODE_19842_length_316_cov_3.639847_g19289_i0.p1 | NODE_19842_length_316_cov_3.639847_g19289_i0 | 1.88595667 | NA | NA | NA | NA | NA | NA | Apolipophorin (Fragment) | 1.81E-30 | 119 | No venom function | NA |
| NODE_2013_length_1139_cov_12.987085_g1769_i0.p1 | NODE_2013_length_1139_cov_12.987085_g1769_i0 | 2.257870907 | Phospholipase A2 | 1.51E-40 | 132 | Phospholipase A2 | 1.60E-41 | 132 | Phospholipase A2 | 4.83E-75 | 231 | Toxin | Phospholipase A2 |
| NODE_203_length_2843_cov_39.225434_g127_i0.p1 | NODE_203_length_2843_cov_39.225434_g127_i0 | 3.085572152 | NA | NA | NA | NA | NA | NA | Cytosol aminopeptidase | 0 | 832 | No venom function | NA |
| NODE_2036_length_1134_cov_232.479147_g1790_i0.p1 | NODE_2036_length_1134_cov_232.479147_g1790_i0 | 20.85148102 | NA | NA | NA | NA | NA | NA | Guanine nucleotide-binding protein subunit | 0 | 645 | No venom function | NA |
| [illegible] | [illegible] | [illegible] | [illegible] | [illegible] | [illegible] | [illegible] | [illegible] | [illegible] | [illegible] | [illegible] | [illegible] | [illegible] | [illegible] |

### Proteome data analysis

| Checked | Protein FDR C | Master | Accession | Description | Exp. q-value: I | Coverage [%] | # Peptides | # PSMs | # Unique Pep | # AAs | MW [kDa] | calc. pI | Score Mascot | # Peptides (by Search Engine): Mascot |
|---|---|---|---|---|---|---|---|---|---|---|---|---|---|---|
| FALSE | High | Master Protein | J646877.p1 GENE.J646877~~J | PF01253.22 2e-27 J646877:688-102( | 0.003 | 35 | 3 | 4 | 3 | 110 | 12.4 | 6.77 | 52 | 3 |
| FALSE | High | Master Protein | J647002.p1 GENE.J647002~~J | PF00254.28 3.6e-36,EF-hand_1\|PF0 | 0 | 44 | 9 | 15 | 9 | 239 | 26.5 | 4.79 | 290 | 9 |
| FALSE | High | Master Protein | NODE_11040_length_424_co | PF01347.22 1.8e-12,DUF2202\|PF09 | 0 | 31 | 4 | 12 | 1 | 140 | 14.8 | 8.9 | 280 | 4 |
| FALSE | High | Master Protein | NODE_11359_length_430_co | PF04357.13 5,TamB\|PF04357.13\|1. | 0 | 30 | 4 | 7 | 4 | 143 | 15.6 | 9.09 | 154 | 4 |
| FALSE | High | Master Protein | NODE_1159_length_1441_co | PF00307.31 3.5e-23,CH\|PF00307.31 | 0 | 33 | 5 | 7 | 5 | 236 | 26.4 | 9.07 | 127 | 5 |
| FALSE | High | Master Protein | NODE_14217_length_383_co | PF01826.17 5.1e-13,cEGF\|PF12662. | 0 | 34 | 2 | 8 | 2 | 85 | 8.8 | 4.82 | 254 | 2 |
| FALSE | High | Master Protein | NODE_145_length_3358_cov_ | PF02516.14 8.8e-141,STT3\|PF0251( | 0.004 | 4 | 3 | 3 | 3 | 705 | 80.1 | 8.13 | 56 | 3 |
| FALSE | High | Master Protein | NODE_150_length_3308_cov_ | PF00009.27 2e-66,EFG_IV\|PF03764 | 0 | 15 | 10 | 16 | 10 | 844 | 94.6 | 6.57 | 150 | 10 |
| FALSE | High | Master Protein | NODE_15838_length_356_co | PF01593.24 5.2e-17,NAD_binding_8 | 0 | 37 | 2 | 4 | 2 | 90 | 9.6 | 4.78 | 137 | 2 |
| FALSE | High | Master Protein | NODE_1794_length_1216_co | PF00903.25 0.0016,Glyoxalase\|PF0( | 0 | 15 | 4 | 7 | 4 | 281 | 32 | 5.76 | 71 | 4 |
| FALSE | High | Master Protein | NODE_18897_length_323_co | PF00578.21 6.9e-33,Redoxin\|PF085 | 0.004 | 39 | 3 | 3 | 3 | 107 | 12 | 7.33 | 57 | 3 |
| FALSE | High | Master Protein | NODE_1926_length_1102_co | PF00838.17 1.2e-62,PfUIS3\|PF1156 | 0 | 31 | 4 | 6 | 4 | 172 | 19.8 | 4.81 | 104 | 4 |
| FALSE | High | Master Protein | NODE_1966_length_1151_co | PF03022.16 5.9e-64,NHL\|PF01436.2 | 0 | 34 | 9 | 16 | 9 | 353 | 38.9 | 5.41 | 222 | 9 |
| FALSE | High | Master Protein | NODE_19706_length_317_co | PF01593.24 4.1e-17,Phage_tail_2\|P | 0.004 | 51 | 4 | 6 | 4 | 92 | 10.2 | 6.79 | 62 | 4 |
| FALSE | High | Master Protein | NODE_19842_length_316_co | GENE.NODE_19842_length_316_cov_3 | 0 | 38 | 4 | 7 | 4 | 105 | 11.6 | 8.97 | 183 | 4 |
| FALSE | High | Master Protein | NODE_2013_length_1139_co | PF05826.12 2.8e-37,Phospholip_A2_ | 0 | 33 | 6 | 8 | 6 | 216 | 24.8 | 5.83 | 88 | 6 |
| FALSE | High | Master Protein | NODE_203_length_2843_cov_ | PF00883.21 2.7e+03,Peptidase_M17 | 0 | 35 | 13 | 25 | 13 | 509 | 54.7 | 7.12 | 241 | 13 |
| FALSE | High | Master Protein | NODE_2036_length_1134_co | PF00400.32 0.00041,WD40\|PF0040( | 0.004 | 30 | 7 | 8 | 7 | 317 | 35.6 | 7.74 | 61 | 7 |
| [illegible] | [illegible] | [illegible] | [illegible] | [illegible] | [illegible] | [illegible] | [illegible] | [illegible] | [illegible] | [illegible] | [illegible] | [illegible] | [illegible] | [illegible] |
