## Supplementary Table 3 for "A common venomous ancestor? Prevalent bee venom genes evolved before the aculeate stinger while few major toxins are bee-specific"

*Xylocopa violacea*

| Transdecoder protein IDs | Transcript ID | Expression value | Blast-P toxprot (sseqid) | E-value | Bitscore | Blast-P UniProt Apocrita (sseqid) | E-value | Bitscore | Blast-P UniProt All (sseqid) | E-value | Bitscore | Physiological association | Venom protein family | Protein Group | Protein ID | Accession | -10lgP | Coverage (%) | #Peptides | #Unique | PTM | Avg. Mass | Description |
|---|---|---|---|---|---|---|---|---|---|---|---|---|---|---|---|---|---|---|---|---|---|---|---|
| | | | | | Transcriptome data analysis | | | | | | | | | Proteome data analysis | | | | | | | | | |
| J678067.p1 | J678067 | 4.49562464 | NA | NA | NA | NA | NA | NA | Uncharacterized protein/putative sodium-c[illegible] | 4.47E-154 | 452 | No venom function | NA | 94 | 67 | J678067.p1 | 122.12 | 14 | 4 | 4 | | 51609 | GENE.J678067~~J678067.p1 ORF type:complete len:514 (+) score=119.51 J678067:205-1542(+) |
| NODE_10042_length_1345_cov_20691.368992_g138_i17.p2 | NODE_10042_length_1345_cov_20691.368992_g138_i17 | 5082.365988 | Apamin | 0.00000833 | 34.7 | Apamin | 8.83E-07 | 34.7 | NA | NA | NA | Toxin | Apamin | 40 | 108 | NODE_10042_length_1 | 131.96 | 59 | 9 | 9 | Carbamidom | 5599 | GENE.NODE_10042_length_1345_cov_20691.368992_g138_i17~~NODE_10042_length_1345_cov_20691.368992_g138_i17.p[illegible] |
| NODE_1035_length_5669_cov_154.173579_g758_i0.p2 | NODE_1035_length_5669_cov_154.173579_g758_i0 | 40.89185579 | NA | NA | NA | NA | NA | NA | Moesin/ezrin/radixin 1 | 4.49E-148 | 429 | No venom function | NA | 177 | 218 | NODE_1035_length_56 | 65.85 | 10 | 2 | 2 | | 27038 | GENE.NODE_1035_length_5669_cov_154.173579_g758_i0~~NODE_1035_length_5669_cov_154.173579_g758_i0.p2 ORF [illegible] |
| NODE_10529_length_1115_cov_107.359615_g7997_i0.p1 | NODE_10529_length_1115_cov_107.359615_g7997_i0 | 57.37424136 | NA | NA | NA | Histone H4 | 1.44E-70 | 199 | Histone H4 (Fragment) | 9.39E-67 | 201 | No venom function | NA | 73 | 84 | NODE_10529_length_1 | 104.87 | 47 | 5 | 5 | Oxidation (M | 11500 | GENE.NODE_10529_length_1115_cov_107.359615_g7997_i0~~NODE_10529_length_1115_cov_107.359615_g7997_i0.p1 [illegible] |
| NODE_11081_length_1183_cov_321.804965_g7134_i1.p1 | NODE_11081_length_1183_cov_321.804965_g7134_i1 | 0 | NA | NA | NA | NA | NA | NA | Superoxide dismutase [Cu-Zn], chloroplastic | 2.53E-88 | 287 | Venom component (parasitoids) | Superoxide dismutase | 110 | 179 | NODE_11081_length_1 | 109.62 | 21 | 2 | 2 | Carbamidom | 18270 | GENE.NODE_11081_length_1183_cov_321.804965_g7134_i1~~NODE_11081_length_1183_cov_321.804965_g7134_i1.p1 [illegible] |
| NODE_11225_length_1162_cov_51432.834688_g2195_i1.p4 | NODE_11225_length_1162_cov_51432.834688_g2195_i1 | 13.00180871 | Apamin | 0.00000146 | 36.6 | Apamin | 1.55E-07 | 36.6 | NA | NA | NA | Toxin | Apamin | 13 | 115 | NODE_11225_length_1 | 197.87 | 56 | 23 | 2 | Carbamidom | 6237 | GENE.NODE_11225_length_1162_cov_51432.834688_g2195_i1~~NODE_11225_length_1162_cov_51432.834688_g2195_i1.p[illegible] |
| NODE_11494_length_1125_cov_206.560748_g8269_i0.p1 | NODE_11494_length_1125_cov_206.560748_g8269_i0 | 98.97275408 | NA | NA | NA | NA | NA | NA | Peptidyl-prolyl cis-trans isomerase | 1.44E-141 | 398 | No venom function | NA | 72 | 87 | NODE_11494_length_1 | 105.47 | 24 | 5 | 5 | Carbamidom | 22861 | GENE.NODE_11494_length_1125_cov_206.560748_g8269_i0~~NODE_11494_length_1125_cov_206.560748_g8269_i0.p1 [illegible] |
| NODE_11707_length_957_cov_56.019274_g8998_i0.p1 | NODE_11707_length_957_cov_56.019274_g8998_i0 | 9.231161332 | NA | NA | NA | NA | NA | NA | Sep15_SelM domain-containing protein | 5.08E-95 | 276 | No venom function | NA | 68 | 75 | NODE_11707_length_9 | 109.2 | 30 | 5 | 5 | Carbamidom | 18018 | GENE.NODE_11707_length_957_cov_56.019274_g8998_i0~~NODE_11707_length_957_cov_56.019274_g8998_i0.p1 ORF [illegible] |
| NODE_12215_length_900_cov_350.141818_g8219_i1.p1 | NODE_12215_length_900_cov_350.141818_g8219_i1 | 213.5472267 | NA | NA | NA | NA | NA | NA | Superoxide dismutase [Cu-Zn], chloroplastic | 2.53E-88 | 287 | Venom component (parasitoids) | Superoxide dismutase | 110 | 178 | NODE_12215_length_9 | 109.62 | 21 | 2 | 2 | Carbamidom | 18270 | GENE.NODE_12215_length_900_cov_350.141818_g8219_i1~~NODE_12215_length_900_cov_350.141818_g8219_i1.p1 ORF [illegible] |
| NODE_12300_length_891_cov_419.590686_g9496_i0.p1 | NODE_12300_length_891_cov_419.590686_g9496_i0 | 21.45689413 | NA | NA | NA | NA | NA | NA | Superoxide dismutase [Cu-Zn] | 2.37E-93 | 272 | Venom component (parasitoids) | Superoxide dismutase | 49 | 42 | NODE_12300_length_8 | 150.66 | 61 | 8 | 8 | Carbamidom | 15698 | GENE.NODE_12300_length_891_cov_419.590686_g9496_i0~~NODE_12300_length_891_cov_419.590686_g9496_i0.p1 ORF [illegible] |
| NODE_12392_length_882_cov_458.973978_g9580_i0.p1 | NODE_12392_length_882_cov_458.973978_g9580_i0 | 291.9235864 | NA | NA | NA | NA | NA | NA | Transgelin | 5.4E-130 | 367 | No venom function | NA | 81 | 89 | NODE_12392_length_8 | 112.78 | 29 | 4 | 4 | Carbamidom | 20570 | GENE.NODE_12392_length_882_cov_458.973978_g9580_i0~~NODE_12392_length_882_cov_458.973978_g9580_i0.p1 ORF [illegible] |
| NODE_12433_length_1006_cov_20793.932702_g1692_i3.p4 | NODE_12433_length_1006_cov_20793.932702_g1692_i3 | 2490.904444 | NA | NA | NA | NA | NA | NA | Uncharacterized/Protein CBR-GRL-22 | 1.20E+00 | 75 | Unknown function | NA | 2 | 241 | NODE_12433_length_1 | 254.68 | 63 | 24 | 21 | Carbamidom | 5986 | GENE.NODE_12433_length_1006_cov_20793.932702_g1692_i3~~NODE_12433_length_1006_cov_20793.932702_g1692_i3.p[illegible] |
| NODE_1293_length_5449_cov_108.784946_g950_i0.p1 | NODE_1293_length_5449_cov_108.784946_g950_i0 | 37.9258919 | NA | NA | NA | NA | NA | NA | Carboxypeptidase D | 0 | 2285 | Venom component | CarboxypeptidaseD | 133 | 322 | NODE_1293_length_54 | 38.7 | 1 | 2 | 2 | | 189105 | GENE.NODE_1293_length_5449_cov_108.784946_g950_i0~~NODE_1293_length_5449_cov_108.784946_g950_i0.p1 ORF [illegible] |
| NODE_1388_length_4997_cov_135.900244_g975_i1.p1 | NODE_1388_length_4997_cov_135.900244_g975_i1 | 53.08652634 | NA | NA | NA | Arginine kinase | 0 | 672 | Arginine kinase | 0 | 716 | Venom component | Arginine kinase | 46 | 18 | NODE_1388_length_49 | 159.52 | 30 | 10 | 10 | Carbamidom | 39904 | GENE.NODE_1388_length_4997_cov_135.900244_g975_i1~~NODE_1388_length_4997_cov_135.900244_g975_i1.p1 ORF [illegible] |
| [illegible] | [illegible] | [illegible] | [illegible] | [illegible] | [illegible] | [illegible] | [illegible] | [illegible] | [illegible] | [illegible] | [illegible] | [illegible] | [illegible] | [illegible] | [illegible] | [illegible] | [illegible] | [illegible] | [illegible] | [illegible] | [illegible] | [illegible] | [illegible] |
