## Supplementary Table 4 for "A common venomous ancestor? Prevalent bee venom genes evolved before the aculeate stinger while few major toxins are bee-specific"

| Venom component | Function | Activity | Reference |
|---|---|---|---|
| Apamin | Toxin | Neurotoxic | Apamin blocks certain neurotransmitter-induced increases in potassium permeability. B. E. C. Banks, C. Brown, G. M. Burgess, G. Burnstock, M. Claret, T. M. Cocks & D. H. Jenkinson. Nature volume 282, pages 415–417 (1979)<br>Apamin, a neurotoxin specific for one class of Ca2+-dependent K+ channels. M Lazdunski.Cell Calcium. 1983. Dec;4(5-6):421-8. |
| Mast cell degranulating protein | Venom component | Neurotoxic, Histamin release | Handbook of Biologically Active Peptides (Second Edition), Chapter 58 - Hymenoptera Insect Peptides, 2013, Pages 416-422, Mario SergioPalma<br>xPharm: The Comprehensive Pharmacology Reference, 2007, Pages 1-3, Mast Cell Degranulating Peptide, Mark A.Simmons |
| Melittin | Toxin | Cytolytic, disrupts surface tension, relea | Biosci Rep (2007) 27:189–223 DOI 10.1007/s10540-006-9030-z, REVIEW ARTICLE, Melittin: a Membrane-active Peptide with Diverse Functions, H. Raghuraman Æ Amitabha Chattopadhyay<br>Review, Bee Venom: An Updating Review of Its Bioactive Molecules and Its Health Applications, Maria Carpena , Bernabe Nuñez-Estevez , Anton Soria-Lopez and Jesus Simal-Gandara, Nutrients, MDPI, 2020 |
| Secapin | Venom component | Antifibronilytic | Secapin, a bee venom peptide, exhibits anti-fibrinolytic, anti-elastolytic, and anti-microbial activities. Kwang SikLeea1Bo YeonKima1Hyung JooYoonbYong SooChoibByung RaeJina. Developmental & Comparative Immunology. Volume 63, October 2016, Pages 27-35 |
| Hyaluronidase | Toxin | Spreading factor, pore forming | Marković-Housley Z., Miglierini G., Soldatova L., Rizkallah P.J., Müller U., Schirmer T. Crystal structure of hyaluronidase, a major allergen of bee venom. Structure. 2000;8:1025–1035. doi: 10.1016/S0969-2126(00)00511-6. [PubMed] [CrossRef] [Google Scholar] [Ref list]<br>Diversity of peptidic and proteinaceous toxins from social Hymenoptera venoms. Dos Santos-Pinto JRA, Perez-Riverol A, Lasa AM, Palma MS. Toxicon. 2018 Jun 15; 148():172-196.<br>Bee Venom: An Updating Review of Its Bioactive Molecules and Its Health Applications. Maria Carpena, Bernabe Nuñez-Estevez, Anton Soria-Lopez, and Jesus Simal-Gandara. Nutrients. 2020 Nov; 12(11): 3360. Published online 2020 Oct 31. doi: 10.3390/nu12113360. PMCID: PMC7693387. PMID: 33142794 |
| Icarapin | Venom component | Undefined | NA |
| Phospholipase A2 | Toxin | Neurotoxic, | Localization of structural elements of bee venom phospholipase A2 involved in N-type receptor binding and neurotoxicity. J P Nicolas 1 , Y Lin, G Lambeau, F Ghomashchi, M Lazdunski, M H Gelb. J Biol Chem. 1997 Mar 14;272(11):7173-81. doi: 0.1074/jbc.272.11.7173.<br>Review, Bee Venom: An Updating Review of Its Bioactive Molecules and Its Health Applications, Maria Carpena , Bernabe Nuñez-Estevez , Anton Soria-Lopez and Jesus Simal-Gandara, Nutrients, MDPI, 2020 |
| Venom antigen 3/5 | Venom component | Undefined | NA |
| Venom dipeptidyl peptidase 4 | Venom component | Processses venom components e.g. me | Bee Venom: An Updating Review of Its Bioactive Molecules and Its Health Applications. Maria Carpena, Bernabe Nuñez-Estevez, Anton Soria-Lopez, and Jesus Simal-Gandara. Nutrients. 2020 Nov; 12(11): 3360. Published online 2020 Oct 31. doi: 10.3390/nu12113360. PMCID: PMC7693387. PMID: 33142794 |
| Venom acid phosphatase | Venom component | Histamin release | Bee Venom: An Updating Review of Its Bioactive Molecules and Its Health Applications. Maria Carpena, Bernabe Nuñez-Estevez, Anton Soria-Lopez, and Jesus Simal-Gandara. Nutrients. 2020 Nov; 12(11): 3360. Published online 2020 Oct 31. doi: 10.3390/nu12113360. PMCID: PMC7693387. PMID: 33142794 |
| Venom serine protease | Venom component | Lethal Melanization repsonse, | Dual Function of a Bee Venom Serine Protease: Prophenoloxidase-Activating Factor in Arthropods and Fibrin(ogen)olytic Enzyme in Mammals. Young Moo Choo1., Kwang Sik Lee1., Hyung Joo Yoon2, Bo Yeon Kim1, Mi Ri Sohn1, Jong Yul Roh3, Yeon Ho Je3, Nam Jung Kim2, Iksoo Kim4, Soo Dong Woo5, Hung Dae Sohn1, Byung Rae Jin1. PLOS one 2010. 5 (5) |
| Tertiapin | Toxin | Neurotoxic | *Gauldie, J; Hanson, JM; Rumjanek, FD; Shipolini, RA; Vernon, CA (1976). "The peptide components of bee venom". European Journal of Biochemistry.* ***61*** *(2): 369–376. doi:10.1111/j.1432-1033.1976.tb10030.x. PMID 1248464.*<br>The Effect of Bee Venom Peptides Melittin, Tertiapin, and Apamin on the Human Erythrocytes Ghosts: A Preliminary Study. Świątły-Błaszkiewicz, A.; Mrówczyńska, L.; Matuszewska, E.; Lubawy, J.; Urbański, A.; Kokot, Z.J.; Rosiński, G.; Matysiak, J. *Metabolites* **2020**, *10* , 191. https://doi.org/10.3390/metabo10050191 |
