## Supplementary Table 6 for "A common venomous ancestor? Prevalent bee venom genes evolved before the aculeate stinger while few major toxins are bee-specific"

**Supplementary Table 5: Overview of the newly-generated transcriptome and proteome datasets for the venom glands of *A. mellilfera*, *H. scabiosae* and *X. violacea*.** Relatively low BUSCO value for *H. scabiosae* transcriptome could be explained by the fact that of the three species its glands are by far the smallest and thus gland-specific transcripts will have higher abundance compared to housekeeping transcripts, as well as computed score being lower than actual due to BUSCO methodology inflating the proportion of missing genes as elaborated by Simão et al (Simão et al. 2015). Due to the protection status of bees and subsequent nature conservation regulations we were limited in the number of specimens and could not dramatically increase the sample size to increase tissue quantities.

|  | ***A. mellifera*** | ***H. scabiosae*** | ***X. violacea*** | |
| --- | --- | --- | --- | --- |
| Illumina reads (paired end, 150 bp) | 26,092,431 | 38,605,338 | 16,077,242 | |
| Assembled transcripts | 31,282 | 80,388 | | 63,006 |
| BUSCO values | 97.7% | 41.6% | 97.4% | |
| Predicted peptides | 74,609 | 94,492 | 144,998 | |
| MS-supported protein matches | 121 | 211 | 187 | |
